# Robust error-minimization in the genetic code across physicochemical metrics and variant codes: a graph-theoretic analysis in GF(2)^6^

**DOI:** 10.64898/2026.04.25.720843

**Authors:** Paul Clayworth, Sergey Kornilov

## Abstract

The standard genetic code reduces the impact of point mutations, but the robustness of this property across physicochemical metrics, naturally occurring variant codes, and codon-reassignment mechanisms remains incompletely quantified. Embedding the 64 codons in GF(2)^6^ represents the hypercube *Q*_6_ as a coordinate-dependent subgraph of the encoding-independent single-nucleotide mutation graph *H*(3, 4), and enables continuous *ρ*-interpolation between the two. Under a quartet-pattern shuffle null (*n* =10,000), the standard code is significantly low-cost across four established, code-independent physicochemical distance metrics with partially overlapping content (Grantham *p* = 0.0062; Miyata *p* < 0.001; Woese polar requirement *p* = 0.003; Kyte–Doolittle hydropathy *p* = 0.001), and the signal strengthens monotonically as *ρ* moves *Q*_6_ → *H*(3, 4). A structure-aware sensitivity analysis under the alignment-derived ProtSub matrix (Jia & Jernigan 2021) yields the most extreme percentile of any measure tested (*p* = 0.0004; all five p-values pass Bonferroni at *α* = 0.05). Across the 27 NCBI translation tables, near-optimality is preserved: 11 of 12 informative-distance variants retain top-5% placement after BH–FDR correction. Natural codon reassignments avoid disrupting codon-family connectivity: under the encoding-independent *H*(3, 4) adjacency, observed events are topology-breaking at relative risk 0.32 versus the candidate landscape (permutation *p* ≤ 10^−4^). The *H*(3, 4) result is stable by construction; the *Q*_6_ decomposition is representation-specific and fails to show depletion under 8 of 24 base-to-bit encodings, so we report *H*(3, 4) as the primary test and *Q*_6_ as a sensitivity. Event-level conditional-logit modelling shows that topology avoidance and local physicochemical cost provide complementary, only weakly correlated signal (*r_s_* = 0.15), and that topology adds explanatory value beyond physicochemistry under both *Q*_6_ and encoding-independent *H*(3, 4) adjacency. Retrospective reanalysis of nine genome-recoding datasets is consistent with codon-family topology operating as an evolutionary-trajectory constraint distinct from acute engineering fitness. The contribution is the second axis: code evolution is jointly constrained by physicochemical smoothness and codon-family topological integrity, and these two constraints are partly independent.

**Highlights:**

- Codon-space geometry links genetic-code robustness and reassignment paths
- Standard and variant codes preserve broad physicochemical error minimization
- Reassignments are depleted for codon-family topology-breaking moves
- Conditional-logit models separate topology from physicochemical similarity
- Synthetic recoding shows boundary conditions for natural-code constraints

## 1 Introduction

The standard genetic code maps 61 sense codons to 20 amino acids through a pattern that has long been recognized as non-random. Woese (1965) first noted that similar codons tend to encode amino acids with similar physicochemical properties, and Freeland and Hurst (1998) demonstrated quantitatively that the standard code sits among approximately 1 in 10^6^ random codes for mutational error minimization under the Woese polar requirement distance. Follow-up analyses refined this claim: Novozhilov et al. (2007) and Novozhilov and Koonin (2009) characterised how the standard code compares against rugged-fitness-landscape and primordial-code null ensembles; alternative selection accounts frame the code as the outcome of gene–code coevolution (Sella and Ardell, 2006) or collective evolution with horizontal information exchange (Vetsigian et al., 2006) a review of the competing origin-of-the-code frameworks appears in this journal in Di Giulio (2005). These findings established that the code’s structure reduces the fitness impact of point mutations, but whether this optimality is specific to a single metric or robust across distinct physicochemical parameterizations has not been systematically tested.

Every codon comprises three nucleotides drawn from {*C, U, A, G*}. By choosing a bijection *φ*: {*C, U, A, G*} → GF(2)^2^ (for instance, *C* ↦ (0, 0), *U* ↦ (0, 1), *A* ↦ (1, 0), *G* ↦ (1, 1)), each codon maps to a vertex of the 6-dimensional binary hypercube *Q*_6_ = GF(2)^6^. The genetic code then becomes a *coloring* of *Q*_6_ by 21 labels (20 amino acids plus the stop signal), and single-nucleotide mutations correspond to edges of *Q*_6_ or, more precisely, to a subgraph of the complete mutation graph *H*(3, 4).

This representation is not new in principle: binary encodings of the genetic code appear in the mathematical biology literature (e.g., (Antoneli and Forger, 2011)). Its value lies not in the encoding itself but in the analytical decomposition it enables: the complete single-nucleotide mutation graph *H*(3, 4) (288 edges) separates into 192 Hamming-distance-1 edges (single-bit changes) and 96 within-nucleotide distance-2 edges (both bits of one nucleotide position flip simultaneously), allowing systematic interpolation via a weight parameter *ρ* ∈ [0, 1]. This decomposition does not correspond to the biological transition/transversion partition: under 16 of the 24 possible 2-bit encodings, the Hamming-1 edges contain an equal mixture of transitions and transversions per nucleotide position (2 of each among 4 Hamming-1 pairs), while the remaining 8 encodings place both transitions on diagonal (Hamming-2) edges. The *ρ* parameter should therefore be interpreted as a diagonal-edge inclusion weight, not a transition/transversion weight. Previous work has not exploited this decomposition to test error-minimization across multiple physicochemical metrics, nor extended the analysis to variant genetic codes. Doing so lets us examine two questions that single-axis analyses cannot. First, whether error-minimization is a property of the code itself rather than of any particular distance function. Because the established physicochemical measures (composition–polarity–volume, polarity–volume, chromatographic polar requirement, hydropathy) are partially overlapping, we do not treat concordance across them as independent replication, but as a convergent sensitivity envelope: if the standard code lies in the low-cost tail under several established but non-identical parameterizations, the result is not confined to a single chosen scale (overlap quantified via pairwise and partial Spearman correlations in §3.1). Second, whether codon-family *connectivity* acts as a second, partly independent axis of constraint on natural reassignment events, distinct from physicochemical cost. If so, code evolution is bounded not only by which codes are low-cost but by which trajectories through code-space leave the decoding substrate intact, a constraint on transitions rather than on states. These two motivating questions are operationalized as three concrete tests:

1. **Is the standard code optimal under multiple physicochemical metrics?** We test whether the standard code’s edge-mismatch score is extreme relative to quartet-pattern shuffle null models across four established, code-independent physicochemical distance measures with partially overlapping content (Grantham, Miyata, Woese polar requirement, and Kyte–Doolittle hydropathy), extending Freeland and Hurst (1998) from a single metric to a cross-metric sensitivity envelope.
2. **Is this structure preserved by evolution?** We ask whether variant genetic codes (NCBI translation tables 2–33) maintain error-minimization, and whether natural codon reassignment events preferentially avoid disrupting the topological connectivity of amino acid codon families.
3. **What are the genomic correlates of disruption?** We test whether organisms whose variant codes break the connectivity of an amino acid’s codon graph show elevated tRNA gene copy numbers for the affected amino acid, using tRNAscan-SE–verified data from 18 genomes spanning 5 variant genetic codes across Alveolata, Opisthokonta, Excavata, and Mollicutes; 15 of these 18 genomes populate the 24-pairing enrichment analysis, with the remaining 3 (Blastocrithidia and two Mycoplasmoides species) discussed as mechanistic boundary cases in §4.3 because their reassignment routes act without changing tRNA gene counts.

Each of these three tests carries an explicit decision criterion. For (1), a null result would be quartet-pattern shuffie null *p* > 0.05 on any of the four physicochemical metrics; discordance among metrics (some significant, some not) would restrict any claim of optimality to a specific parameterization rather than to the code itself. For (2), a null result would be either failure of BH-corrected per-table significance in the majority of informative-distance tables (*d_H_* ≥ 3 reassignments from the standard code), or an observed topology-breaking rate not distinguishable from the candidate-landscape rate (hypergeometric *p* > 0.05 or risk-ratio 95% CI including 1.0). For (3), the originally specified decision gate was the **worst-case** Stouffer *p*-value across all maximal independent pairing subsets; that gate was not met (*p* = 0.104; Section 3.6), so the tRNA result is reported as **exploratory** rather than as confirmatory support for the compensation-via-tRNA-duplication hypothesis. We report the median and full MIS distribution descriptively (median *p* = 0.037; 264 of 332 MIS below 0.05); these motivate an exploratory decoding-accommodation signal rather than a confirmatory enrichment claim. The pre-specified topology-breaking-only subset (*n* = 4) is examined as a complementary mechanistic test and is also null (Stouffer *p* = 0.387). We report all three tests with the corresponding numbers regardless of outcome.

To delimit the framework’s scope, we also test four conjectural extensions previously associated with this project (one of us, P.C.; (Clayworth, 2026)): a Serine distance-4 invariance claim, PSL(2,7) symmetry, a holomorphic embedding extending a character of GF(8)^∗^, and a KRAS–Fano clinical prediction. These tests distinguish supported graph-theoretic results from encoding artifacts and unsupported algebraic extensions (Section 3.9; KRAS–Fano detail in Supplement §S13).

The paper is organized as follows. Section 2 describes the encoding formalism, graph decomposition, null models, and statistical methods, including a retrospective cross-study reanalysis of nine published genome-recoding datasets (eight analyzed quantitatively). Section 3 presents the four supported findings (cross-metric coloring optimality, per-table preservation, *ρ*-robustness, and topology avoidance), the exploratory tRNA enrichment result, the synthetic-recoding reanalysis, and additional exploratory observations. Section 4 discusses the graph-theoretic interpretation, the relationship to frozen-accident versus adaptive hypotheses (Koonin and Novozhilov, 2009), and an exploratory three-layer interpretation. All analyses are reproducible via the open-source `codon-topo` pipeline (version 0.6.1, seed 135325).

## 2 Methods

### 2.1 Binary encoding of the genetic code

We encode each nucleotide base as a 2-bit vector via the default bijection *φ*: *C* ↦ (0, 0), *U* ↦ (0, 1), *A* ↦ (1, 0), *G* ↦ (1, 1). A codon *b*_1_*b*_2_*b*_3_ is then represented as the concatenation *φ*(*b*_1_) ‖ *φ*(*b*_2_) ‖ *φ*(*b*_3_) ∈ GF(2)^6^, and the 64 codons become the 64 vertices of the 6-dimensional hypercube *Q*_6_. This default bijection was adopted in companion methodological work (Clayworth (2026)) because it places the standard code’s nine 2-fold-degenerate amino acids on bit-5 differences. The visualization-clarity rationale, however, is mildly circular: the bit-5 two-fold filtration is itself classified as Tautological in the present claim hierarchy (Supplement §S1), so the encoding was chosen to make a tautological property visible. We therefore use the default encoding only as an analytical convenience and make no claim that it is biologically privileged. Two specific encoding-dependent results are reported under all 24 base-to-bit bijections: (i) the coloring-optimality Grantham quartet-pattern shuffle null (Supplement §S2), and (ii) the *Q*_6_ topology-avoidance landscape depletion (Supplement §S4). The primary topology-avoidance test uses the encoding-independent *H*(3, 4) adjacency (§2.3.4). The conditional-logit models M1–M4 are not swept across encodings; their *H*(3, 4) verification variants replace the Δ_topo,Q_6_ topology feature with the encoding-independent Δ_topo,H(3,4)_feature but retain a local physicochemical feature evaluated on *Q*_6_ Hamming-1 edges, so those variants are best described as **topology-verification** rather than as fully encoding-independent combined models.

Under this encoding, two codons that differ by a single-nucleotide substitution in which only one bit of the 2-bit pair changes are adjacent in *Q*_6_ (Hamming distance 1). However, transversions that flip both bits within a nucleotide position correspond to Hamming distance 2. The full single-nucleotide mutation graph is therefore the Hamming graph *H*(3, 4) = *K*_4_ ▫ *K*_4_ ▫ *K*_4_ (the Cartesian product of three complete graphs on the four nucleotide states; equivalently, *K*^▫3^), with 64 vertices, regular degree 9, and 288 undirected edges. It contains *Q*_6_ as a 192-edge subgraph: *Q*_6_ contributes the 192 Hamming-distance-1 edges (single-bit changes within one 2-bit nucleotide), while the remaining 96 within-nucleotide diagonal (Hamming-distance-2) edges complete *H*(3, 4). To address the concern that *Q*_6_ misses approximately one-third of single-nucleotide mutations, we introduce the weighted score *F_ρ_* that interpolates between pure *Q*_6_ (*ρ* = 0) and full *H*(3, 4) (*ρ* = 1); see Section 2.3. Throughout this paper *H*(3, 4) denotes this nucleotide-level mutation graph, which is encoding-independent (every two-bit bijection from {*A, C, G, U*} to {0, 1}^2^ yields the same *H*(3, 4)); *Q*_6_ is encoding-dependent, since the partition into Hamming-1 vs Hamming-2 edges depends on the bijection.

There are 4! = 24 such bijections. The two encoding-dependent analyses in this work (cross-metric coloring optimality and the *Q*_6_ topology-avoidance depletion) are evaluated under all 24 bijections; the M1–M4 conditional-logit models and the primary *H*(3, 4) topology-avoidance test are encoding-invariant by construction and are not swept. Encoding-invariant properties (such as Serine’s disconnection at *ε* = 1, where *ε* denotes the Hamming distance threshold in GF(2)^6^) are noted as such. Coloring optimality is significant (*p* < 0.05) under every encoding (full sweep in the Supplementary Material).

### 2.2 Edge-mismatch objective function

The genetic code assigns each vertex *v* ∈ *Q*_6_ a label *c*(*v*) ∈ *A*, where *A* comprises the 20 amino acids and the stop signal. The edge-mismatch score is

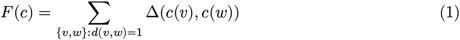

where the sum ranges over all 192 edges of *Q*_6_ (pairs of vertices at Hamming distance 1), and Δ is a physicochemical distance between amino acids. We test four established, code-independent physicochemical distance measures with partially overlapping content: the Grantham (1974) composite distance (composition, polarity, volume; range 5–215), the Miyata et al. (1979) normalized Euclidean distance (polarity and volume only; range 0.06–5.13), the Woese polar requirement absolute difference (range 0–8.2; (Woese, 1973, Woese et al., 1966); used by Freeland and Hurst (1998)), and the Kyte and Doolittle (1982) hydropathy absolute difference (range 0–9.0; used by Haig and Hurst (1991)). All four are **code-independent** physicochemical scales, derived from properties of amino acids in isolation (chromatographic partition coefficient, composition, polarity, volume, hydropathy) rather than from observed substitution patterns in coded proteins; Kosiol et al. (2004) review criteria for principled amino-acid classification schemes in this same code-independent regime. This design avoids the tautology that Di Giulio (2001) identified in PAM-derived measurements of code optimality, where the substitution matrix itself reflects the genetic code structure under study; Koonin and Novozhilov (2009) explicitly endorse code-independent physicochemical scales for exactly this reason. As a structure-aware sensitivity analysis, we also report results under the MSA-derived ProtSub matrix (Jia and Jernigan (2021) 2,320 Pfam domains), which is alignment-derived and therefore code-dependent in the sense of Di Giulio (2001). ProtSub is treated as a robustness check rather than as primary evidence for code-origin optimality. Synonymous edges contribute 0; edges involving a stop codon receive a fixed penalty scaled proportionally to each metric’s maximum. A lower *F* indicates a more error-minimizing code.

Stop codons are held fixed across all null models (Section 2.3) because their assignment is constrained by release-factor recognition geometry (eRF1 binding UAR/UGA) rather than tRNA decoding, and permuting stops would conflate two evolutionarily separate optimization problems. The stop-codon contribution is therefore a constant offset; sensitivity analysis across penalty values (0, 150, 215, 300) confirms this is immaterial to the ranking.

### 2.3 Null models

#### 2.3.1 Quartet-pattern shuffle null

The randomisation ensemble used throughout the coloring-optimality tests is a **quartet-pattern shuffle**: we group the 64 codons into 16 blocks of 4, defined by shared first-two-base prefix (the codon “quartets”). Each quartet’s internal 4-tuple of amino-acid labels is held as an atomic pattern, and the 14 non-stop quartet patterns are permuted uniformly at random among the 14 non-stop quartet slots (the 2 stop-containing quartets are held fixed). The null preserves each quartet’s internal four-position amino-acid split shape (whether it is a (4), (2, 2), (3, 1), or singleton block) and each named amino acid’s total codon count, while permuting **which** codons carry those labels; no quartet is broken up, but amino-acid labels move between quartet slots.

The reassignment biology literature also uses a related null often cited to Haig and Hurst (1991) and Freeland and Hurst (1998): it fixes the standard code’s 20 sense-codon families (the unlabeled partition of the 61 sense codons into synonymous groups, holding stop positions fixed) and permutes the 20 amino-acid labels uniformly across those families. This classical Haig–Hurst / Freeland–Hurst amino-acid-permutation null is a different randomisation ensemble from the quartet-pattern shuffle we implement. The two ensembles preserve non-nested structural properties: the quartet-pattern shuffle moves the standard code’s atomic 4-codon quartet patterns among the 14 non-stop quartet slots, so each named amino acid’s total codon count is preserved but which specific codons decode as that amino acid is not; the AA-permutation null fixes the codon-family partition but allows a named amino acid’s codon count to change (e.g., Trp’s single-codon family may become a 6-codon family if its label is swapped with Leu’s). The two ensembles admit different numbers of alternative codes (≈ 14! ≈ 8.7 × 10^10^ for the quartet-pattern shuffle versus ≈ 20! ≈ 2.4 × 10^18^ for the AA-permutation null) and can produce different null tails. We report the four physicochemical metrics under **both** nulls in Supplement §S2.1: treating the eight reported *p*-values (four metrics × two nulls, *n* =10,000 draws each, seed 135325) as one test family, all eight pass Bonferroni at *α* = 0.05/8 = 0.00625. For three of the four code-independent metrics the classical Haig–Hurst null gives a **more** extreme *p* than the quartet-pattern shuffle, and for Kyte–Doolittle the direction reverses. We therefore do not describe either null as uniformly “more stringent” and treat the classical null as a same-direction sensitivity check on the quartet-pattern-shuffle headline values.

For the standard code, we drew *n* =10,000 null samples (seed 135325) from the quartet-pattern shuffle and computed the conservative *p*-value as 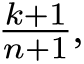 where *k* is the number of null scores satisfying *F*_null_ ≤ *F*_obs_; ties at the observed score are included in *k*.

#### 2.3.2 Per-table null

The NCBI Genetic Codes registry (https://www.ncbi.nlm.nih.gov/Taxonomy/Utils/wprintgc.cgi; gc.prt v4.6, retrieved 2026-04-25) currently lists 27 translation tables: codes 1–6, 9–16, and 21–33 (codes 7, 8, and 17–20 are deprecated and absent from the current registry). All 27 are analyzed in this work. Two pairs of tables share identical sense-codon mappings and differ only in their start-codon assignments: tables 1 (Standard) and 11 (Bacterial / Archaeal / Plant Plastid), and tables 27 (Karyorelict Nuclear) and 28 (Condylostoma Nuclear). Both pairs are retained as separate entries to match NCBI numbering, but produce identical results for analyses that depend only on codon→amino-acid mappings; the 27 NCBI tables thus correspond to 25 distinct sense-codon colorings. Each table was tested against its own quartet-pattern shuffle null (*n* =10,000 per table, common-seed design with seeds = base seed + table ID). *P*-values were corrected for multiple comparisons using the Benjamini and Hochberg (1995) false discovery rate (FDR) procedure. For NCBI tables with dual-function stop/ sense codons (e.g., table 27 lists UGA as “Stop or Trp”; table 28 lists UAA, UAG as “Gln or Stop” and UGA as “Trp or Stop”), the primary codon→amino-acid analyses use the amino-acid label given in the NCBI AAs row of the gc.prt definition; stop functionality is handled only in analyses explicitly involving stop labels.

#### 2.3.3 Graph family ***G_ρ_*** and rho sweep

We define a family of mutation graphs *G_ρ_* parameterized by *ρ* ∈ [0, 1], where *G*_0_ = *Q*_6_(192 Hamming-1 edges) and *G*_1_ = *H*(3, 4) (all 288 single-nucleotide substitution edges). The weighted mismatch score on *G_ρ_* is

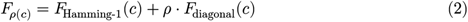

where *F*_diagonal_ sums over the 96 within-nucleotide distance-2 edges. This formulation treats *Q*_6_ not as the biological object but as one endpoint of a continuous interpolation toward the full mutation graph. We evaluated five values of *ρ* ∈ {0, 0.25, 0.5, 0.75, 1} with *n* =10,000 quartet-pattern shuffie null samples per value (common-seed design). Conservative p-values, computed as 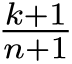 where *k* counts null samples with *F*_null_ ≤ *F*_obs_ (ties at the observed score are included in *k*): *p* (*k* = 61; 60 *_ρ_*_=0_ = 0.0062 strict + 1 tie), *p_ρ_*_=0.25_ = 0.0023 (*k* = 22), *p_ρ_*_=0.5_ = 0.0007 (*k* = 6), *p_ρ_*_=0.75_ = 0.0003 (*k* = 2), *p_ρ_*_=1_ = 0.0003 (*k* = 2). The last two are near the Monte Carlo resolution limit 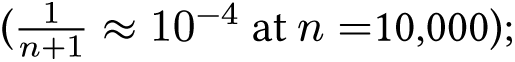 the qualitative conclusion (that the observed code lies in the extreme low-cost tail at every *ρ*) does not depend on *n*.

#### 2.3.4 Table-preserving permutation null (topology avoidance)

To test whether natural codon reassignment events avoid disrupting amino acid connectivity, we define a single primary candidate universe of single-codon reassignments and treat all reasonable alternatives as a sensitivity analysis. From the standard code *C*, the primary candidate set ℳ(*C*) pairs each of the 64 codons with each of the 20 alternative labels drawn from A_20_ ∪ {Stop}:

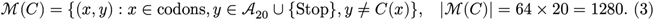

This **21-label, identity-excluded** universe (denoted U1 in Supplement §S5) gives every codon exactly 20 alternative labels, admits biologically attested stop-codon reassignment, and excludes identity moves (*y* = *C*(*x*)), which contribute no signal. Two strict variants (the **amino-acid-only, identity-excluded** universe with |ℳ| = 61 × 19 + 3 × 20 =1,219 (U2) and the **stop-inclusive with no-ops** universe with |ℳ| = 64 × 21 =1,344 (U4)) are reported as sensitivity analyses in Supplement §S5; topology-avoidance results are qualitatively identical under all three.

For each amino acid *a* and each adjacency *A* ∈ {*Q*_6_, *H*(3, 4)}, let *G_A_*^a^ denote the subgraph induced on codons assigned to *a* with edges given by adjacency *A* (*Q*_6_: binary Hamming distance ≤ 1 under the default encoding; *H*(3, 4): single-nucleotide substitution). A reassignment is *topology-breaking* under adjacency *A* if it increases the number of connected components of *G_A_*^a^ for any amino acid *a* (the **increase-in-components** definition Δ*β*_0_ (*G_A_*^a^) > 0 used by the conditional logit; the **new-disconnection-in-previously-connected-family** definition is reported in parallel as a sensitivity check; see Supplement §S3). The primary topology-avoidance test uses *A* = *H*(3, 4); the *A* = *Q*_6_ counterpart is reported as an encoding-dependent decomposition (§3.4, Supplement §S4). From the 27 NCBI translation tables analyzed in this work (Section 2.3.2), we catalogued the table-specific reassignment events relative to the standard code; de-duplicating by (codon, target amino acid) tuple yields the unique-event set on which the hypergeometric and table-preserving permutation tests operate. The conditional logit model (Section 2.3.5) operates on the full table-specific event-step list, which preserves recurrent reassignments across independent lineages.

Two tests were applied: (i) a hypergeometric test treating the observed events as a sample from the finite candidate landscape (*N* =1,280, *K* = 846 topology-breaking under the primary cell of *H*(3, 4) adjacency with Δ*β*_0_ > 0 definition, against *n* = 28 observed, *x* = 6 topology-breaking observed); and (ii) a table-preserving permutation test (*n* =10,000, seed 135325) that permutes target amino acids among a table’s reassigned codons, preserving within-table codon and target structure to address phylogenetic non-independence. Risk ratios with 95% log-normal confidence intervals were computed as RR = (*x*/*n*)/(*K*/*N*). The full 2 × 2 definition × adjacency audit (*Q*_6_ vs *H*(3, 4) adjacency × new-disconnection vs Δ*β*_0_ > 0 definitions) is given in Supplement §S3; risk ratios fall in 0.28–0.33 across all four cells.

#### 2.3.5 Conditional logit model of reassignment choice

To test whether topology avoidance contributes independent explanatory power beyond physicochemical optimization, we fit event-level conditional logit (discrete-choice) models to natural reassignment events. Each observed reassignment is treated as a choice from the set of all ≈1,280 possible single-codon reassignments available at the current code state. For a table with *k* reassignment events, the code state evolves sequentially: at each step, the model assigns a probability to each candidate move based on a linear utility score, and the observed move is compared against all alternatives.

For each candidate move *m* from code state *C*, we computed three features: (i) Δ_phys_, the change in local Grantham mismatch cost summed over Hamming-1 edges incident to the reassigned codon; (ii) Δ_topo_, the total increase in connected components across all amino acid codon graphs at *ε* = 1; and (iii) Δ_tRNA_, the Hamming distance from the reassigned codon to the nearest codon already encoding the target amino acid (a heuristic proxy for tRNA repertoire disruption). The conditional logit probability of observing move *m*^∗^ is

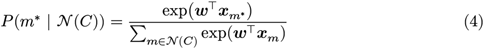

where ***w*** is the weight vector and ***x****_m_* is the feature vector for move *m*. Features were *z*-scored across all candidates for numerical stability.

Since the temporal ordering of reassignment events within a table is unknown, we marginalized the likelihood over all *k*! orderings for tables with *k* > 1 changes:

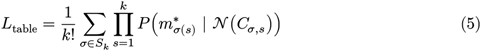

For tables with *k* ≤ 6 events (including the largest, yeast mitochondrial, *k* = 6), we enumerate all *k*! orderings exactly (up to 720 for *k* = 6); for the rare cases of *k* > 6, we sample 720 random orderings with the seeded RNG. The total likelihood is the product across all 25 tables with reassignment events (66 total event-steps).

Four candidate models were compared under *Q*_6_ topology; likelihood-ratio tests were applied only to nested pairs (M1 and M2 are non-nested alternatives, both nested within M3; M3 is nested in M4): M1 (physicochemistry only, *w*_topo_ = *w*_tRNA_ = 0), M2 (topology only, *w*_phys_ = *w*_tRNA_ = 0), M3 (physicochemistry + *Q*_6_ topology, *w*_tRNA_ = 0), and M4 (all three features). Two additional verification variants were fit using the encoding-independent *H*(3, 4) topology feature in place of *Q*_6_: M2_H(3,4)_ (topology only, *H*(3, 4)) and M3_H(3,4)_ (physicochemistry + *H*(3, 4) topology). The *H*(3, 4) variants test whether M3 dominance is robust to the choice of topology graph; the 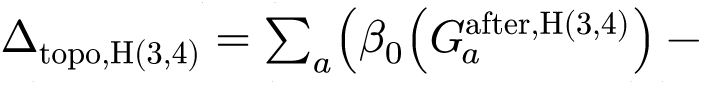 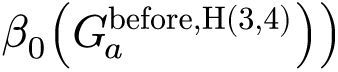 feature replaces the *Q*_6_ component-count change. Weights were estimated by maximum likelihood (scipy.optimize, Nelder–Mead with L-BFGS-B refinement). Model comparison used the corrected Akaike Information Criterion (AICc) and likelihood-ratio tests for nested pairs.

#### 2.3.6 Fisher–Stouffer test for tRNA enrichment

For each variant-code organism paired with a phylogenetically proximate standard-code control, we built a 2 × 2 contingency table comparing the proportion of tRNA genes encoding the reassigned amino acid versus all other amino acids. The sampling model treats tRNA gene counts as draws from a hypergeometric distribution conditional on the row and column marginals (total tRNAs per organism, total tRNAs for the focal amino acid), which is appropriate when the question is whether a specific amino acid’s share of the tRNA repertoire differs between two genomes. Fisher’s exact test (one-sided, alternative = “greater”) was applied per pairing, and *p*-values were combined via Stouffer’s *Z* method. To address non-independence from shared control organisms, we constructed a conflict graph (edges connect pairings sharing an organism) and enumerated all maximal independent sets (MIS) via the Bron–Kerbosch algorithm applied to the complement graph. For each MIS with ≥ 2 members we computed the Stouffer combined *p*-value and report the full distribution across MIS (best, median, worst) together with a pre-specified topology-breaking subset (*n* = 4 pairings whose underlying reassignment creates a new amino-acid disconnection under *Q*_6_: yeast mitochondrial Thr, *Scenedesmus* mitochondrial Leu, *Pachysolen* nuclear Ala, *Candida* nuclear Ser) as the mechanistic complement.

tRNA gene counts for 18 organisms were generated in this work by running tRNAscan-SE 2.0.12 (Chan and Lowe, 2019) with Infernal 1.1.4 on NCBI genome assemblies (eukaryotic mode for ciliates and yeasts; bacterial mode for *Mycoplasmoides*). Fifteen of these 18 tRNAscan-verified assemblies populate the 24-pairing enrichment analysis as either variant-code or standard-code-control repertoires; the remaining three (*Blastocrithidia nonstop* P57, *Mycoplasmoides genitalium* G37, *Mycoplasmoides pneumoniae* M129) are used only as mechanistic boundary cases in §4.3 because their reassignment routes act through anticodon-stem or base-modification changes on a single tRNA gene without changing gene-count distributions. The primary enrichment set is 24 variant–control pairings involving 24 unique organisms and 48 repeated organism-slots, drawn from NCBI translation tables 3 (*Saccharomyces cerevisiae* mitochondrial CUN→Thr), 6 (ciliate UAR→Gln), 10 (*Euplotes* UGA→Cys), 12 (*Candida albicans* CUG→Ser), 15 (*Blepharisma stoltei* UGA→Trp via a dedicated suppressor tRNA-Trp(UCA) (Singh et al., 2023), notwithstanding the legacy NCBI table-15 UAG→Gln nomenclature), 22 (*Scenedesmus obliquus* mitochondrial CUN→Leu; the chlorophycean mitochondrial equivalent of table 16), and 26 (*Pachysolen tannophilus* CUG→Ala). The remaining nine organism-slots have tRNA counts sourced from primary literature (including *S. cerevisiae* mitochondrial from Bonitz et al. (1980) and *Yarrowia lipolytica* mitochondrial from Kerscher et al. (2001)), from GtRNAdb (Chan and Lowe, 2016), or from published genome annotation; each is enumerated row-by-row with its compartment, source class, and assembly accession or primary citation in Supplement Table S11 (§S10.1). The *S. cerevisiae* mitochondrial Thr pairing is retained in the primary set because it is the sole tested example of a topology-breaking CUN→Thr reassignment (it appears in every maximal independent set of the pairing conflict graph); its non-tRNAscan provenance is called out separately in §S10.4.

### 2.4 Retrospective cross-study reanalysis of synthetic recoding outcomes

To test whether GF(2)6 topology predicts codon recoding outcomes in synthetic biology experiments, we performed a retrospective cross-study reanalysis across nine published genome recoding datasets, with quantitative analysis of eight (the ninth, (Ding et al., 2024), is included for cross-kingdom scope without quantitative extraction). For each codon substitution reported in these studies, we computed three topology features under the default encoding (*C* ↦ (0, 0), *U* ↦ (0, 1), *A* ↦ (1, 0), *G* ↦ (1, 1)):

1. **Boundary crossing** (*ε* = 1): whether the source and target codons lie in different connected components of the amino acid’s codon graph at Hamming distance 1. Generalized beyond serine to all amino acids with potential disconnections (including leucine, arginine).
2. **Local mismatch change** (Δ*F*_local_): the difference in total Grantham distance summed over all Hamming-1 neighbors between the target and source codon positions: Δ*F*_local_ = ∑*_n_*_∈*N*(*t*)_ Δ(*c*(*t*), *c*(*n*)) − ∑*_n_*_∈*N*(*s*)_ Δ(*c*(*s*), *c*(*n*)), where *N*(*v*) denotes the 6 Hamming-1 neighbors of vertex *v* in *Q*_6_.
3. **Hamming distance**: the number of bit positions differing between source and target vectors in GF(2)6.

Codon positions were extracted from published GenBank genome files by CDS-level comparison using Biopython (Cock et al., 2009), with codon frame offsets (`/codon_start`) handled explicitly. CDS features were matched between parent and recoded genomes by locus tag. The primary analysis used the Syn57 dataset (Robertson et al., 2025), which provides a within-study contrast: 37,146 serine recodings cross the UCN↔AGY family boundary, while 22,859 alanine recodings remain within the GCN family. Syn57 RNA-seq differential expression data (Data file S10) were used as the outcome measure. Design-to-final genome deviations were identified by comparing the vS33A7 design genome (Data file S2) against the verified Syn57 genome (Data file S8), filtering genes with >10 CDS-level differences as structural rearrangements (sensitivity analysis across thresholds 3–50 reported in the Supplementary Material). The Ostrov et al. (2016) segment viability data (Table S4) were analyzed as a case-control comparing segments with and without lethal design exceptions; Bonferroni correction was applied for three simultaneous tests.

All cross-study reanalysis code is released in the `codon_topo.analysis.codonsafe` subpackage of the `codontopo` repository (https://github.com/biostochastics/codontopo, version 0.6.1) and can be run via `codon-topo codonsafe`. Raw data files and download provenance are documented in `data/ codonsafe/DATA_MANIFEST.md`.

### 2.5 Structural-preservation index (visualization-only)

For Figure 5A we use a heuristic **structural-preservation index** *S*(*m*) ∈ [0, 1] that combines four discrete indicators for each candidate single-codon reassignment *m*: whether the post-reassignment code preserves the two-fold bit-5 filtration (*I*_tw_), whether it preserves the four-fold prefix filtration (*I*_ff_), whether Serine remains disconnected at *ε* = 1 (*I*_ser_), and how many amino acids are disconnected at *ε* = 1 (*n*_disc_). Explicitly

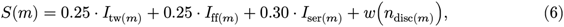

where *w*(1) = 0.20, *w*(0) = 0.10, and *w*(*n*) = 0.00 for *n* ≥ 2. Because every component is discrete, *S*(*m*) takes only four values across the 1,280-move candidate universe (*S* ∈ {0.55, 0.75, 0.80, 1.00}; 193 / 27 / 738 / 322 events; the four visible bars in Figure 5A). The weights are a hand-tuned prioritisation of the four structural features and are not derived from any calibration; the index is descriptive, not inferential, and is not used in any inferential test in this paper. Exact implementation: Supplement §S12 and `src/codon_topo/analysis/synbio_feasibility.py` of the `codontopo` repository.

### 2.6 Software and reproducibility

The analysis pipeline is implemented in Python 3.11 (NumPy, SciPy, statsmodels, joblib) with R 4.4 (ggplot2, ggpubr) for figures and Typst 0.14 for typesetting; source at https://github.com/ biostochastics/codontopo. All Monte Carlo procedures use seed 135325; Python dependency versions are pinned in `requirements.lock`. Use of generative-AI systems in scaffolding analysis code is disclosed in the end-of-manuscript declaration.

### 2.7 Claim hierarchy

All 15 scientific claims evaluated in this work are registered in a formal claim hierarchy with pre-specified status categories (Table 1): SUPPORTED (meets its prespecified claim-specific decision rule under the stated null or model and multiplicity control), EXPLORATORY (descriptive or hypothesis-generating; does not meet the relevant confirmatory gate), REJECTED (falsified or pre-rejected in literature), FALSIFIED (directly contradicted by data in this or prior work), and TAUTOLOGICAL (true by construction). Each claim’s decision rule is stated in the “Justification” column of the full hierarchy in the supplementary materials.

**Table 1:** Summary of 15 evaluated claims. The implemented null is a **quartet-pattern shuffle** (16 first-two-base quartets, each with its internal AA pattern held atomic and shuffied across quartet slots) rather than the classical Freeland–Hurst amino-acid permutation across synonymous codon groups. “MIS” = maximal independent set enumeration. *p*-values are the primary summary per claim (MIS median for the tRNA row; most conservative test elsewhere). Shaded rows are non-inferential: FALSIFIED and REJECTED rows record claims found to be wrong, and TAUTOLOGICAL rows record claims that are true by construction and carry no evidentiary weight.

| Status | Claim | Null model | *p*-value |
| --- | --- | --- | --- |
| Supported | Cross-metric coloring optimality | quartet-pattern shuffle × 4 metrics | $\leq 0.0062$ |
| | Per-table optimality (26/27) | Per-table quartet-pattern shuffle + BH-FDR | $< 0.05$ |
| | ρ-robustness ($\rho \in [0, 1]$) | Quartet-pattern shuffle (weighted edges) | 0.006 |
| | Topology avoidance ($Q_6 + H(3, 4)$) | Table-pres. perm. + clade excl. | $10^{-4}$ |
| Exploratory | tRNA enrichment (MIS median) | Fisher–Stouffer (332 MIS) | 0.037 |
| | Bit-position bias (dedup.) | $\chi^2$ uniform | 0.075 |
|  | Mechanism boundary conditions | Descriptive | — |
|  | Atchley F3/Serine convergence | Qualitative | — |
|  | Disconnection catalogue (4 cases) | Descriptive | — |
| Falsified | KRAS–Fano clinical prediction | Fisher–Bonferroni | 1.0 |
| Rejected | Serine min-distance-4 invariant | Counterexample | — |
|  | PSL(2,7) symmetry | Literature | — |
|  | Holomorphic embedding | Verification | — |
| Tautological | Two-fold bit-5 filtration | — | — |
|  | Four-fold prefix filtration | — | — |

**Table 2:** Cross-metric sensitivity analysis. Quartet-pattern shuffie null (*n* = 10,000, seed 135325) under four code-independent physicochemical distance measures and the alignment-derived, code-dependent ProtSub matrix (Jia and Jernigan, 2021) as a structure-aware sensitivity analysis. All five measures show significant optimality after Bonferroni correction at *α* = 0.05 (per-test threshold *α*/5 = 0.01). Effect size *z* = (*μ*_null_ − *F*_obs_)/*σ*_null_. “Improv.” = percent improvement of observed score over null mean. Under a weaker degeneracy-only null (preserving only codon family sizes, not block structure), all measures yield *z* > 9 and *p* < 10^−4^. Under the quartet-pattern shuffie null, ProtSub places the standard code at the 0.04% quantile (*n* = 10,000), the most extreme percentile of any measure tested.

| Metric | Observed | Null mean ± SD | $z$ | $p$ | Improv. |
| --- | --- | --- | --- | --- | --- |
| Grantham (Grantham, 1974) | 13,477 | $14,954 \pm 628$ | 2.35 | 0.006 | 9.9% |
| Miyata (Miyata et al., 1979) | 235.3 | $275.7 \pm 12.9$ | 3.14 | $< 0.001$ | 14.6% |
| Polar requirement (Woese, 1973) | 347.3 | $410.5 \pm 23.1$ | 2.73 | 0.003 | 15.4% |
| Kyte–Doolittle (Kyte and Doolittle, 1982) | 457.2 | $570.7 \pm 40.9$ | 2.78 | 0.001 | 19.9% |
| ProtSub (Jia and Jernigan, 2021)<sup>1</sup> | 1089.5 | $1180.2 \pm 29.7$ | 3.05 | $5 \times 10^{-4}$ | 7.7% |

## 3 Results

### 3.1 Cross-metric coloring optimality

Under the quartet-pattern shuffle null with *n* = 10,000 permutations, the standard genetic code is significantly low-cost across all four physicochemical distance metrics examined (Table 2). For Grantham distance, the observed score is *F* = 13,477 versus a null mean of 14,954 ± 628 (*z* = 2.35, quantile 0.61%, *p* = 0.0062). The same pattern holds for Miyata distance (*F* = 235.3 vs 275.7 ± 12.9, *z* = 3.14, *p* < 0.001), Woese polar requirement (*F* = 347.3 vs 410.5 ± 23.1, *z* = 2.73, *p* = 0.003), and Kyte–Doolittle hydropathy (*F* = 457.2 vs 570.7 ± 40.9, *z* = 2.78, *p* = 0.001). Relative to null expectations, the observed code lowers the mismatch score by 9.9%, 14.6%, 15.4%, and 19.9%, respectively. Beta-posterior credible intervals for the empirical *p*-values remain entirely below 0.01 for all four metrics (computed from the conservative 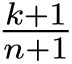 estimator with Beta(*k* + 1, *n* − *k* + 1) posterior; per-metric intervals are listed in `output/coloring_optimality.json` under `beta_credible_intervals`). Since Freeland and Hurst (1998) established optimality using polar requirement alone, the cross-metric concordance demonstrates that error-minimization is robust across multiple distinct physicochemical parameterizations, not an artifact of any single distance metric. Under a weaker degeneracy-only null that preserves only codon family sizes without maintaining block contiguity, the signal strengthens substantially (*z* > 9, *p* < 10^−4^ for all metrics), confirming that the quartet-pattern shuffle null provides a stringent test.

### Convergent operationalisation across five measures, including ProtSub

The four code-independent measures in Table 2 are intercorrelated by construction. Across the 190 unordered amino-acid pairs (20 standard AAs), pairwise Spearman correlations range from *ρ* = 0.36 (Grantham vs Kyte–Doolittle) to *ρ* = 0.73 (Miyata vs Woese polar requirement; median *ρ* = 0.49; full matrix in Supplement §S20). At the code level (ranking 2,000 random codes drawn from the quartet-pattern shuffle null by their *F*-scores), pairwise Spearman correlations are higher (range 0.71–0.89, median 0.82), as expected. To probe whether the panel captures non-identical residual structure rather than a single latent factor, we computed **partial** Spearman correlations controlling simultaneously for all other measures: four of ten pairs become approximately uncorrelated after adjustment (|*ρ*_partial_| < 0.1; a descriptive point-estimate statement, not a test of statistical independence): Grantham vs polar requirement (*ρ*_partial_ = +0.03), Miyata vs ProtSub (−0.06), polar requirement vs Kyte–Doolittle (+0.01), and polar requirement vs ProtSub (+0.08). Substantial shared variance coexists with non-identical residual structure across the panel; we therefore describe the cross-metric concordance as a convergent sensitivity envelope rather than as independent replication. Under the same quartet-pattern shuffle null, the alignment-derived ProtSub matrix (Jia and Jernigan, 2021) places the standard code at the most extreme percentile of any measure tested (Table 2). With five tests at *α* = 0.05, the Bonferroni-corrected per-test threshold is 0.01; all five p-values satisfy it (Grantham *p* = 0.0062; others *p* ≤ 0.003). ProtSub is presented as a code-dependent sensitivity analysis rather than as primary evidence for code-origin optimality (Section 2.2): Buschmann et al. (2026) demonstrate that all substitution-matrix families (BLOSUM62, AlphaFold-derived AFSM, and 16 others) perform similarly because they implicitly encode the same physicochemical reality, which bounds rather than eliminates the Di Giulio (2001) circularity concern for alignment-derived matrices.

Decomposing *F* by nucleotide position (Figure 5, panel B) reveals that the second codon position contributes 49.3% of the total mismatch, the first position 38.2%, and the wobble position only 12.5%. This gradient mirrors the biochemical hierarchy of mutational impact: second-position changes are the most physicochemically disruptive, first-position changes are intermediate, and wobble-position changes are largely synonymous (Freeland and Hurst, 1998, Woese, 1965).

### 3.2 Robustness across diagonal-edge weight ρ

The weighted score *F_ρ_* (Equation 2) remains significantly optimal across all values of *ρ* from 0 to 1 (Table 3). At *ρ* = 0 (pure *Q*_6_), *p* = 0.0062 (*z* = 2.35); at *ρ* = 1 (full *H*(3, 4), all 288 single-nucleotide edges equally weighted), *p* < 0.001 (*z* = 3.37). No value of *ρ* yields *p* > 0.0062. The optimality signal strengthens monotonically as diagonal edges are included, addressing the concern that the *Q*_6_ representation ignores approximately one-third of single-nucleotide mutations.

**Table 3:** Robustness of coloring optimality across diagonal-edge weight *ρ*. Quartet-pattern shuffie null, *n* = 10,000 per *ρ*. All *p*-values < 0.01. Quantiles reported as empirical rank; values ≤ 0.02% indicate ≤ 2 null samples below the observed score.

| $\rho$ | Observed | Null mean | Quantile | $p$-value | $z$ |
| --- | --- | --- | --- | --- | --- |
| 0 | 13,477 | 14,954 | 0.61% | 0.0062 | 2.35 |
| 0.25 | 15,335 | 17,020 | 0.22% | 0.0023 | 2.77 |
| 0.5 | 17,193 | 19,085 | 0.06% | < 0.001 | 3.09 |
| 0.75 | 19,050 | 21,150 | 0.02% | < 0.001 | 3.28 |
| 1 | 20,908 | 23,216 | 0.02% | < 0.001 | 3.37 |

**Table 4:** Topology avoidance in natural codon reassignments under the encoding-independent *H*(3, 4) Hamming graph (primary cell). Topology-breaking = Δ*β*_0_ > 0, an increase in the total number of connected components summed across amino acid codon graphs. The **Expected (chance)** column shows what would be observed under the null hypothesis that the 28 observed events are drawn uniformly at random from the 1,280-move candidate landscape: *E*[breaks] = 28 × *K*/*N* ≈ 18.5 versus 6 observed. RR = risk ratio (observed rate / candidate rate) with 95% log-normal CI; depletion fold = (candidate rate)/(observed rate). The remaining three cells of the 2 × 2 definition × adjacency audit (*Q*_6_ vs *H*(3, 4) × new-disconnection vs Δ*β*_0_ > 0) are reported in Supplement §S3 (Table 5 of this manuscript reproduces them inline), with risk ratios in the range 0.28–0.33 and hypergeometric *p* < 10^−5^ throughout.

| Quantity | Observed | Candidate landscape | Expected (chance) |
| --- | --- | --- | --- |
| Topology-breaking ($\Delta\beta_0 > 0$) | 6 (21.4%) | 846 (66.1%) | 18.5 (66.1%) |
| Topology-preserving ($\Delta\beta_0 = 0$) | 22 (78.6%) | 434 (33.9%) | 9.5 (33.9%) |
| Total | 28 | 1,280 | 28 |
| Depletion fold (cand. rate / obs. rate) | — | $3.1\times$ | $1.0\times$ |
| RR (95% CI) | — | 0.32 (0.16–0.66) | 1.00 (ref.) |
| Hypergeometric $p$ (one-sided) | — | $1.28 \times 10^{-6}$ | (null reference) |
| Permutation $p$ (table-pres.) | — | $\leq 10^{-4}$ | (null reference) |

### 3.3 Per-table optimality preservation

Of the 27 NCBI translation tables, 26 remain in the top 5% of their own quartet-pattern shuffie null after Benjamini–Hochberg FDR correction (Figure 2, panel C). The mean quantile across all tables is 1.3%. The single marginal exception is translation table 3 (yeast mitochondrial code, 6 codon reassignments; *p*_BH_ =0.073).

**Figure 1:**
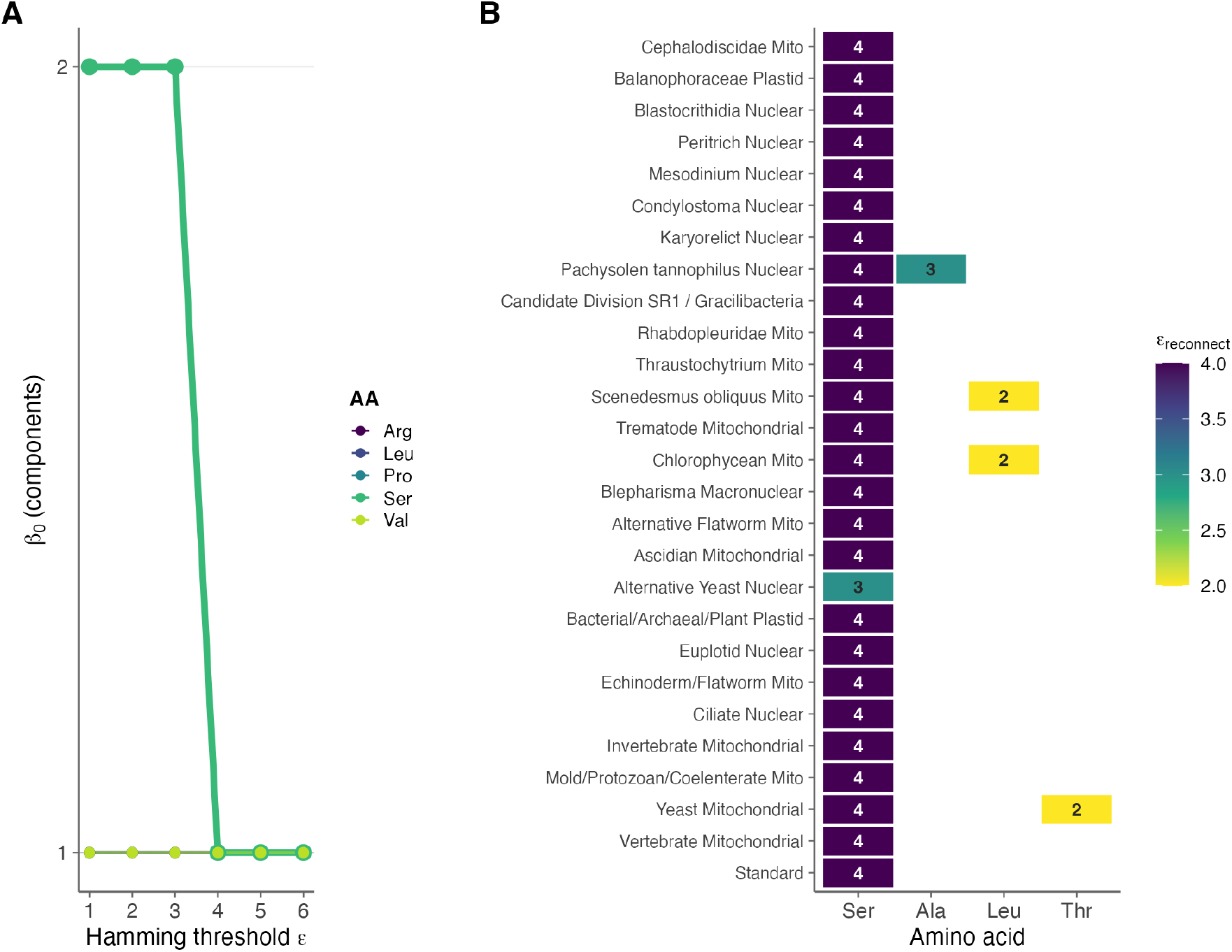
Core topology of the genetic code in GF(2)^6^. **(A)** Persistent homology: connected components (*β*_0_) of amino acid codon graphs as a function of Hamming distance threshold *ε*. Serine (bold) is the only amino acid disconnected at *ε* = 1 across all 27 NCBI translation tables (encoding-invariant); under the default encoding it reconnects at *ε* = 4 (reconnection depth is not encoding-invariant; 16 of 24 encodings give *ε* = 2). **(B)** Disconnection catalogue across all translation tables. Each tile denotes an amino acid disconnected at *ε* = 1; the number is the reconnection *ε* under the default encoding. Serine is universally disconnected; Leu, Thr, and Ala appear as variant-code disconnections.

**Figure 2:**
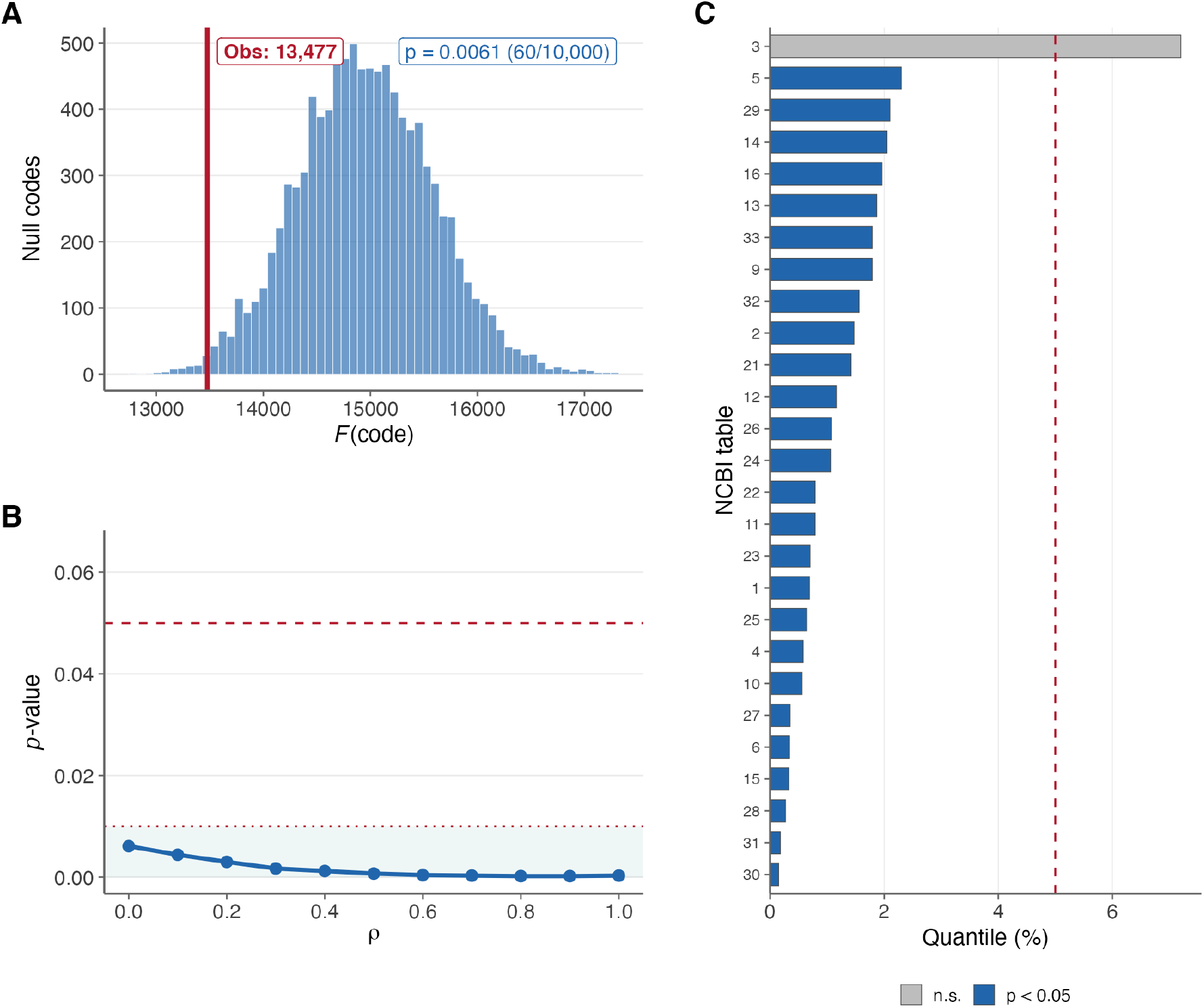
Coloring optimality of the genetic code. **(A)** Null distribution of Grantham edge-mismatch scores *F* under the quartet-pattern shuffle null (*n* = 10,000). The observed standard code (red line, *F* = 13,477) falls at the 0.61st percentile (*p* = 0.0062). **(B)** *p*-value across diagonal-edge weight *ρ* from 0 (*Q*_6_) to 1 (full *H*(3, 4)); all values below 0.01, with optimality strengthening monotonically. The curve interpolates over 11 sample points shown as markers; the 5 pre-specified inferential grid values *ρ* ∈ {0, 0.25, 0.5, 0.75, 1} (Table 3) are used for multiplicity correction, and the remaining 6 points are descriptive interpolation shown for shape only. **(C)** Per-table quantile for each of 27 NCBI translation tables under their own quartet-pattern shuffle null (*n* =10,000); only table 3 (yeast mito.) exceeds the 5% threshold.

**Figure 3:**
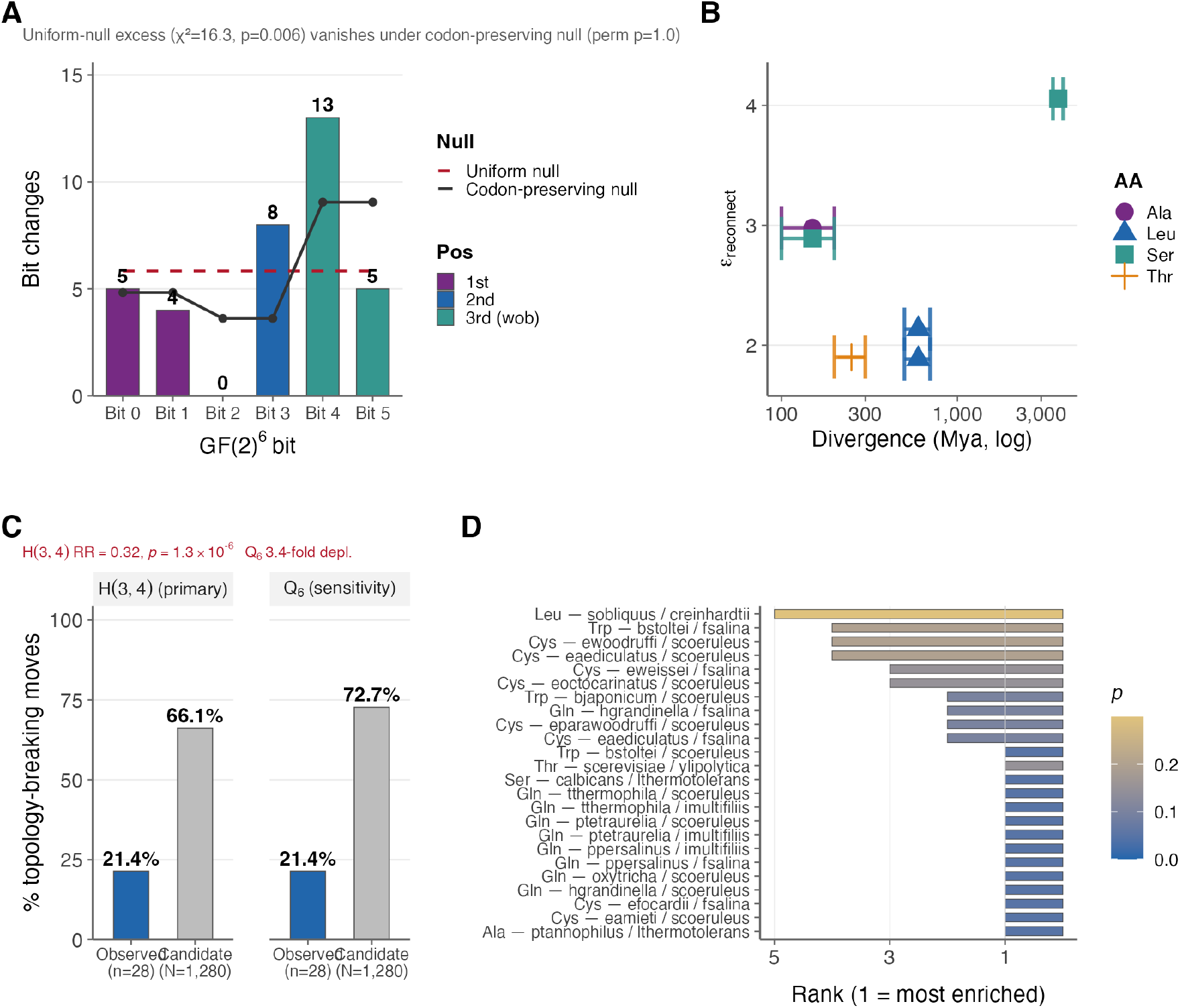
Evolutionary evidence for structural constraints on codon reassignment. **(A)** Bit-position bias: distribution of bit-flips across GF(2)^6^ coordinates in natural reassignment events; dashed line = uniform expectation. **(B)** Evolutionary depth calibration: reconnection *ε* vs estimated divergence age (log scale) — 6 calibration points spanning 4 amino acids (universal Serine as the pre-LUCA baseline plus variant-code Thr, Leu, and Ala; Serine and Leucine each contribute two points from different lineages). **(C)** Topology avoidance shown for the encoding-independent *H*(3, 4) Hamming graph (left panel, primary, using the **increase-in-components** Δ*β*_0_ > 0 definition matched to the conditional-logit feature) and the encoding-dependent *Q*_6_ subgraph (right panel, sensitivity, using the **new-disconnection** definition for continuity with the paper’s *Q*_6_ narrative). Under *H*(3, 4), observed natural reassignments break topology at 21.4% versus 66.1% of the candidate landscape (RR 0.32, permutation *p* ≤ 10^−4^). The four-cell 2 × 2 audit (definition × adjacency) is reported in Supplement §S3 and yields risk ratios 0.28–0.33 across all cells. **(D)** tRNA enrichment: rank of the reassigned amino acid among all 20 AAs by tRNA gene proportion in variant-code vs standard-code organism pairings; rank 1 = most enriched.

The headline number conflates two distinct evidentiary regimes, which we disaggregate. Variant tables differing from the standard code by ≥ 3 reassignments (“informative-distance” tables) sit far enough away that their per-table quartet-shuffle null contains many distant permutations; tables with ≤ 2 reassignments (“near-standard” tables) sit so close that their null is dominated by near-standard permutations. A standard-code-proximity audit (Supplement §S8) makes the rationale explicit: every variant sits at the extreme low-*d_H_* tail of its own null (every null draw has *d_H_* ≥ 30, while every variant has *d_H_* ≤ 6), so no distance-matched null draw exists. The per-table *p*-value therefore **cannot** separate proximity-driven from independently-optimized codes: it measures whether each variant is low-cost **relative to its own quartet-shuffle ensemble**, not whether that optimality is independent of standard-code proximity. The informative-distance vs near-standard split rests on the absolute *d_H_* ≥ 3 cutoff, a descriptive threshold on reassignment distance rather than an identified boundary between regimes.

Of the 12 informative-distance **registry entries**, 11 retain top-5% placement of their own quartet-shuffie ensembles after BH–FDR correction, the lone marginal case being yeast mitochondrial. Because NCBI tables 27 and 28 (Karyorelict and Condylostoma nuclear codes) share identical sense-codon mappings, this registry corresponds to 11 distinct sense-codon colorings, 10 of which retain top-5% placement. Of the 14 near-standard tables, 14 are formally significant under the same correction, but those results reflect proximity to the standard code rather than independent variant-specific optimization. The descriptive claim is therefore that, among variant codes distant enough for the per-table test to admit a nontrivial range of permutations, all but one preserve low physicochemical cost relative to their own ensemble. Even the most reassigned code (table 3, 6 codons reassigned) sits at the 7.4th percentile of its quartet-shuffle null, marginally above the 5% threshold but still in the lower tail.

### 3.4 Topology avoidance in natural codon reassignments

We adopt the encoding-independent *H*(3, 4) Hamming graph as the primary adjacency for the topology-avoidance test, with the *Q*_6_ subgraph reported as a coordinate-dependent decomposition (motivated below). Under *H*(3, 4), of the 1,280 candidate single-codon relabelings (Methods §2.3.4), 846 (66.1%) increase the connected-component count of some amino acid’s codon graph relative to the standard code (Δ*β*_0_ > 0). Only 6 of 28 observed natural reassignment events (21.4%) do so, yielding a 3.1-fold depletion (RR = 0.32, 95% CI [0.16, 0.66]; hypergeometric *p* = 1.28 × 10^−6^; table-preserving permutation *p* ≤ 10^−4^).

Under *Q*_6_ (Hamming-1 adjacency in the default GF(2)^6^ encoding), the corresponding numbers are 931 of 1,280 candidates (72.7%) creating new amino-acid disconnections, against only 6 of 28 observed events (21.4%); *Q*_6_ depletion is 3.4-fold (hypergeometric *p* = 1.58 × 10^−8^; permutation *p* ≤ 10^−4^). The Q_6 candidate-rate is somewhat higher than the H(3,4) rate because some *Q*_6_-disconnected pairs become connected when within-nucleotide Hamming-2 edges are admitted. Both adjacencies yield highly significant depletion under both topology-breaking definitions (new disconnection in a previously connected family; Δ*β*_0_ > 0 increase in components), with risk ratios 0.28–0.33 across the four cells of the 2 × 2 definition × adjacency audit (Supplement §S3) — the avoidance is not specific to either the hypercube subgraph or either definition. We give precedence to the *H*(3, 4) result because *Q*_6_ is encoding-dependent: across all 24 base-to-bit bijections, the *Q*_6_ candidate-landscape rate ranges from 36% (8 encodings) to 73% (default encoding), with median hypergeometric *p* = 6 × 10^−6^ but *p* > 0.5 in 8 of 24 encodings (Supplement §S4). *H*(3, 4), by contrast, depends only on nucleotide identity and is robust by construction.

Following Sengupta et al. (2007), we performed clade-exclusion sensitivity analysis, iteratively removing each major taxonomic group (all ciliates, all metazoan mitochondria, all CUG-clade yeasts, etc.) and retesting. Depletion is significant in every one of the seven regimes, but the threshold depends on the topology-breaking definition: under the primary *H*(3, 4)/Δ*β*_0_ > 0 definition all seven exclusions satisfy *p* < 10^−3^ (largest *p* ≈ 2.03 × 10^−4^, from excluding all metazoan mitochondria); under the sensitivity *Q*_6_/new-disconnection definition all seven satisfy *p* < 10^−5^. The result nonetheless requires cautious interpretation, since the topology-breaking events are not broadly distributed across clades. The full 7-row clade-exclusion table is persisted for both the *Q*_6_ / new-disconnection adjacency and the primary *H*(3, 4) / Δ*β*_0_ > 0 adjacency (Supplement §S9). For the yeast-mitochondrial-excluded regime the direct calculation is: 4 of the 6 *H*(3, 4) breakers come from a single clade (yeast mitochondrial, NCBI translation table 3; all 4 of its de-duplicated reassignments are *H*(3, 4)-breakers), and excluding it leaves 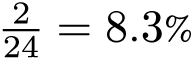 breakers, moving the *H*(3, 4), Δ*β*_0_ > 0 hypergeometric *p* from 1.28 × 10^−6^ to ≈ 4.2 × 10^−9^ (direct one-sided calculation on *N* = 1280, *K* = 846, *n* = 24, *x* = 2). The *Q*_6_ / new-disconnection variant of the same single-row calculation gives *p* ≈ 3.6 × 10^−11^ (Supplement §S9) and is not the primary cell. The clade-exclusion robustness is thus a denominator effect: removing clades that contributed zero or one breaker each leaves the breakage rate essentially unchanged, rather than evidence of repeated, independently arising topology-breaking events across many lineages. The cross-lineage signal is therefore the **avoidance** of topology-breaking moves, supported by 22 of 28 events across many lineages; the cross-clade structure of the breaker set itself is underpowered.

NCBI translation table 3 (yeast mitochondrial) appears as the marginal case in three diagnostics: per-table optimality (§3.3), the lowest-ranking observed move under the M3 conditional logit (§3.5), and the topology-breaker concentration here. These three observations share a *k*-magnitude effect by construction (table 3 has the most reassignments in the registry, and is the only variant code requiring acquisition of a novel tRNA from a different parent: tRNA^Thr^ derived from tRNA^His^ (Su et al., 2011)), so we do not interpret their convergence as independent confirmation of any specific selectionist mechanism. The pattern is consistent with topology-breaking reassignments being feasible but appearing concentrated in unusually perturbed translation systems.

All four cells of the adjacency × definition audit (Table 5) yield hypergeometric *p* < 10^−5^ with risk ratios 0.28–0.33, so the topology-breaking definition matters little. The encoding-sensitivity audit matters more: the *H*(3, 4) result is constant across all 24 base-to-bit bijections, but under the 8 encodings placing the *Q*_6_ candidate rate near 36% the observed rate (21–36%) does not differ from it (*p* > 0.5). We therefore treat *H*(3, 4) as the primary topology-avoidance result and *Q*_6_ as a coordinate-dependent decomposition (Supplement §S4). Denominator sensitivity (alternative candidate universes |*M*| = 1219, |*M*| = 1344) is reported in Supplement §S5 and does not change the qualitative conclusion.

**Table 5:** Sensitivity analysis: the 2 × 2 topology-breaking definition × adjacency audit. All four cells share the same denominator (1,280 candidates) and the same observed-event total (28 de-duplicated reassignments); the observed topology-breaking count is 5–7 of 28 across cells. All four cells yield depletion in the same direction with comparable risk ratios (0.28–0.33). The *H*(3, 4) Hamming graph is encoding-independent; the *Q*_6_ subgraph is encoding-dependent and 8 of 24 base-to-bit bijections give no *Q*_6_ depletion at all (Supplement §S4). The main text uses the *H*(3, 4), Δ*β*_0_ > 0 cell as primary.

| Adjacency | Definition | $K/N$ | Hyper. $p$ | RR (95% CI) |
| --- | --- | --- | --- | --- |
| $H(3, 4)$ (primary) | $\Delta\beta_0 > 0$ | 846 / 1,280 | $1.28 \times 10^{-6}$ | 0.32 (0.16–0.66) |
| $H(3, 4)$ | new disconnection | 822 / 1,280 | $4.95 \times 10^{-7}$ | 0.28 (0.13–0.62) |
| $Q_6$ | $\Delta\beta_0 > 0$ | 963 / 1,280 | $2.23 \times 10^{-8}$ | 0.33 (0.17–0.63) |
| $Q_6$ | new disconnection | 931 / 1,280 | $1.58 \times 10^{-8}$ | 0.29 (0.14–0.6) |

**Table 6:**
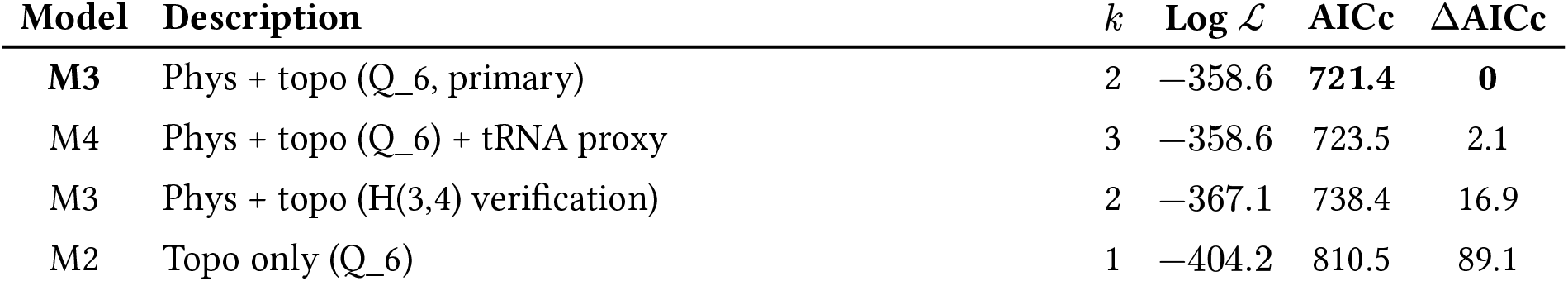
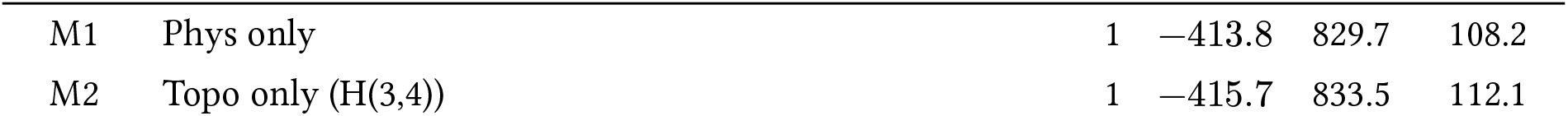
Event-level conditional logit model comparison. Six models share the same candidate set (≈ 1,280 single-codon moves per event-step) and the same likelihood structure marginalised over event orderings; they differ only in the feature subset used. The first four models use the encoding-dependent *Q*_6_ topology feature (the legacy primary, paired with the encoding sweep in §S4); the two M-variants ending in *H*(3, 4) replace Δ_topo,Q_6_ with the encoding-independent Δ_topo,H(3,4)_feature, providing the encoding-robustness verification. M3 (phys + topo) is favoured under both topology graphs; the gap between M3_Q_6_ and M3_H(3,4)_ reflects which topology feature carries more information per parameter, not whether topology is itself meaningful. *k* = number of estimated parameters. Bold = best model by AICc.

| Model | Description | $k$ | Log $\mathcal{L}$ | AICc | $\Delta AICc$ |
| --- | --- | --- | --- | --- | --- |
| M3 | Phys + topo ($Q_6$, primary) | 2 | −358.6 | 721.4 | 0 |
| M4 | Phys + topo ($Q_6$) + tRNA proxy | 3 | −358.6 | 723.5 | 2.1 |
| M3 | Phys + topo ($H(3,4)$ verification) | 2 | −367.1 | 738.4 | 16.9 |
| M2 | Topo only ($Q_6$) | 1 | −404.2 | 810.5 | 89.1 |

Table 6: Event-level conditional logit model comparison.
| Model | Description | $k$ | Log $\mathcal{L}$ | AICc | $\Delta\text{AICc}$ |
| --- | --- | --- | --- | --- | --- |
| M1 | Phys only | 1 | -413.8 | 829.7 | 108.2 |
| M2 | Topo only ($H(3,4)$) | 1 | -415.7 | 833.5 | 112.1 |

### 3.5 Explanatory modeling: topology as an independent predictor

To test whether topology avoidance is reducible to physicochemical optimization, we fit the six event-level conditional logit models of Section 2.3.5 to the 66 reassignment event-steps across 25 variant-code tables, treating each observed reassignment as a choice among ≈1,280 candidate single-codon moves (four share the encoding-dependent *Q*_6_ topology feature; two are *H*(3, 4) verification variants). The combined physicochemistry-plus-topology model under *Q*_6_ topology (M3) was strongly favored over all alternatives by AICc (AICc = 721.4), outperforming the physicochemistry-only model (M1; ΔAICc = 108.2) and the topology-only model (M2; ΔAICc = 89.1). The encoding-independent verification model M3_H(3,4)_ also outperforms the physicochemistry-only baseline (ΔAICc(M1 → M3_H(3,4)_) = 91.3) and the H(3,4)-topology-only baseline (ΔAICc(M2_H(3,4)_ → M3_H(3,4)_) = 95.1), so the conditional-logit thesis does not depend on the choice of topology graph. Adding a heuristic tRNA-complexity proxy did not improve fit (M3→M4: ΔAICc = 2.1).

Likelihood-ratio tests confirmed that each feature class adds substantial explanatory value to the other: adding topology to physicochemistry yields LR = 110.4 (*p* ≪ 10^−10^), and adding physicochemistry to topology yields LR = 91.2 (*p* ≪ 10^−10^). The two feature classes are only weakly associated across the full candidate landscape, indicating limited confounding.

In the best-fitting model (M3), observed natural reassignments preferentially populate moves that reduce local physicochemical mismatch (β^_phys_ = −0.004 per Grantham unit) and strongly avoid moves that increase amino acid codon-family disconnection (β^_topo_ = −3.32 per additional connected component, with the conditional-logit feature evaluated under *Q*_6_ adjacency using the **increase-in-components** (Δ*β*_0_ > 0) convention). Because *Q*_6_ adjacency is encoding-dependent (8 of 24 base-to-bit bijections give no *Q*_6_ topology depletion at the landscape level; Section 3.4), the encoding-independent *H*(3, 4) refit is the key check. The encoding-robustness comparison gives ΔAICc(M1 → M3_H(3,4)_) = 91.3 and ΔAICc(M2_H(3,4)_ → M3_H(3,4)_) = 95.1, both well above the conventional ΔAICc > 10 threshold treated as strong evidence in the model-comparison literature (Burnham and Anderson, 2002), and similar in magnitude to the *Q*_6_ counterparts (ΔAICc(M1 → M3_Q_6_) = 108.2). This threshold was empirically calibrated on linear-regression model-comparison contexts; we use it here as a conventional reference rather than as a formally calibrated cut-off in the conditional-logit setting. The M3 dominance is therefore not an artifact of the *Q*_6_ encoding choice. Under M3, observed natural reassignments rank on average at the 89.5th percentile among all candidate moves (Figure 4, panel B), with the recurrent UGA→Trp reassignment consistently ranking above the 98th percentile. The one outlier is the yeast mitochondrial CUU→Thr reassignment (30th percentile), consistent with NCBI translation table 3 being the sole marginal exception in the per-table optimality analysis (Section 3.3). The non-significance of the heuristic tRNA-distance proxy (LR = 0.12, *p* = 0.73) speaks to the inadequacy of Hamming-distance-to-nearest-target-AA-codon as a proxy for tRNA-mediated mechanistic feasibility; it should not be interpreted as evidence against tRNA effects in general (see Section 3.6 for direct tRNA-gene-count tests). A parametric predictive simulation under M3 reproduced the observed topology-breaking rate at the event-step level under *Q*_6_ adjacency using the **increase-in-components** (Δ*β*_0_ > 0) definition (observed 5/66 event-steps = 0.076 vs simulated mean 0.077 ± 0.033 over *n* =10,000 draws; parametric predictive *p* = 0.60; Supplement §S19.5, Figure S8 panel C). This event-step / *Q*_6_ / Δ*β*_0_ > 0 rate differs from the lineage-collapsed 6/28 = 0.21 rate quoted for *H*(3, 4) adjacency with the **increase-in-components** (Δ*β*_0_ > 0) definition in §3.4 (Section 3.4) — the same definition, but a different denominator (66 event-steps vs 28 events) and a different adjacency (*Q*_6_ vs *H*(3, 4)). The predictive check supports model calibration on its own denominator/adjacency/ definition combination rather than in-sample AICc improvement alone.

**Figure 4:**
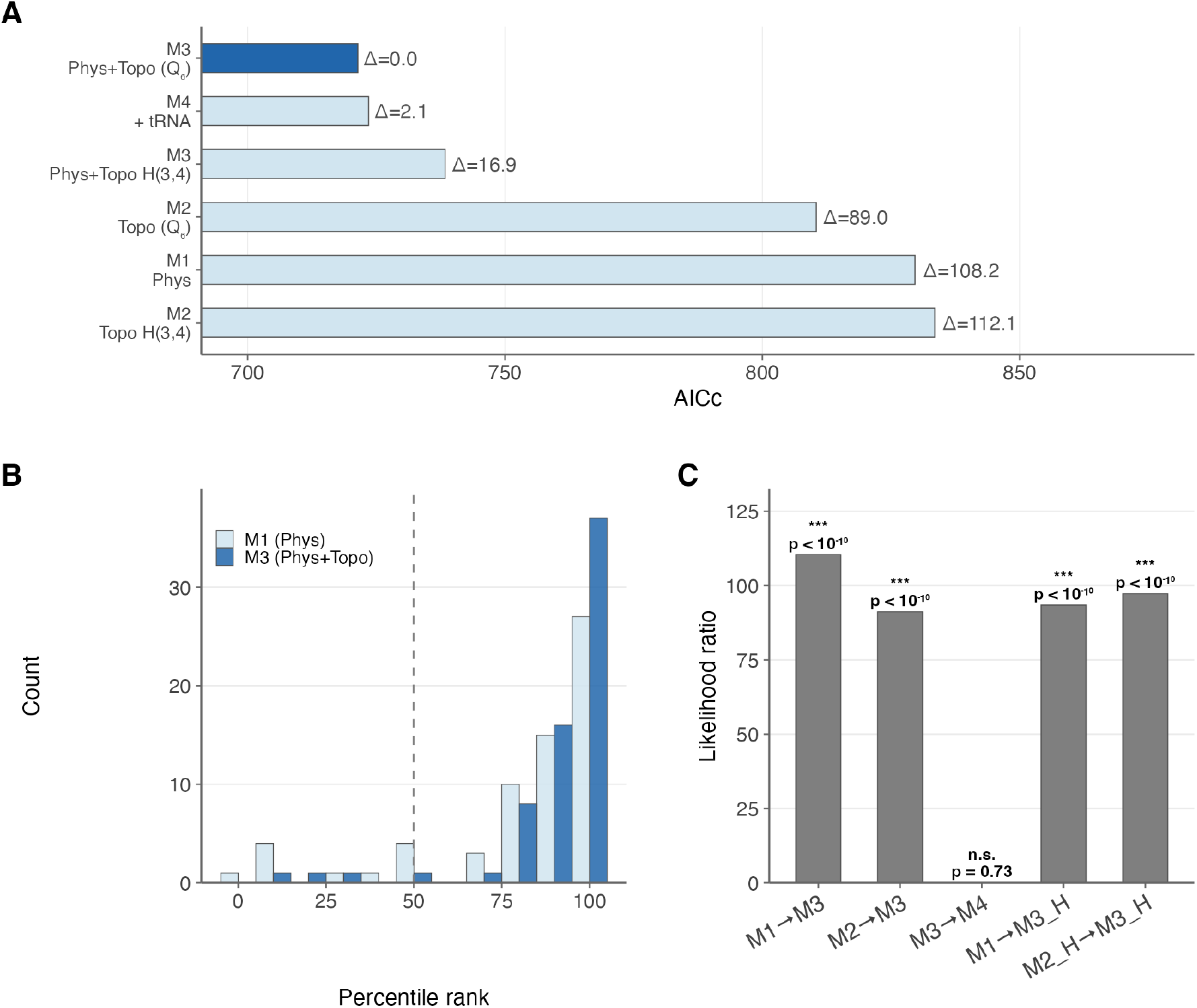
Event-level explanatory modeling of natural codon reassignments. **(A)** AICc comparison of the six candidate conditional-logit models (M1–M4 under *Q*_6_ plus the two *H*(3, 4)-verification variants; likelihood-ratio tests were applied only to nested pairs); lower is better. The combined physicochemistry-plus-topology model M3 has the lowest AICc; the single-feature baselines M1 and M2 are 89.1–112.1 AICc units worse, the *H*(3, 4)-topology verification model M3_H(3,4)_ is 16.9 units worse than M3_Q_6_, and M4 (adding the heuristic tRNA-distance proxy) is 2.1 units worse than M3. **(B)** Distribution of observed move percentile ranks under M1 (physicochemistry only, light) versus M3 (combined, dark); M3 concentrates ranks toward the top of the candidate set. Dashed line = chance expectation (50th percentile). **(C)** Likelihood-ratio tests for each feature class added to its complement; both topology and physicochemistry contribute highly significant independent information (*p* < 0.001); the heuristic tRNA proxy does not (*p* = 0.73).

**Figure 5:**
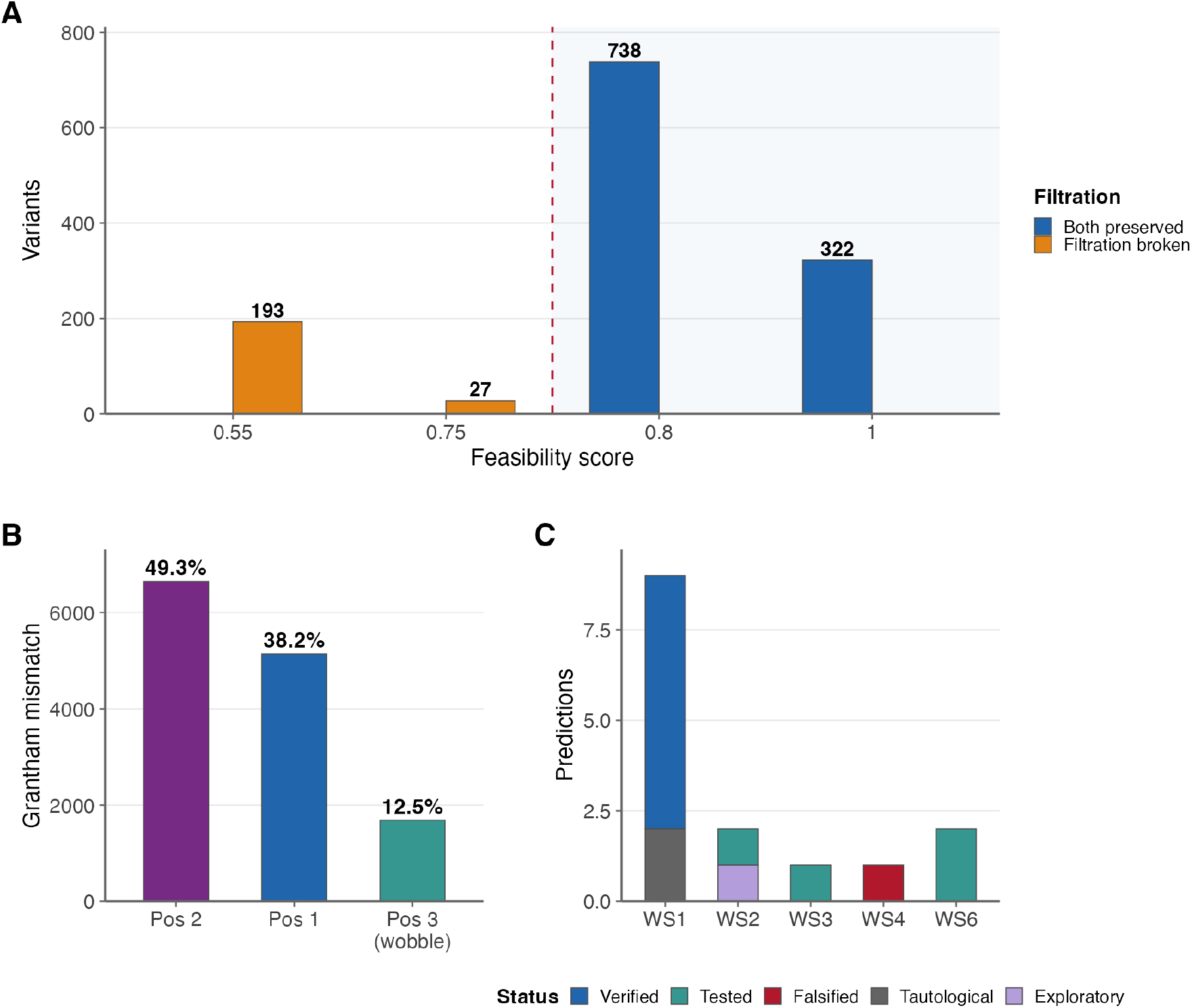
Translational applications and prediction catalogue. **(A)** Structural-preservation index (Supplement §S12) for 1,280 single-codon reassignments from the standard code, colored by two-fold/ four-fold filtration status. The index is a hand-weighted discrete composite of four indicators and takes only four values across the candidate universe (*S* ∈ {0.55, 0.75, 0.80, 1.00}); it is a visualization heuristic, not an inferential test. **(B)** Grantham mismatch score decomposition by nucleotide position: position 2 dominates (49.3%), consistent with the biochemical hierarchy of mutational impact. **(C)** Prediction catalogue: 15 evaluated claims grouped by workstream (WS1–WS4 plus WS6; WS5 is a synthesis workstream that generates the catalogue itself and has no independent claims), colored by the five registered claim-hierarchy categories: Supported, Rejected, Exploratory, Tautological, and Falsified (the KRAS–Fano decision gate). The per-prediction counts shown here (7 Supported / 4 Rejected / 1 Exploratory / 2 Tautological / 1 Falsified) sum to 15 and are the per-prediction workstream tally; the per-claim tally in the registered claim hierarchy (Table S1) is 4 Supported / 5 Exploratory / 3 Rejected / 1 Falsified / 2 Tautological, differing because multiple predictions can map to one claim.

Like all conditional logit models, M1–M4 assume Independence of Irrelevant Alternatives (IIA): the relative probability between any two candidate moves is unaffected by adding or removing other candidates. We use the model as an explanatory rather than predictive tool: the reported ΔAICc and likelihood-ratio statistics test whether topology adds explanatory value beyond physicochemical cost, not whether the model accurately predicts which specific reassignment will occur next; full IIA discussion appears in Supplement §S6. A separate concern is that the unrestricted candidate set (≈1,280 single-codon moves) admits strongly deleterious alternatives (reassigning AUG-Met, multi-codon changes implicit in single-step framing, reassignments to stop in essential codons) that selection has already removed. Models with strongly negative coefficients on Δ_topo_ and Δ_phys_ may therefore partly rediscover that natural reassignments are not catastrophic, inflating apparent ΔAICc gaps. To bound this concern we refit M1–M4 (and the *H*(3, 4) verification variants) on a **restricted candidate set** containing only candidates whose target amino acid is already accessible at Hamming distance ≤ *d* from the reassigned codon (Δ_tRNA_ ≤ *d*, with *d* ∈ {1, 2, 3} reported as a **bracketing** sensitivity rather than a designated primary; Supplement §S6.1). The strictest cut, *d* = 1, is the most defensible biological-plausibility floor (canonical wobble tolerance); *d* = 3 is a loose upper bound. Under *d* = 2 the candidate set shrinks from ≈1,280 to ≈ 727 candidates per choice set, and M3 still dominates with ΔAICc(M1 → M3) = 60 and ΔAICc(M2 → M3) = 77, both well above the conventional ΔAICc>10 reference. The qualitative explanatory claim survives even the most stringent *d* = 1 filter (≈ 275 candidates per choice set), where ΔAICc(M1 → M3) = 13.8 shrinks but remains above the conventional 10 threshold and ΔAICc(M2 → M3) = 73 remains large; under the looser *d* = 3 filter (≈ 1088 candidates), the gap recovers to ΔAICc(M1 → M3) = 95. As expected, tighter cuts shrink the ΔAICc gaps, but the qualitative claim “topology adds explanatory value beyond physicochemistry” is robust at every threshold. We therefore read the unrestricted-set ΔAICc magnitudes as **upper bounds** on the topology–physicochemistry separation and treat the *d* = 1 floor as the most defensible effect-size bracket. (We do not interpret ΔAICc(M3 → M4) from the restricted set: because the restriction is itself defined on Δ_tRNA_, the M4 tRNA-feature distribution shifts mechanically with the filter, so the M4 comparison is informative only under the unrestricted set, where it remains uninformative.)

Conditional-logit clade-exclusion sensitivity (refitting M1–M4 with each major clade dropped, matching the regime applied to the topology-avoidance hypergeometric in Sengupta et al. (2007)) is reported in Supplement §S7. Across all seven exclusion regimes, M3 is consistently favored over M1 (ΔAICc ≥ 49, median 101, max 117) and over M2 (ΔAICc ≥ 32, median 88). Even the minimum ΔAICc(M1→M3) of 49 (from excluding all metazoan mitochondrial codes, leaving 40 events) sits comfortably above the conventional ΔAICc>10 reference threshold; the qualitative conclusion (that topology adds explanatory value beyond physicochemistry) survives every clade-exclusion regime. We rely most heavily on the parametric predictive calibration check reported above (*p* = 0.60), since it does not require importing a threshold from a different statistical regime.

### 3.6 tRNA enrichment for reassigned amino acids

Organisms with variant genetic codes tend to show elevated tRNA gene copy numbers for the reassigned amino acid relative to standard-code controls. Across 24 variant–control pairings derived from 24 organisms with mixed provenance (15 of 18 tRNAscan-SE–verified genomes plus 9 organism-slots from literature/GtRNAdb/annotation; the three tRNAscan-verified genomes not in this analysis are boundary-case mechanisms discussed in §4.3; full 24-row catalogue with per-pair 2×2 counts in Supplement §S10.1), Fisher’s exact test combined via Stouffer’s *Z* method yields *p* = 1.31 × 10^−7^ (*Z* = 5.15) for the all-pairings analysis; this pooled value does not adjust for shared control organisms. To address the non-independence directly, we enumerated all maximal independent sets (MIS) of the conflict graph via Bron–Kerbosch on the complement, obtaining 332 MIS of size 6. Across the 332 sets the Stouffer combined *p*-value has median 0.037, best 0.012, and worst 0.104 264 of 332 (79.5%) sets fall below the 0.05 threshold and 0 of 332 below 0.01 (Table 7). A pre-specified topology-breaking subset (n=4: yeast mitochondrial Thr, *Scenedesmus* mitochondrial Leu, *Pachysolen* nuclear Ala, *Candida* nuclear Ser) is null (Stouffer *p* = 0.387), so the direct mechanistic link between tRNA-gene duplication and codon-family disconnection is not supported by the topology-breaking subset in isolation, and the elevated-tRNA pattern reflects a heterogeneous accommodation-of-decoding signal across variant-code lineages rather than a specific compensation-for-disconnection mechanism. The pattern is also not universal at the mechanistic level: *Blastocrithidia nonstop* (NCBI translation table 31) achieved UGA→Trp via anticodon stem shortening rather than gene duplication (Kachale et al., 2023), and *Mycoplasmoides* species with UGA→Trp use a single tRNA-Trp with anticodon modification. These boundary cases motivate the genome-size-dependent mechanistic landscape discussed in Section 4.3; the full 18-organism tRNAscan catalogue (including the three boundary-case genomes not in the pairings) and per-lineage mechanistic detail (*Tetrahymena thermophila* with 54 Gln tRNAs including 39 first-pass suppressors; six *Euplotes* species carrying TCA-anticodon tRNA-Cys reading UGA; and the *Blastocrithidia*/*Mycoplasmoides* boundary cases) are given in Supplement §S10, with the 24-row pairing/provenance table in §S10.1.

**Table 7:** Summary of tRNA enrichment tests across 24 pairings among 24 organisms with mixed provenance (15 of 18 tRNAscan-SE-verified genomes + 9 organism-slots from literature/GtRNAdb/ annotation; three tRNAscan-verified genomes not in this analysis — *Blastocrithidia nonstop, Mycoplas-moides genitalium, Mycoplasmoides pneumoniae* — are boundary-case mechanisms discussed in §4.3). MIS = maximal independent set, enumerated via the Bron–Kerbosch algorithm applied to the complement of the conflict graph (edges = shared organisms). The 24-pairing graph admits 332 MIS, all of size 6. Across the 332 sets the Stouffer combined *p*-value distribution has median 0.037, best 0.012, worst 0.104 264 of 332 (79.5%) fall below the 0.05 threshold and 0 of 332 below 0.01. The topology-breaking subset restricts to the four pairings whose underlying reassignment creates a new amino-acid disconnection under *Q*_6_ (yeast mitochondrial Thr, *Scenedesmus* mitochondrial Leu, *Pachysolen* nuclear Ala, *Candida* nuclear Ser); this subset yields Stouffer *p* = 0.387 and is null, so the tRNA-duplication mechanism is not directly evidenced by the topology-breaking cases in isolation. The all-pairings (*n* = 24) Stouffer *p* is not adjusted for shared control organisms and is retained as the pooled point estimate only.

| Test | *n* | Statistic | *p*-value |
| --- | --- | --- | --- |
| Fisher+Stouffer (all pairings) | 24 | $Z = 5.15$ | $1.31 \times 10^{-7}$ |
| MIS median (across all 332 MIS) | 6 | $Z \approx 1.79$ | 0.037 |
| **MIS worst-case** | **6** | **$Z \approx 1.26$** | **0.104** |
| MIS best-case | 6 | $Z \approx 2.24$ | 0.012 |
| Topology-breaking subset | 4 | $Z \approx 0.29$ | 0.387 |

### 3.7 Exploratory observations

Three exploratory observations are recorded here without inferential weight (full detail in Supplement §S23): (i) an apparent bit-position bias in codon reassignments (*χ*^2^*p* = 0.006 under a uniform null) attenuates after de-duplication (*p* = 0.075) and vanishes under codon-preserving permutation (*p* = 1.0), indicating that the signal reflects which codons are recurrently reassigned rather than a positional preference; (ii) a systematic catalogue across all 27 NCBI translation tables identifies four variant-code amino-acid disconnections at *ε* = 1 beyond the universal Serine disconnection: Thr in yeast mitochondrial (table 3), Leu in chlorophycean mitochondrial (tables 16, 22), Ala in *Pachysolen tannophilus* (table 26), and a tripartite Ser in *Candida*-clade nuclear (table 12); and (iii) Serine has the most extreme Atchley Factor 3 score (*F*_3_ = −4.760; Atchley et al. (2005)) and is uniquely disconnected under every encoding, but these two views of Serine’s anomaly are not fully independent, since both reflect its disproportionate codon diversity relative to its physicochemical footprint.

### 3.8 Retrospective cross-study reanalysis of synthetic genome recoding outcomes

To assess whether the structural properties identified in Sections 3.1–3.5 translate to measurable phenotypic consequences in synthetic biology, we performed a retrospective cross-study reanalysis of nine published genome recoding datasets, with quantitative analysis of eight (>217,000 codon-level observations; Table 8). Each codon substitution was classified by its GF(2)6 topology: whether it crosses a connected-component boundary at *ε* = 1, the change in local physicochemical mismatch cost (Δ*F*_local_, Grantham distance), and the Hamming distance between source and target vectors.

**Table 8:** Published genome recoding datasets analyzed. “Boundary?” indicates whether the synonymous codon swaps cross a connected-component boundary at *ε* = 1 in GF(2)6. ^†^bioRxiv preprint; not peer-reviewed at time of writing. Among quantitatively analyzed datasets, Syn57 provides a **topology-fixed exposure** (every Ser swap crosses the UCN↔AGY boundary; no Ala swap does) paired with a null gene-level transcriptomic test; the codon-level Δ*F*_local_ contrast within Syn57 is design-confounded and reported descriptively (see §3.8.1.2). Ding et al. (2024) is included qualitatively for cross-kingdom scope without quantitative extraction. The Syn57 row total of 60,240 codon-level observations comprises 37,146 serine recodings + 22,859 alanine recodings + 235 stop-codon recodings.

| Study | Swap type | $n$ | Boundary? | Key result |
| --- | --- | --- | --- | --- |
| Robertson et al. (2025) Syn57 | 4 Ser + 2 Ala + Stop | 60,240 | Ser=100%, Ala=0% | Topology-fixed exposure (gene-level null) |
| Fredens et al. (2019) Syn61 | Ser (TCG/TCA → AGC/AGT) | 18,218 | All cross | Informative null |
| Ostrov et al. (2016) | 7 codon types | 62,214 | Mixed | Segment viability |
| Napolitano et al. (2016) | Arg (AGR → CGN) | 12,888 | None | SRZ covariates |
| Grome et al. (2025) Ochre | Stop (TGA → TAA) | 1,195 | N/A | Growth data |
| Nyerges et al. (2024)<sup>†</sup> | 7 codon types | 62,007 | Mixed | Multi-omics |
| Lajoie et al. (2013) C321.ΔA | Stop (UAG → UAA) | 321 | N/A | Baseline |
| Frumkin et al. (2018) | Arg (CGU/CGC → CGG) | 60 | None | Translation efficiency |
| Ding et al. (2024) | Ser (TCG, mam-malian) | Variable | Yes | Cross-organism |

#### 3.8.1 A three-layer interpretation of recoding outcomes (exploratory)

This subsection is exploratory and hypothesis-generating. We propose a three-layer view of how codon-space structure relates to recoding outcomes, but the partition into evolutionary, recoding-burden, and design-deviation layers was **not** prespecified before the analyses below. The layering is therefore a working synthesis, not a tested model: each contrast below is individually informative, but their integration into a single three-layer schema is interpretive and would require an independent dataset built around a sharper hypothesis to confirm.

##### 3.8.1.1 Boundary crossing: validated but not predictive of transcriptomic perturbation

In the Robertson et al. (2025) Syn57 dataset (the only large-scale dataset with within-study variation in boundary crossing), all 37,146 serine recodings crossed the UCN↔AGY disconnection boundary at *ε* = 1, while all 22,859 alanine recodings remained within the compact GCN family. This provides an exact within-study topology contrast. However, genes containing exclusively serine-type versus exclusively alanine-type recodings did not differ in transcriptomic perturbation (Mann–Whitney *p* = 0.40, *n*_Ser_ = 129, *n*_Ala_ = 49), and the fraction of boundary-crossing recodings per gene was uncorrelated with | log_2_ FC| (*ρ* = 0.012, *p* = 0.46, *n* =3,510 genes). In a separate dataset, all 18,218 Syn61 serine recodings (Fredens et al., 2019) crossed the boundary; the seven design-to-final corrections that the Syn61 authors required for cell viability (Supplementary Data 19 of Fredens et al. (2019)) concentrated at specific functional units (a Ser70 site in *yaaY*, an intergenic separation between *ftsI* and *murE*, a Ser4 site in *map*, five positions in the *yceQ* pseudogene needed to preserve viable expression of the downstream essential gene *rne*) rather than at randomly distributed sites. Because all 18,218 forward swaps share the same two AA-identity-preserving swap subtypes (TCG→AGC and TCA→ AGT), |Δphys| at the AA-identity level is structurally constrained across all sites, and the 7 corrections do not lie at extreme |Δphys| or unusual Hamming-accessibility positions relative to the 18,211 non-correction sites. We therefore interpret the corrections as protein-level functional pressure at specific residues and operon-context constraints, not a global topology-driven burden, and treat this contrast as descriptive rather than as a hypothesis-test of GF(2)6 features. Together with the gene-level Syn57 null, this indicates that boundary crossing is a coarse geometric property of codon-family organization: validated as a structural distinction over evolutionary timescales but not predictive of acute transcriptomic perturbation in synthetic-biology engineering, where competing ecological pressures are absent and expression constructs are highly optimized.

##### 3.8.1.2 Local mismatch geometry: a fine-scale burden axis with positive signal

The second layer examines whether reassignment changes the physicochemical quality of the immediate mutational neighborhood. In the Syn57 contrast, serine swaps moved codons into better local Grantham neighborhoods (Δ*F*_local_ = −37.2 ± 30.5), whereas alanine swaps moved codons into worse neighborhoods at a fixed Δ*F*_local_ = +19.0 ± 0.0 (zero variance). This contrast is design-confounded and we report it descriptively rather than inferentially: every Syn57 alanine swap goes to the same target codon, so all alanine Δ_local_values are identical, and the Mann–Whitney *U* = 0, *p* < 10^−16^ comparison reflects design-level non-overlap rather than a within-group inferential test. The methodologically clean inferential test comes from a topology-fixed setting (arginine synonymous recodings (Napolitano et al., 2016), where all 12,888 AGR→CGN swaps remain within one connected component at *ε* = 1), in which local mismatch change correlated with established recoding-burden covariates: Δ*F*_local_ vs RBS deviation (*ρ* = −0.33, *p* < 10^−15^) and vs mRNA folding deviation (*ρ* = −0.12, *p* < 10^−15^). Among the four CGN targets, CGG has the lowest local mismatch cost (246 Grantham units) while CGC has the highest (421), and target-specific RBS deviation spans four orders of magnitude, suggesting that local mismatch and SRZ covariates capture partially independent dimensions of recoding burden.

##### 3.8.1.3 Design deviations: accessibility-dominated, not topology-directed

To test whether engineered systems escape design constraints along topology-defined low-burden directions, we compared the Syn57 design genome (Data file S2; (Robertson et al., 2025)) against the final verified genome (Data file S8) and identified 727 CDS-level codon differences. After filtering 7 genes with >10 deviations each (structural rearrangements, predominantly *bioF* with 311 changes), 326 genuine point deviations remained (267 synonymous, 59 nonsynonymous). Among the 163 deviations that changed local mismatch, 76 moved to a better neighborhood and 87 to a worse one, showing no directional bias (two-sided exact binomial *p* = 0.43, Clopper–Pearson 95% CI [0.39, 0.55]; Wilcoxon signed-rank *p* = 0.67). This null result was stable across all filtering thresholds tested (*p* > 0.5 at cutoffs 3, 5, 10, 15, 20, and 50 deviations per gene). By contrast, deviations strongly favored mutationally proximate moves: 83% were single-bit changes (Hamming distance 1 under the default encoding; note that Hamming distance is encoding-dependent, though nucleotide edit distance is invariant), and the mean Hamming distance was 1.21. Thus, realized deviations from the Syn57 design do not preferentially descend toward lower local mismatch; instead, they follow accessibility-dominated escape routes, favoring nearby codon states regardless of neighborhood quality. The Ding et al. (2024) mammalian TCG recoding independently confirms the universality of the serine disconnection across standard-code organisms, providing cross-kingdom validation.

##### 3.8.1.4 Segment-level recoding burden in the Ostrov 57-codon design

As an orthogonal test, we examined segment-level outcomes from the Ostrov et al. (2016) 57-codon genome design. Among 44 segments with fitness data (of 87 total), segments with more recoded codons in essential genes showed a suggestive correlation with worse doubling time (*ρ* = 0.34, raw *p* = 0.022, Bonferroni-corrected *p* = 0.066), while total recoding load showed no association (*ρ* = −0.10, *p* = 0.53). Problem segments (13 with lethal exceptions) trended toward higher essential-gene recoding load (73.5 vs 39.8 recoded sites, Mann–Whitney *p* = 0.086). Among 44 fully characterized segments, 17 showed spontaneous codon reversions, indicating ongoing design instability at positions where the engineered code was under selection pressure.

#### 3.8.2 Summary of cross-study reanalysis scope

Across the four synthetic-recoding contrasts, the results are consistent with (but do not prove) a three-layer view in which different structural descriptors may operate at different biological layers (family topology, local mismatch, and Hamming accessibility, respectively). Specific testable predictions follow: (i) a Syn-style recoding experiment that varies family-boundary crossing while holding local mismatch fixed should fail to predict transcriptomic burden (already partly observed in Syn57); (ii) a recoding scan that varies Δ*F*_local_ within a single boundary regime should predict RBS-context proxies (consistent with the Napolitano arginine result); (iii) directed evolution starting from a recoded design should produce escape mutations whose Hamming distribution is shaped more by local accessibility than by mismatch optimisation (consistent with the Syn57 design-deviation analysis). Each prediction admits a direct experimental test that would either confirm or falsify the layer structure proposed here.

### 3.9 Scope of the framework: rejected and null findings

Four conjectural extensions of the GF(2)^6^ representation were tested and cleanly separated from the surviving picture: (i) the KRAS–Fano clinical prediction was falsified against 1,670 MSK-IMPACT (Zehir et al., 2017) mutations (*p* = 1.0; Supplement §S13); (ii) the claim that Serine’s inter-family Hamming distance equals 4 under all 24 base-to-bit encodings is false (16 encodings yield distance 2 and only 8 yield distance 4), though Serine is disconnected at *ε* = 1 under every encoding (Supplement §S14); (iii) PSL(2,7) has no 64-dimensional irreducible linear representation and is therefore pre-rejected as a symmetry group under that specific realisation (Antoneli and Forger, 2011) (this narrower claim does not exclude other group actions on a 64-element set; Supplement §S15); and (iv) the coordinate-wise map GF(2)^6^ → ℂ^3^ sending base pairs to fourth roots of unity fails the required group-character identity *χ*(*x* + *x*) = *χ*(*x*)^2^ since *i*^2^ = −1 ≠ 1; the “holomorphic” qualifier adds no analytic content on a finite discrete domain (Supplement §S16). A separate null result on source-neighborhood burden (Mann–Whitney *U* = 301, *p* = 0.70; Supplement §S17) indicates that reassignment is not driven by local escape from costly source neighborhoods, and clarifies that the topology-avoidance constraint (§3.4) operates at the global graph-connectivity level rather than per-codon.

## 4 Discussion

### 4.1 An information-theoretic view of the genetic code

The central finding of this work is that the standard genetic code minimizes the physicochemical disruption caused by single-bit errors in GF(2)^6^ coordinates. This is not a new conclusion (Freeland and Hurst (1998) established error-minimization using nucleotide-level mutation models), but the hypercube representation makes the optimality principle geometrically explicit: the code is a good *coloring* of a structured graph, in the graph-theoretic sense that adjacent vertices (codons differing by one bit) tend to share labels (amino acids) or, when they differ, differ by small physicochemical distances.

The score decomposition (Figure 5, panel B) concentrates optimization at the second (49.3%) and first (38.2%) codon positions, with the wobble position contributing only 12.5% — an emergent property of the code’s structure rather than a model parameter. The *ρ*-robustness result (Figure 2, panel B) demonstrates that optimality is not an artifact of restricting attention to *Q*_6_: when the full *H*(3, 4) mutation graph is considered (*ρ* = 1), the signal strengthens. This error-minimization is complementary to, but distinct from, the finding of Itzkovitz and Alon (2007) that the code is also nearly optimal for carrying parallel regulatory information within protein-coding sequences.

The GF(2)^6^ representation contributes two analytical tools beyond a pure *H*(3, 4) analysis. First, the coordinatized decomposition of *H*(3, 4) into the 192 Hamming-1 edges of *Q*_6_ plus 96 within-nucleotide diagonal (Hamming-2) edges enables the weighted-mismatch score *F_ρ_* (Equation 2) to interpolate continuously from *ρ* = 0 (*Q*_6_ only) to *ρ* = 1 (full *H*(3, 4)); the monotonic strengthening of optimality as *ρ* → 1 (Table 3, §3.2) does not exist in a pure *H*(3, 4) analysis, where the diagonal partition is not coordinatized. Second, the Hamming-distance filtration [inine] provides a natural persistence parameter for the disconnection catalogue (Figure 1, panel A) and for the conditional-logit topology feature (Δ*β*_0_). The 24-encoding sensitivity audit is a necessary control rather than a contribution: it identified *Q*_6_ topology depletion as encoding-dependent (8 of 24 bijections give no signal; §3.4, Supplement §S4) and prompted promotion of *H*(3, 4) to primary, but does not extend the framework’s reach. Pure *H*(3, 4) has no encoding choice and needs no such audit.

The binary representation is therefore best understood as an analytical decomposition tool; its distinct contributions are the *ρ*-sweep continuum and the *ε*-filtration, with *ρ* a diagonal-edge weight rather than a transition/transversion weight (Methods §2.1, Supplement §S2).

### 4.2 Evolutionary preservation and topology avoidance

The per-table analysis (Figure 2) shows that 26 of 27 NCBI translation tables remain low-cost relative to their own quartet-shuffle ensembles. Because the per-table null lacks distance-matched draws (every null draw has *d_H_* ≥ 30 from the standard code, while every variant has *d_H_* ≤ 6; Supplement §S8), this result is **compatible** with preservation of error-minimization during reassignment but does not identify independent variant-specific optimization. The marginal exception is the most extensively reassigned code: translation table 3 (yeast mitochondrial, 6 codon changes) falls only slightly above the 5% threshold.

The topology avoidance result (Figure 3) provides a mechanistic complement: natural reassignments avoid creating new amino acid disconnections at a rate far below chance expectation. This depletion is not confined to the *Q*_6_ representation: under the full *H*(3, 4) single-nucleotide mutation graph, the observed proportion of topology-breaking reassignments remains 21.4% against a possible rate of 66.1% (RR = 0.32, 95% CI [0.16, 0.66], *p* ≤ 10^−4^). The signal magnitude is similar to the *Q*_6_ result (RR = 0.29); some *Q*_6_-disconnected pairs become connected when all single-nucleotide edges are admitted, slightly lowering the possible-rate denominator, but the qualitative conclusion survives: codon-family connectivity is functionally important under either the hypercube subgraph or the fuller single-substitution graph. The yeast mitochondrial threonine reassignment illustrates this cost: the CUN→Thr change required acquisition of a novel tRNA^Thr^ derived from tRNA^His^ via anticodon mutation (Su et al., 2011), creating the topology-breaking disconnection that makes translation table 3 the sole marginal outlier in the per-table optimality analysis.

Together these analyses indicate that code evolution is constrained along two partly independent axes, not one. Physicochemical smoothness preserves **protein function under mistranslation**: a reassignment that lands in a physicochemically similar neighborhood is one whose mistranslation errors remain tolerable. Topological integrity preserves **something different**: the connectivity of an amino-acid codon family determines whether existing decoding machinery (a tRNA serving wobble-related codons) can continue to service the family without new molecular hardware. The conditional logit analysis (Table 6) makes the separation explicit. The topology term retains major explanatory value (ΔAICc = 108 relative to a physicochemistry-only model) after accounting for local physico-chemical cost, while physicochemical cost retains comparable value (ΔAICc = 89) after accounting for topology; the two feature classes are only weakly correlated across the candidate landscape (*r_s_* = 0.15), so neither term is a redundant restatement of the other.

To rule out a coordinate artifact, the M3 refit with the encoding-independent *H*(3, 4) topology feature retains a similar explanatory value (ΔAICc(M1 → M3_H(3,4)_) = 91 vs. ΔAICc(M1 → M3_Q_6_) = 108; §3.5), confirming that the conditional-logit thesis does not depend on the choice of topology graph.

We do not interpret this as direct selection on topology as an abstract graph property; topology may still proxy aspects of decoding architecture (tRNA cross-recognition range, ribosomal A-site constraints) not explicitly modeled here. The central claim is structural: within the tested event landscape, topology avoidance cannot be dismissed as a by-product of physicochemical optimization or as a coordinate artifact of the binary representation. The contribution beyond single-axis treatments is the second axis itself: code evolution moves through a low-cost region **and** avoids fragmentation of the decoding substrate, with the two constraints partly orthogonal and neither reducible to the other.

### 4.3 Exploratory tRNA accommodation across variant-code systems

The tRNA enrichment result (Figure 3, panel D; Table 7) describes an **accommodation-of-decoding** pattern across variant-code lineages: variant-code genomes tend to carry elevated tRNA-gene counts for the reassigned amino acid relative to standard-code sisters (median MIS Stouffer *p* = 0.037 across 332 MIS of size 6). The signal is driven principally by topology-**preserving** stop-to-sense reassignments — UAR→Gln in ciliates, UGA→Cys in *Euplotes*, and UGA→Trp in *Blepharisma stoltei* — rather than by the four topology-**breaking** pairings, whose subset is null (Stouffer *p* = 0.387). *Blastocrithidia nonstop* and the two *Mycoplasmoides* species are excluded mechanistic boundary cases in the 24-pairing analysis (their reassignments proceed through single-tRNA anticodon-stem or base-modification changes without gene-copy expansion) and therefore do not enter the statistical signal. The result does not therefore establish compensation for codon-family graph disconnection; it is consistent with a broader “accommodation of decoding” whose distribution across lineages is heterogeneous. Case examples illustrate the variety of mechanisms in this heterogeneous set: *Tetrahymena thermophila* carries 54 Gln tRNAs including a large suppressor complement; *Blastocrithidia nonstop* achieves UGA→Trp via anticodon stem shortening rather than tRNA-gene duplication (Kachale et al., 2023); *Mycoplasmoides* species with UGA→Trp read both codons with a single tRNA-Trp through anticodon modification. The correlation between the mechanistic route (duplication vs anticodon modification vs base modification) and genome size that these three cases motivate is a **hypothesis** about how genome size might constrain the mechanistic repertoire, not a statistical association test in this manuscript.

### 4.4 Synthetic recoding outcomes: a three-layer interpretation

The retrospective cross-study reanalysis helps delimit which structural quantities in GF(2)6 are functionally relevant and at which biological layer. Boundary crossing is a valid codon-space distinction (serine and alanine provide a clean contrast in which one class always crosses a family boundary and the other never does), yet this distinction did not predict RNA-seq perturbation in Syn57. We therefore do not interpret codon-family boundary crossing as a general transcriptomic burden variable. Its relevance, if any, likely lies at a different biological layer, such as decoding architecture, translational fidelity, or evolutionary accessibility, rather than steady-state expression change.

By contrast, local mismatch geometry showed a clearer positive signal. In both the Syn57 contrast and the topology-fixed arginine setting, codon changes differed systematically in the quality of the physicochemical neighborhood they entered, and these differences tracked established recoding-burden covariates (Napolitano et al., 2016). This suggests that local mismatch is best understood as a fine-scale recoding-friction variable: not a universal master predictor, but a meaningful descriptor of neighborhood quality within an already specified codon-family structure.

The design-deviation analysis further clarifies scope. Genuine deviations from the Syn57 design were overwhelmingly short-range (83% single-bit in GF(2)6), with no directional bias in mismatch change. This indicates that realized escape is governed more by mutational accessibility than by systematic optimization of local neighborhood cost: once a design is under strain, the available exits appear to be chosen primarily by mutational proximity.

These observations are consistent with, but do not establish, a working hypothesis that “topology” should not be treated as a single explanatory variable. Under that hypothesis, different structural descriptors would matter at different biological layers: family topology for the geometry of reassignment space (Section 3.4), local mismatch for some aspects of recoding burden (Section 3.8.1.2), and Hamming accessibility for the short-range routes by which engineered designs deviate. Each of the three layer claims rests on a single retrospective contrast and the partition into layers was not prespecified, so the schema should be read as hypothesis-generating rather than as a tested model (see §3.8.2 for explicit predictions). That the Fredens et al. (2019) Syn61 design tolerated 18,218 boundary-crossing serine swaps as a class (a genome-wide engineering success rate, not a per-codon-position viability test), while the NCBI variant codes show 3.1-fold depletion of topology-breaking changes over evolutionary timescales, is consistent with this layered reading: boundary crossing may be more relevant to evolutionary trajectory constraints than to acute engineering costs, but a within-study experiment that varies family-boundary crossing while holding local mismatch fixed is required to test that prediction.

### 4.5 Broader connections

The GF(2)^6^ framework connects to three adjacent literatures that we treat only briefly here. A complementary line of BioSystems work treats the code algebraically using 2-adic dynamical systems on ℤ_2_ (Dragovich and Mišić, 2019, Khrennikov and Kozyrev, 2007; Yurova Axelsson and Khrennikov, 2024a, 2024b). The two descriptions use different geometries (ultrametric attractors vs. finite Hamming graphs) and we do not identify them. As a finite bridge, the Walsh–Hadamard / 2-adic spectral depth of the standard code (544, encoding-invariant across all 24 bijections) sits far below a block-size matched null mean of 689.3 ± 8.2 at *n* =2,000 (*z* = −17.74), but is mathematically constant under a stricter wobble-box-preserving label-permutation null; the Walsh signature is therefore an algebraic invariant of the wobble-box structure rather than a separate beyond-wobble optimality signal (Supplement §S21; the complementary wobble-free label-spectrum fraction *S* = 0.7514 is included as a second signature of the same structure).

A second line of work by Gonzalez et al. (2016) proposes a **non-power** binary representation of the code based on a redundant signed-binary numeration with Fibonacci-like positional weights [1, 1, 2, 4, 7, 8], in which codon-to-integer maps become intentionally many-to-one, mirroring code degeneracy. The 24 base-to-bit bijections we sweep are all **power** representations, in the sense that each codon’s 6-bit vector is the concatenation of three per-base 2-bit encodings; the non-power family is strictly larger and would give a different hypercube-like object with an edge structure not factorable across the three codon positions. Our primary topology-avoidance result is defined on the encoding-independent *H*(3, 4) mutation graph (nucleotide-level, invariant under any codon-level bijection whatsoever), so the paper’s principal claim is not exposed to this concern. The *Q*_6_ subgraph analysis, which is already encoding-dependent within the power family (§3.4, §S4), would benefit from a non-power extension; we flag this as a natural continuation but do not attempt the full sweep here.

Separately, Tsour et al. (2026)‘s deep proteomic survey of alternate RNA decoding in over 1,000 human samples identified 8,746 unique amino-acid substitutions; among the 5,611 unambiguous-pair events, 65.0% involve substitutions whose closest source–target codon pair differs at a single nucleotide, versus a baseline of 39.5%. The naive event-level binomial statistic ignores clustering of substitutions by substitution type, gene, sample, and measurement opportunity; the effect-size contrast (65.0% vs 39.5%) is the primary summary, and type-level analyses are given in Supplement §S22. This provides external empirical grounding for the codon-adjacency mechanism implicit in our edge-mismatch objective (Equation 1): the same single-nucleotide neighbourhood structure governs the observed alternate-translation events in mammalian proteomes, independent of the amino-acid distance metric.

### 4.6 Limitations

Several limitations should be noted.

### Encoding choice

The base-to-bit encoding is not unique. While coloring optimality holds across all 24 encodings, specific score values and rank orderings are encoding-dependent (Supplementary Material).

### tRNA enrichment fragility

The tRNA enrichment signal is present in the median-independent-subset sense (median Stouffer *p* = 0.037 across 332 MIS of size 6) but does not survive the strict worst-case subset criterion (worst-case *p* = 0.104), and the pre-specified topology-breaking subset (*n* = 4) is null (Stouffer *p* = 0.387). We therefore classify the result as exploratory rather than as demonstrating a universal or mechanism-specific compensation for codon-family disconnection. Per-genome tRNAscan-SE pseudogene fractions vary across the 18 verified assemblies and are reported in Supplement §S10 (`Pseudo` column of the tRNAscan-verified organisms table); we do not report a formal fragmentation-based sensitivity test in this manuscript.

### Conditional logit scope

The conditional logit model (Section 3.5) is event-level explanatory, not direct proof of biological causality. The candidate universe comprises all ≈1,280 single-codon reassignments regardless of mechanistic plausibility, and the tRNA-complexity proxy (Hamming distance to nearest target-AA codon) is heuristic. The non-significance of this particular proxy does not exclude that a richer model of tRNA repertoire change would contribute explanatory value.

### Survivorship bias

All event-level analyses operate on reassignments that *persisted* in extant lineages. Topology-breaking reassignments that produced fitness collapse and lineage extinction are unobservable. The depletion result is therefore consistent with both selection against attempting topology-breaking moves and selection against the lineages that attempted them; cross-sectional NCBI data cannot adjudicate. Both interpretations support the conclusion that codon-family connectivity constrains evolutionary trajectories, with different mechanistic implications for the locus of selection.

### Phylogenetic non-independence ceiling

The 28 de-duplicated reassignment events used in the topology-avoidance hypergeometric test cluster at the codon level (e.g., the recurrent UGA→Trp event is observed across many distantly related mitochondrial lineages and is collapsed to a single event for the de-duplicated test) but they are **not** independent at the level of evolutionary origin: mitochondrial codes share an ancestral mitochondrial-decoding regime, ciliate nuclear codes share a ciliate-specific eRF1 trajectory, and so on. A conservative lower bound on the number of phylogenetically independent topology-preservation origins is on the order of 4–6, well below the 22 non-breakers the de-duplicated count reports; the hypergeometric *p* should be read against that ceiling rather than against *n* = 22. The conditional-logit clade-exclusion sensitivity (Supplement §S7) and the within-clade enumerations in §3.4 jointly bound how strongly the depletion magnitude can be claimed.

Multiple-comparison correction was applied within analysis families rather than across all quantities; the eight test families address conceptually distinct, non-nested questions, and all headline results survive cross-family Bonferroni at 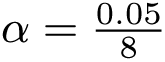 except the exploratory tRNA enrichment (detailed breakdown in Supplement §S18).

## 5 Conclusion

The standard genetic code is consistently low-cost: across four amino acid distance metrics and across a continuum of mutation graphs spanning *Q*_6_ and the complete single-nucleotide graph *H*(3, 4), it sits in the extreme tail of quartet-pattern shuffle null models (*p* ≤ 0.0062 under every combination). Across the 27 NCBI translation tables this near-optimality is preserved at the variant level: of the 12 codes whose distance from the standard makes the per-table test informative, 11 retain top-5% placement after BH–FDR correction, with only yeast mitochondrial (table 3) marginally above the 5% threshold; the 14 near-standard tables are formally significant in the same correction but we interpret these as primarily confirmatory of standard-code geometry. Code evolution therefore appears to occur within a constrained low-cost region rather than freely across code space.

Two partly independent constraints govern code evolution within that region. Persistent natural reassignments preferentially enter neighborhoods of lower local Grantham mismatch, and they avoid fragmenting amino-acid codon-family connectivity. Topology-breaking moves are approximately 3.1-fold depleted under the encoding-independent *H*(3, 4) adjacency, and conditional-logit decomposition shows that topology adds explanatory value beyond physicochemistry, and vice versa (ΔAICc ≥ 89 under *Q*_6_ topology; ΔAICc ≥ 91 under encoding-independent *H*(3, 4) topology). The two axes are only weakly correlated across the candidate landscape (*r_s_* = 0.15); neither is a redundant restatement of the other. The topology coefficient survives all seven phylogenetic clade exclusions (ΔAICc(M1 → M3) ≥ 49 in the worst regime), and a parametric predictive simulation under M3 reproduces the observed event-step topology-breaking rate under *Q*_6_ / Δ*β*_0_ > 0 (5/66 = 0.076 vs simulated mean 0.077; *p* = 0.60). Several variant-code lineages show exploratory tRNA-gene enrichment for the reassigned amino acid (median MIS Stouffer *p* = 0.037 across 24 pairings among 24 organisms of mixed provenance; worst-case *p* = 0.104; topology-breaking subset *p* = 0.39), consistent with heterogeneous accommodation-of-decoding rather than a specific compensation-for-disconnection mechanism. A retrospective cross-study reanalysis of eight published genome-recoding experiments (>217,000 codon-level observations) sharpens the scope of the topology constraint. Family-boundary crossing is a real structural distinction over evolutionary timescales but did **not** predict acute transcriptomic perturbation in the Syn57 gene-level contrast, while Syn61 had earlier tolerated 18,218 boundary-crossing serine swaps as a class (Fredens et al., 2019) (a genome-wide success rate, not a per-codon-position viability measure). This pattern is consistent with topology acting primarily as a constraint on evolutionary trajectories: gradual fragmentation of a codon family could create periods of decoding fragility that are bypassed in synthetic systems where decoding machinery is engineered up-front. Codon-space topology may therefore be a constraint on how genetic codes can change rather than on how they currently function: path-dependent rather than instantaneous. The three-layer integration of these descriptors (evolutionary topology, fine-scale recoding burden, accessibility-driven escape) is exploratory and hypothesis-generating: each layer-claim rests on a single retrospective contrast, the partition was not pre-specified, and confirmation requires the prospective tests outlined in §3.8.2.

The GF(2)^6^ representation is best understood as an analytical decomposition rather than as a claim about biologically privileged binary coordinates. Its main contributions are the *ρ*-sweep interpolating between *Q*_6_ and *H*(3, 4), the *ε*-filtration of codon-family connectivity, and the encoding-sensitivity audit required for representation-dependent analyses. The KRAS–Fano clinical prediction, Serine distance-4 invariant, PSL(2,7) symmetry, and holomorphic-embedding claims were tested and rejected (Table 1): several invariants of the representation turned out to be encoding artifacts. The surviving picture is encoding-independent: the genetic code is a low-cost coloring of *H*(3, 4), and observed reassignment trajectories preserve both physicochemical similarity and codon-family connectivity. Whether the topology axis reflects direct selection or a proxy for a specific feature of decoding architecture remains open. The present results support a working model in which natural genetic-code evolution is constrained by two partly independent axes: physicochemical smoothness and codon-family connectivity.

## Acknowledgements

We thank the NCBI, GtRNAdb, and cBioPortal teams for maintaining public databases essential to this work. tRNAscan-SE 2.0.12 was developed by Chan and Lowe at UC Santa Cruz.

## CRediT author contribution statement

**Paul Clayworth:** Conceptualization, Methodology, Formal analysis, Writing – original draft, Writing – review and editing. **Sergey Kornilov:** Conceptualization, Methodology, Software, Formal analysis, Data curation, Visualization, Validation, Writing – original draft, Writing – review and editing.

## Declaration of generative AI and AI-assisted technologies

During the preparation of this work the authors used GPT-5.2-Pro, GPT-5.5-Pro, Claude Opus 4.5, Claude Opus 4.7, GLM-5.1, Kimi-K2.5, MiniMax-M2.7, and Gemini 3 Pro to scaffold and review code, review the manuscript outline, and assist in drafting and revising the manuscript text. After using these tools, the authors reviewed and edited the content as needed and take full responsibility for the content of the publication. No generative AI or AI-assisted tools were used to create or alter figures or images in this manuscript.

## Ethical statement

This study did not involve human subjects, animal experiments, or clinical samples. All analyses were performed on publicly available data: NCBI genome assemblies, NCBI gc.prt translation table definitions, GtRNAdb tRNA gene catalogues, the MSK-IMPACT mutation registry as released through cBioPortal, and previously published genome-recoding datasets. No new biological materials were generated.

## Data and code availability

All code, raw data, and intermediate analysis outputs are publicly released in the `codontopo` repository at https://github.com/biostochastics/codontopo (version 0.6.1, tag `v0.6.1`). Every displayed number, table, and figure is reproducible from a clean clone by one canonical command:

`git clone` https://github.com/biostochastics/codontopo && `cd codontopo`

`git checkout v0.6.1`

`pip install-e “.[dev]”`

`bash scripts/build_publisher_release.sh`

The wrapper `scripts/build_publisher_release.sh` runs the analysis pipeline (`codon-topo all -- seed=135325 --n=10000`), the provenance-emit and table-generation scripts, both R figure scripts, and the two Typst PDF compilations, in that order. All inline statistics, tables, and figures in this manuscript and supplement are rendered directly from the pipeline outputs by the Typst sources (`manuscript.typ`, `supplement.typ`) included in the same repository. NCBI genome assembly accessions are listed in Supplement §S10.

## Footnotes

1 Code-dependent (MSA-derived), reported as a structure-aware robustness check; see Section 2.2 and Di Giulio (2001).

