## Supplementary Materials for "Robust error-minimization in the genetic code across physicochemical metrics and variant codes: a graph-theoretic analysis in GF(2)^6^"

#### Contents

S1 Claim hierarchy with full justifications ... S2
S2 Encoding sensitivity analysis (coloring optimality) ... S5
S2.1 Null-model choice: quartet-pattern vs classical Haig-Hurst AA-permutation ... S5
S3 Topology-breaking definitions: 2 × 2 audit ... S8
S4 Topology avoidance: $Q_6$ encoding-sweep sensitivity ... S10
S5 Reassignment candidate-universe denominator sensitivity ... S10
S6 Conditional logit: IIA assumption and explanatory framing ... S12
S6.1 Candidate-set composition: restricted-candidate sensitivity ... S13
S7 Conditional logit: clade-exclusion sensitivity ... S14
S8 Per-table optimality: standard-code-proximity audit ... S15
S9 Topology avoidance: clade-exclusion sensitivity (hypergeometric/permutation) ... S17
S10 Complete tRNA gene count data ... S19
S10.1 24-pairing input table with provenance and Fisher $2 \times 2$ counts ... S19
S10.2 tRNAscan-SE verified organisms ... S22
S10.3 MIS (maximal independent set) analysis ... S23
S10.4 *Saccharomyces cerevisiae* literature-derived control ... S24
S11 Complete reassignment database ... S24
S12 Structural-preservation index (visualization-only) ... S26
S13 KRAS–Fano clinical prediction: detailed results ... S26
S14 Serine minimum inter-family distance across encodings ... S27
S15 PSL(2,7) symmetry: pre-rejection by irrep dimensions ... S27
S16 Holomorphic embedding: character-identity failure ... S28
S17 Source-neighborhood burden: null result ... S28
S18 Cross-family multiple-comparison correction ... S28
S19 Event-level conditional logit model: full details ... S29
S19.1 Model specification and candidate universe ... S29
S19.2 Feature definitions ... S29
S19.3 Fitted coefficients ... S30
S19.4 Likelihood ratio tests ... S30
S19.5 Confounding diagnostic ... S31
S19.6 Observed move percentile ranks ... S31
S19.7 Order-averaging implementation ... S31
S20 ProtSub matrix and metric correlation analysis ... S32
S20.1 ProtSub as a code-dependent robustness check ... S32
S20.2 Metric correlation matrices ... S32
S21 Walsh–Hadamard / 2-adic spectral probe ... S33
S21.1 Block-size null ... S34
S21.2 Wobble-box-preserving label-permutation null ... S34
S21.3 Encoding invariance ... S34
S21.4 A second Walsh invariant: the wobble-free label spectrum ... S34

S22 Slavov (Tsour et al. 2026) SAAP cross-analysis ........ S36
S23 Exploratory observations: full detail ........ S38
S23.1 Bit-position bias in codon reassignments ........ S38
S23.2 Variant-code disconnection catalogue ........ S38
S23.3 Atchley Factor 3 and Serine convergence ........ S39
S23.4 Three-tier tRNA mechanistic landscape (from main-text §3.6) ........ S39
S24 Software and reproducibility ........ S40
References ........ S41

**Roadmap.** This supplement is organised around three roles. Sections §S1–§S5 give the **evidentiary scaffolding** underpinning every main-text claim: a registered hierarchy of all 15 claims with status and justification (Section 1), and the four sensitivity analyses that probe whether each headline result depends on a definitional choice: the 24-encoding sweep for cross-metric coloring optimality (Section 2), the $2 \times 2$ adjacency $\times$ topology-breaking-definition audit (Section 3), the per-encoding $Q_6$ topology-avoidance sweep (Section 4), and the four candidate-universe denominators (Section 5). Sections §S6–§S9 give the **robustness checks for the conditional-logit and topology-avoidance tests**: the IIA discussion and restricted-candidate sensitivity (Section 6, Section 6.1), conditional-logit clade exclusion (Section 7), the standard-code-proximity audit for the per-table optimality test (Section 8), and the hypergeometric/permutation clade-exclusion counterpart (Section 9). Sections §S10–§S19 give the **complete experimental and computational record**: the tRNAscan-SE 2.0.12 dataset on all 18 verified genomes (Section 10), the per-table reassignment database (Section 11), the structural-preservation index (visualization-only; Section 12), the per-variant KRAS–Fano falsification (Section 13), and the full event-level conditional-logit specification with fitted coefficients and likelihood-ratio tests (Section 19). Sections §S20–§S22 give the **additional bridging analyses**: the ProtSub matrix and metric correlation analysis (Section 20), the Walsh–Hadamard / 2-adic spectral probe (Section 21), and the Slavov (Tsour et al. 2026) SAAP cross-analysis (Section 22). Sections §S23–§S24 close with the full exploratory-observations catalogue (Section 23) and software/version/reproducibility metadata (Section 24). All numbers reported here are rendered from the same `manuscript_stats.json` and per-analysis JSON artifacts that drive the main text, so within a single pipeline run the two documents render from shared versioned artifacts; CI checks compare rendered values against those artifacts to reduce drift.

### S1 Claim hierarchy with full justifications

The 15 evaluated claims are organised into five evidentiary tiers (Supported / Exploratory / Falsified / Rejected / Tautological) and listed with full justifications in Table S1. The hierarchy is registered in code (`src/codon_topo/reports/claim_hierarchy.py` in https://github.com/biostochastics/codontopo) so that every analysis run produces a verifiable status report (`codon-topo claims`) rather than the status table being maintained as prose. The "Falsified" and "Rejected" rows record claims this paper or earlier work has tested and found to be wrong; we list them so that the framework's negative scope is visible alongside its positive results.

| Claim ID and statement | Status | Justification |
|---|---|---|
| **hypercube_coloring_optimality** The standard genetic code is significantly error-minimizing under four established, code-independent physicochemical distance measures with partially overlapping content: | Supported | Decision rule: all four code-independent metrics below the within-family Bonferroni threshold $\frac{\alpha}{4} = 0.0125$ under the quartet-pattern shuffle null ($n$ =10,000). Result: Grantham $p = 0.0062$, Miyata $p < 0.001$, polar requirement $p = 0.003$, |

| Claim ID and statement | Status | Justification |
|---|---|---|
| Grantham ($p = 0.0062$), Miyata ($p < 0.001$), Woese polar requirement ($p = 0.003$), and Kyte–Doolittle hydropathy ($p = 0.001$). | | Kyte–Doolittle $p = 0.001$. Stop penalty sensitivity (0/150/215/300): immaterial. |
| **per_table_optimality_preservation** 26 of 27 NCBI translation tables remain in the top 5% of their own quartet-pattern shuffle null for Grantham edge-mismatch (BH–FDR corrected); only translation table 3 (yeast mito) exceeds the threshold. | Supported | Per-table quartet-pattern shuffle null applied to all 27 NCBI tables; significant fraction (BH–FDR $p < 0.05$) and per-table quantiles are reported in the main text and the per-table CSV `output/tables/T4_per_table_optimality.csv` (released in the `codontopo` repository). Translation table 3 (yeast mito) is the marginal exception. |
| **optimality_rho_robustness** Coloring optimality is robust across all diagonal-edge weights $\rho \in [0, 1]$, including the full Hamming graph $H(3, 4) = K_4 \Box K_4 \Box K_4$ of single-nucleotide substitutions ($p < 0.01$ at all $\rho$ values). | Supported | $\rho$-sweep at $n$ =10,000: $p \leq 0.0062$ at all $\rho$ values; effect-size $z$ increases monotonically from $\rho = 0$ to $\rho = 1$. |
| **topology_avoidance_depletion** Natural codon reassignments are depleted for topology-breaking changes (moves that fragment an amino acid's codon family) at approximately 21% of observed events vs 66–73% of the candidate landscape, robust to adjacency definition ($Q_6$ vs the full Hamming graph $H(3, 4)$) and clade exclusion. | Supported | Decision rule: hypergeometric $p < 10^{-3}$ under the primary $H(3, 4)/\Delta\beta_0 > 0$ specification, replicated under the $Q_6$/new-disconnection sensitivity. Result: permutation $p \leq 10^{-4}$ under both adjacencies; hypergeometric $p = 1.6 \times 10^{-8}$ ($Q_6$, new disconnection) and $p = 1.3 \times 10^{-6}$ ($H(3, 4)$, $\Delta\beta_0 > 0$). 5–7 of 28 de-duplicated events are topology-breaking versus $\approx$ 64–75% of the candidate landscape ($\approx$ 3.0–3.4-fold depletion). $Q_6$ is encoding-dependent (8 of 24 bijections give no depletion); $H(3, 4)$ is encoding-independent and is the primary test. Clade-exclusion sensitivity (7 regimes): all $p < 10^{-3}$ under $H(3, 4)/\Delta\beta_0 > 0$ (largest $p \approx 2.03 \times 10^{-4}$); all $p < 10^{-5}$ under $Q_6$/new-disconnection. |
| **trna_enrichment_reassigned_aa** Organisms with variant genetic codes tend to show elevated tRNA gene copy numbers for the reassigned amino acid, but the enrichment does not attain a strict worst-case-subset threshold and the topology-breaking-only subset is null. | Exploratory | 332 maximal independent sets of size 6 (Bron–Kerbosch on the complement of the conflict graph): median Stouffer $p = 0.037$, best $p = 0.012$, worst $p = 0.104$; 264/332 (79.5%) sets fall below 0.05. Topology-breaking subset ($n = 4$) Stouffer $p = 0.387$ (null). 18 tRNAscan-SE-verified assemblies (15 variant + 3 standard controls), 24 pairings across 5 variant codes. |
| **bit_position_bias_weighted** Codon reassignment bit-flip distribution shows positional skew in $GF(2)^6$ coordinates under a uniform null, but the signal does not survive nulls that respect the source-codon non-independence of recurrent reassignments. Reported as exploratory/null rather than a positive finding. | Exploratory | Uniform null $p = 0.006$ but inflated by recurrent-reassignment non-independence (e.g., the recurrent UGA→Trp event contributes the same source codon across many lineages). De-duplicating to unique (codon, target) pairs gives $p = 0.075$. The codon-preserving null (which permutes target amino acids while holding the source codon fixed and is the appropriate non-independence-respecting null for this question) absorbs the apparent |

| Claim ID and statement | Status | Justification |
|---|---|---|
| | | bias entirely. We retain the row in the registry as a cautionary record of how the choice of null can flip a putative positive into a null finding, and we treat the bit-position-bias claim as not supported. |
| **mechanism_boundary_conditions** tRNA gene duplication accompanies codon reassignment in large nuclear genomes (ciliates, yeasts) but not in streamlined genomes (*Blastocrithidia*: anticodon stem shortening; *Mycoplasmoides*: anticodon modification). | Exploratory | Three-tier pattern: duplication / stem shortening / modification. Descriptive. |
| **atchley_f3_serine_convergence** Serine's extreme Atchley Factor 3 score ($F_3 = -4.760$, most extreme of 20 amino acids, 2.24 SD below mean) converges with the $\mathrm{GF}(2)^6$ topological disconnection: $F_3$ captures the mismatch between Serine's small physicochemical footprint and its disproportionate codon diversity, which the geometric framework identifies as maximal inter-family Hamming distance among 6-codon amino acids (4 vs 1 for Leu and Arg). | Exploratory | Serine $F_3 = -4.760$, 2.24 SD below mean. Complementary, not independent. |
| **variant_code_disconnection_catalogue** Systematic survey of Hamming-graph connectivity across 27 NCBI translation tables identifies 4 lineage-collapsed variant-code amino-acid disconnections at $\varepsilon = 1$ under the default encoding: Thr in yeast mitochondrial code (table 3), Leu in chlorophycean mitochondrial codes (tables 16 and 22, collapsed to a single algal-mito event), Ala in *Pachysolen tannophilus* nuclear code (table 26), and tripartite Ser in *Candida*-clade alternative yeast nuclear code (table 12). | Exploratory | 4 variant-code cases at $\varepsilon = 1$ in $Q_6$ (default encoding): Thr (Table 3, yeast mito), Leu (Tables 16/22, chlorophycean and *S. obliquus* mito; collapsed to one algal-mito event), Ala (Table 26, *Pachysolen*), Ser (Table 12, *Candida*). Separately, Table 32 (Balanophoraceae plastid, UAG⟶Trp) creates a 2-fold Trp pair (UGG, UAG) that remains connected at ε=1 but breaks the bit-5 two-fold filtration (a filtration finding, not a disconnection). |
| **kras_fano_clinical_prediction** XOR "Fano" relationships in $\mathrm{GF}(2)^6$ predict enrichment of specific amino acids at KRAS G12 co-mutation sites. | Falsified | $p = 1.0$ across all 6 G12 variants. $n =$1,670 MSK-IMPACT mutations. |
| **serine_min_distance_4_invariant** Serine's minimum UCN–AGY Hamming distance = 4 holds invariantly across all 24 base-to-bit encodings. | Rejected | 16/24 encodings give distance 2. Only 8/24 give distance 4. |
| **psl_2_7_symmetry** $\mathrm{PSL}(2, 7)$ is the fundamental symmetry group of the genetic code. | Rejected | No 64-dim irrep. Antoneli and Forger (2011). |
| **holomorphic_embedding** The coordinate-wise map $\mathrm{GF}(2)^6 \to \mathbb{C}^3$ sending base-pairs to fourth roots of unity is | Rejected | Domain is finite discrete. Character identity fails: $i^2 = -1 \neq 1$. |

| Claim ID and statement | Status | Justification |
|---|---|---|
| a holomorphic embedding extending a character of GF(8)*. | | |
| **two_fold_bit_5_filtration** All 2-fold synonymous codon pairs in the standard code differ at exactly bit position 5 under the default encoding. | Tautological | Forced by encoding choice. Holds in 16/24 encodings. |
| **four_fold_prefix_filtration** All 4-fold synonymous codon groups share a 4-bit prefix with 2-bit suffixes exhausting $GF(2)^2$. | Tautological | Trivial under any bijection from 4 bases to $GF(2)^2$. |

Table S1: Complete claim hierarchy with full claim statements and justifications.

#### S2 Encoding sensitivity analysis (coloring optimality)

There are $4! = 24$ distinct bijections from $\{C, U, A, G\}$ to $GF(2)^2$ (24 ways to assign the four nucleotide letters to the four 2-bit binary patterns 00/01/10/11). The default encoding used in the main text is $C \mapsto 00$, $U \mapsto 01$, $A \mapsto 10$, $G \mapsto 11$ (rationale stated in main-text §2.1). To check that the coloring-optimality result is not specific to that choice, we re-ran the Grantham quartet-pattern shuffle null analysis under each of the 23 alternative encodings, holding everything else fixed (per-encoding $n$ =10,000 null draws, seed 135325). All 24 encodings yield significant optimality ($p < 0.05$ under the quartet-pattern shuffle null), with all per-encoding conservative $p$-values bracketed by 0.0002 (best) and 0.0278 (worst). The mean per-encoding quantile is 1.09% (range 0.01%–2.77%), confirming that the result is not an artifact of the default encoding.

Properties that are encoding-invariant:

- Serine disconnection at $\varepsilon = 1$ (holds under all 24 encodings)
- Coloring optimality significance (all 24 significant)
- Four-fold prefix filtration (tautological under any bijection)
- $H(3, 4)$ adjacency graph (encoding-independent by construction)

Properties that are encoding-dependent:

- Serine inter-family minimum Hamming distance (4 in 8/24 encodings, 2 in 16/24)
- Two-fold bit-5 filtration (holds in 16/24 encodings)
- $Q_6$ (Hamming-1) adjacency partition of $H(3, 4)$, and the topology-avoidance signal computed under $Q_6$ adjacency (see §S4)
- Specific score values and rank orderings

Full per-encoding results are written to `output/coloring_optimality.json` (block `encoding_sensitivity`) by the standard `codon-topo all` invocation, and reproducible from the `codontopo` repository.

##### S2.1 Null-model choice: quartet-pattern vs classical Haig-Hurst AA-permutation

The primary coloring-optimality tests throughout this paper use the **quartet-pattern shuffle** null (main-text §2.3.1: the 64 codons are grouped into 16 first-two-base quartets, and the amino-acid pattern at each

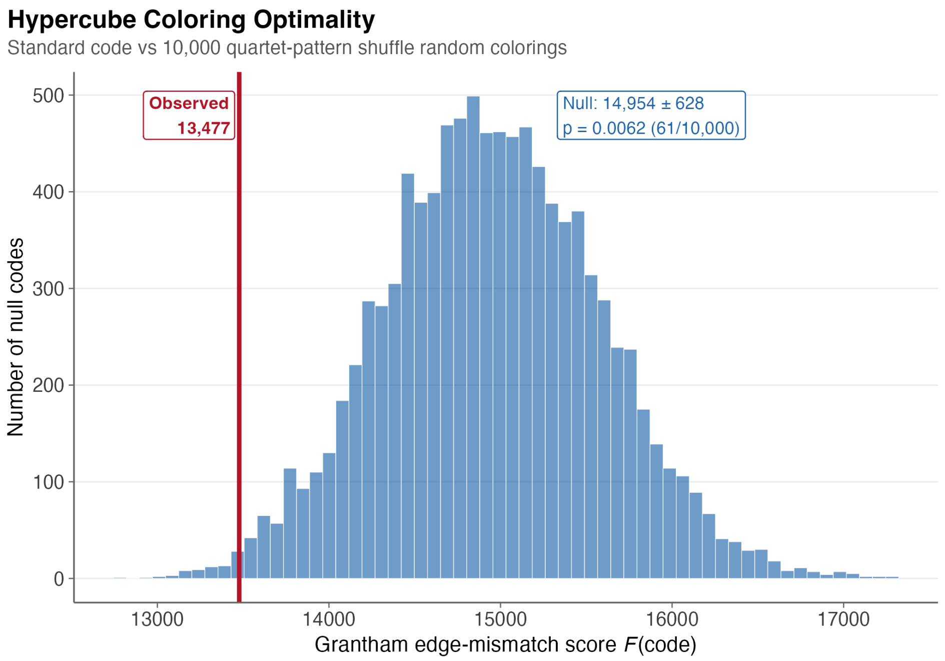


Figure S1: Hypercube coloring null distribution (extended). Histogram of Grantham edge-mismatch scores $F$ across the quartet-pattern shuffle null draws under the default encoding (C=00, U=01, A=10, G=11); the observed standard-code score is marked and its quantile shown. Companion to main-text Figure 2A, retained here at higher resolution together with the per-encoding sweep and per-table breakdown below.

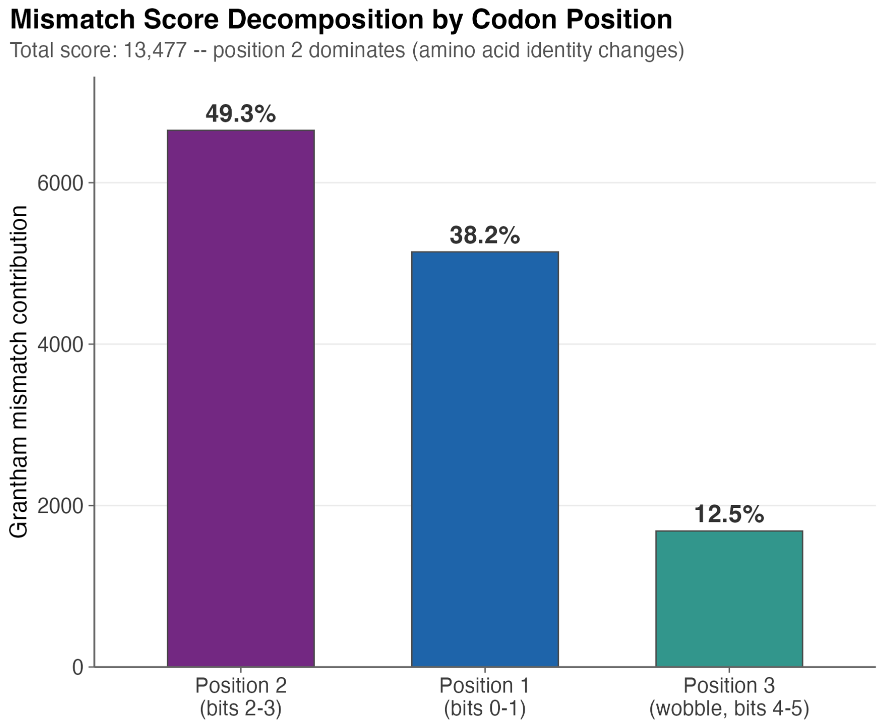


Figure S2: Grantham mismatch score decomposition by nucleotide position. Contribution of each codon position (1, 2, 3) to the total Grantham edge-mismatch score $F$ under the default encoding. Position 2 dominates, consistent with the biochemical hierarchy of mutational impact. This is a descriptive breakdown of the summed score, not an inferential test.

quartet is held as an atomic labeled 4-tuple; the two stop-containing quartets in the standard code (UA and UG) are held fixed and the remaining 14 patterns are permuted uniformly at random among the 14 non-stop quartet slots). This is distinct from the classical Haig–Hurst 1991 / Freeland–Hurst 1998 null commonly cited in the code-optimization literature, which fixes the standard code's 20 sense-codon families (the unlabeled partition of the 61 sense codons into synonymous groups) and permutes the 20 amino-acid labels uniformly across those families with stop positions held fixed. The two ensembles preserve different structural properties (Table S2) and admit different numbers of alternative codes ($\approx 14!$ for the quartet-pattern shuffle versus $\approx 20!$ for the AA-permutation null). The quartet-pattern-shuffle count is 14! rather than 16! because in the standard code both stop-containing quartets (UA and UG) are held fixed, leaving 14 mobile quartet slots.

| **Property** | **Quartet-pattern shuffle** | **Classical Haig–Hurst AA-permutation** |
|---|---|---|
| Stop-codon positions | fixed | fixed |
| Codon-family partition (which codons decode to a common amino acid, as an unlabeled clustering) | permuted | fixed |
| Per-named-amino-acid codon count (e.g., Leu's total codon count) | fixed | permuted |
| Sorted degeneracy multiset ($(6, 6, 6, 4, 4, ..., 1)$) | fixed | fixed |
| Quartet-slot AA-pattern shape at each first-two-base slot | permuted (atomic 4-tuples redistribute across slots) | fixed (labels within each slot swap only) |
| Approx. number of possible codes | $14! \approx 8.7 \times 10^{10}$ | $20! \approx 2.4 \times 10^{18}$ |
| Historical reference | not standard; introduced here | (Freeland and Hurst, 1998, Haig and Hurst, 1991) |

Table S2: **Preservation properties of the two null ensembles.** The two nulls preserve non-nested structural properties. The quartet-pattern shuffle preserves each named amino acid's total codon count (because the atomic 4-tuples that move carry their internal AA composition with them) but permutes which specific codons decode to a given amino acid; the classical Haig–Hurst AA-permutation preserves which codons decode together (the family partition) but allows a named amino acid's codon count to change (Trp's single-codon family may become a 6-codon family if its label is swapped with Leu's). Neither ensemble is uniformly "more stringent." The quartet-pattern-shuffle state space is 14! rather than 16! because in the standard code both stop-containing quartets (UA, containing UAA/UAG, and UG, containing UGA) are held fixed, leaving 14 mobile quartet slots.

We reran the four physicochemical metrics under the classical Haig–Hurst AA-permutation null ($n$ =10,000 draws, seed 135325) for direct comparison against the quartet-pattern shuffle values reported in main-text Table 2. Results are in Table S3.

| **Metric** | **Observed $F$** | **QP null $z$** | **QP $p$** | **HH-AA null $z$** | **HH-AA $p$** | **Δ (HH – QP)** |
|---|---|---|---|---|---|---|
| Grantham | 13477 | 2.35 | 0.0062 | 2.76 | 0.0021 | HH more extreme |
| Miyata | 235.3 | 3.14 | 0.0004 | 4.13 | 0.0001 | HH more extreme |
| Polar requirement | 347.3 | 2.73 | 0.0026 | 3.45 | 0.0001 | HH more extreme |
| Kyte–Doolittle | 457.2 | 2.78 | 0.0014 | 2.68 | 0.0027 | QP more extreme |

Table S3: **Coloring-optimality $p$-values under the quartet-pattern shuffle (QP, primary) vs the classical Haig–Hurst AA-permutation null (HH-AA, sensitivity).** Both use $n$ =10,000 draws, seed 135325. Conservative $p$-values are $(k+1)/(n+1)$ with the tail convention $f \leq F_{\text{obs}}$ (draws as extreme as or more extreme than the observed score). Treating the eight reported $p$-values as one test family, all eight pass Bonferroni at $\alpha = 0.05/8 = 0.00625$. For three of the four code-independent metrics the classical HH-AA null gives a **more** extreme $p$: because the HH-AA ensemble admits a wider space of alternative codes (20! vs 14!), the standard code sits further into its low-cost tail. Kyte–Doolittle inverts: hydropathy is unusually well-aligned with the wobble-quartet structure that QP preserves, so the QP null contains many low-hydropathy-mismatch alternatives that the HH-AA null (which breaks that alignment) does not, and the observed code's percentile is deeper against QP than against HH-AA. The conclusion — that the standard code is significantly error-minimizing across all four code-independent physicochemical metrics — is robust to the choice of null.

**Why we do not describe either null as "more stringent" than the other.** The two ensembles preserve non-nested structural properties (Table S2), and empirical tail behaviour (Table S3) shows the metric-specific direction of the $p$-value shift is not monotone across the four metrics. We therefore report both, note the null under which each headline $p$-value was computed (the QP shuffle), and treat the HH-AA null as a same-direction sensitivity check rather than as a validation or a refutation of the QP tail.

#### S3 Topology-breaking definitions: 2 × 2 audit

The topology-avoidance result depends on two independent definitional choices: (i) which adjacency graph defines codon-family connectivity ($Q_6$ Hamming-1 in the default $\mathrm{GF}(2)^6$ encoding, vs the encoding-independent $H(3,4)$ single-nucleotide graph), and (ii) what counts as a "topology-breaking" candidate move (a candidate move creates a **new** disconnection in a previously connected family, vs the candidate move strictly increases $\beta_0$ summed across families). To prevent silent dependence on either choice, we report all four combinations.

We define two notions of "topology-breaking" candidate move:

1. **New disconnection in a previously connected family**: a candidate move makes some amino acid disconnected at $\varepsilon = 1$ that was connected in the standard code (i.e., the amino acid was not previously in the disconnection catalogue but is after the move).
2. **Increase in components** ($\Delta\beta_0 > 0$, the conditional-logit feature): the total number of connected components, summed across amino-acid codon graphs, strictly increases:

$$\Delta_{\text{topo}} = \sum_a \beta_0\big(G_a^{\text{after}}\big) - \sum_a \beta_0\big(G_a^{\text{before}}\big) > 0.$$

Both definitions are reported under both $Q_6$ adjacency (Hamming-1 in the default $\mathrm{GF}(2)^6$ encoding) and $H(3,4)$ adjacency (full single-nucleotide adjacency, encoding-independent), giving four cells. All four share the same denominators (1,280 candidate moves, 28 de-duplicated observed events). The full $2 \times 2$ result is shown in Table S4.

| Adjacency | Topology-breaking definition | $K/N$ | $x/n$ | Hyper. $p$ | RR (95% CI) |
|---|---|---|---|---|---|
| $H(3,4)$ (primary) | $\Delta\beta_0 > 0$ (increase in components) | 846 / 1280 | 6 / 28 | $1.28 \times 10^{-6}$ | 0.32 (0.16–0.66) |
| $H(3,4)$ | new disconnection in connected family | 822 / 1280 | 5 / 28 | $4.95 \times 10^{-7}$ | 0.28 (0.13–0.62) |
| $Q_6$ | $\Delta\beta_0 > 0$ (increase in components) | 963 / 1280 | 7 / 28 | $2.23 \times 10^{-8}$ | 0.33 (0.17–0.63) |

| Adjacency | Topology-breaking definition | $K/N$ | $x/n$ | Hyper. $p$ | RR (95% CI) |
|---|---|---|---|---|---|
| $Q_6$ | new disconnection in connected family | 931 / 1280 | 6 / 28 | $1.58 \times 10^{-8}$ | 0.29 (0.14–0.6) |

Table S4: **Topology-avoidance $2 \times 2$ definition $\times$ adjacency audit.** All four cells share the same denominators (1280 candidate moves, 28 de-duplicated observed events) and differ only in (i) which adjacency graph defines codon-family connectivity ($Q_6$ Hamming-1 vs encoding-independent $H(3,4)$) and (ii) what counts as a topology-breaking move ($\Delta\beta_0 > 0$ summed over amino acids vs creation of a **new** disconnection in a previously connected family). $K/N$ = topology-breaking candidates among all candidates; $x/n$ = topology-breaking observed events among all observed events. All four cells yield depletion in the same direction with comparable risk ratios (0.28–0.33) and hypergeometric $p < 10^{-5}$. The main text uses the $H(3,4)$, $\Delta\beta_0 > 0$ cell as primary (encoding-independent adjacency, definition matched to the conditional-logit feature).

The full machine-readable audit is released as part of the `codontopo` repository's `output/` artifacts (the `definitions_audit` block in the topology-avoidance results file).

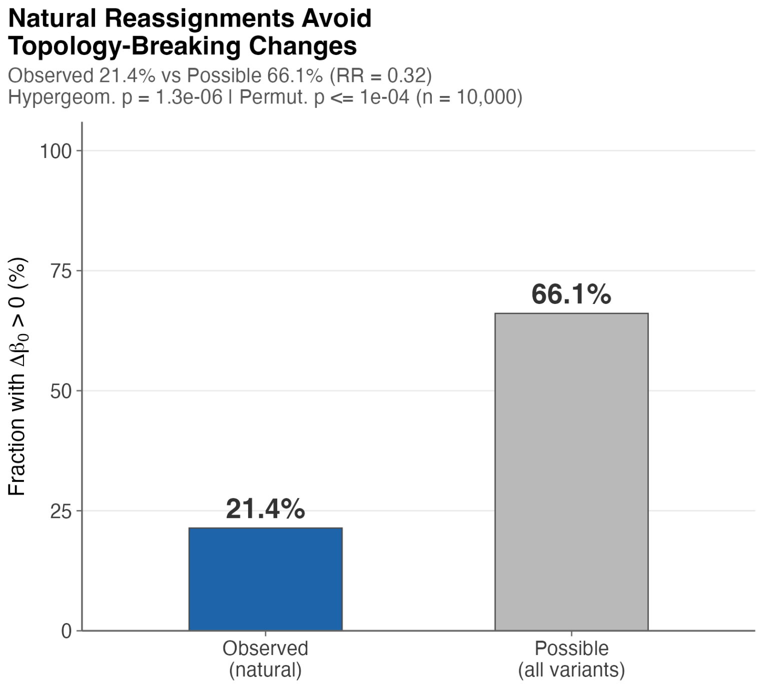


Figure S3: Topology-avoidance depletion under the primary $H(3,4)$ adjacency, $\Delta\beta_0 > 0$ definition (row 1 of Table S4). Observed rate of topology-breaking events (bar with permutation $p$-value annotation) contrasted with the candidate-landscape rate (dashed reference line). All four cells of the $2 \times 2$ audit above give qualitatively equivalent depletion (RR 0.28–0.33, hypergeometric $p < 10^{-5}$); this figure visualises the primary cell.

### S4 Topology avoidance: $Q_6$ encoding-sweep sensitivity

This section documents what motivated the demotion of $Q_6$ from primary to secondary topology adjacency in the manuscript.

The $H(3,4)$ Hamming graph is encoding-independent: every two-bit bijection from $\{A, C, G, U\}$ to $\{0,1\}^2$ produces the same nucleotide-level adjacency, because $H(3,4)$ depends only on which nucleotide letters differ between two codons, not on their binary encoding. The $Q_6$ subgraph, however, depends on the encoding because the partition into Hamming-1 (single-bit-change) edges versus Hamming-2 (within-nucleotide diagonal) edges is bijection-specific: under one encoding two codons that differ at exactly one nucleotide may project to a Hamming-1 edge of $Q_6$, and under another encoding they may project to a Hamming-2 edge. To test whether the $Q_6$ topology-avoidance result survives this representation choice, we recomputed the $Q_6$ candidate-landscape rate, observed rate, depletion fold, and hypergeometric $p$-value under all 24 base-to-bit bijections, holding the same 1,280 candidate moves and 28 observed events.

Across all 24 encodings:

- candidate-landscape rate: min 36%, median 69.9%, max 72.7%
- observed rate: min 21.4%, median 28.6%, max 35.7%
- depletion fold: min 1.01, median 2.45, max 3.39
- hypergeometric $p$: min $1.58 \times 10^{-8}$, median $6.08 \times 10^{-6}$, max 0.572

The default encoding (C=00, U=01, A=10, G=11) gives the largest depletion and the smallest $p$. Eight of 24 encodings give a candidate-landscape rate of approximately 36%, under which the observed rate of 21–36% does not significantly differ from candidate ($p > 0.5$). The $Q_6$ result is therefore not encoding-invariant. We accordingly present the encoding-independent $H(3,4)$ result as the primary topology-avoidance test, with $Q_6$ reported as a representation-specific decomposition for continuity with the broader $\mathrm{GF}(2)^6$ framework. The default encoding was adopted in companion methodological work (Clayworth (2026)) prior to the present encoding sweep, on the grounds of visualization clarity (it places the standard code's nine 2-fold-degenerate amino acids on bit-5 differences). We make no claim that this encoding is biologically privileged. The full per-encoding sweep is released as part of the `codontopo` repository's `output/` artifacts (the `Q6_encoding_sweep` block in the topology-avoidance results file); Figure S4 visualizes the per-encoding depletion fold and hypergeometric $p$.

### S5 Reassignment candidate-universe denominator sensitivity

The hypergeometric and permutation tests of topology avoidance, and the conditional-logit denominator, both depend on a definition of the candidate-move universe $\mathcal{M}(C)$ from the standard code $C$. Two definitional choices interact: whether stop-codon **targets** are admitted ($y \in \mathcal{A}_{20}$ vs $y \in \mathcal{A}_{20} \cup \{\text{Stop}\}$), and whether identity moves ($y = C(x)$) are excluded. The four combinations give the variants below; we adopt U1 as primary throughout and report U2 and U4 as sensitivity universes. All four universes admit stop **sources** (i.e. $x \in \mathcal{C}$ for all 64 codons); the earlier framing that referred to a source-stop axis was inaccurate and has been removed.

Formally, the primary candidate-universe is $\mathcal{M}(C) = \{(x, y) : x \in \mathcal{C}, y \in \mathcal{A}_{20} \cup \{\text{Stop}\}, y \neq C(x)\}$ with $|\mathcal{M}(C)| = 64 \times 20$ =1,280. The four definitional variants and their sizes are listed in Table S5.

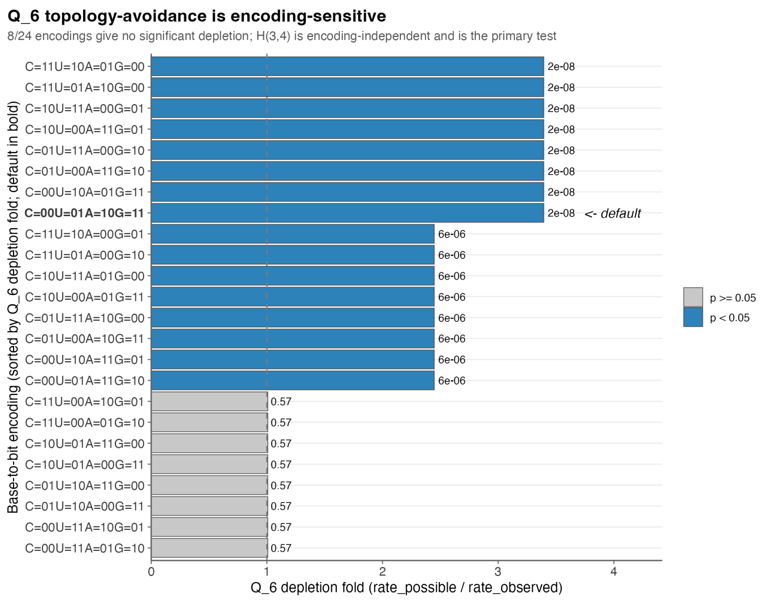


Figure S4: Per-encoding $Q_6$ topology-avoidance depletion across all 24 base-to-bit bijections. Bars are sorted by depletion fold; the dashed line at 1.0 marks no-depletion. Bars highlighted in blue are statistically significant ($p < 0.05$); grey bars are not. Eight of 24 encodings (depletion fold $\approx 1.0$, $p > 0.5$) place the $Q_6$ candidate-landscape rate at $\approx 36\%$ rather than 73%, eliminating the depletion signal. The default encoding (C=00, U=01, A=10, G=11) yields the largest depletion (3.4-fold). The encoding-independent $H(3,4)$ result is constant across all 24 bijections and is the primary topology-avoidance test in this work.

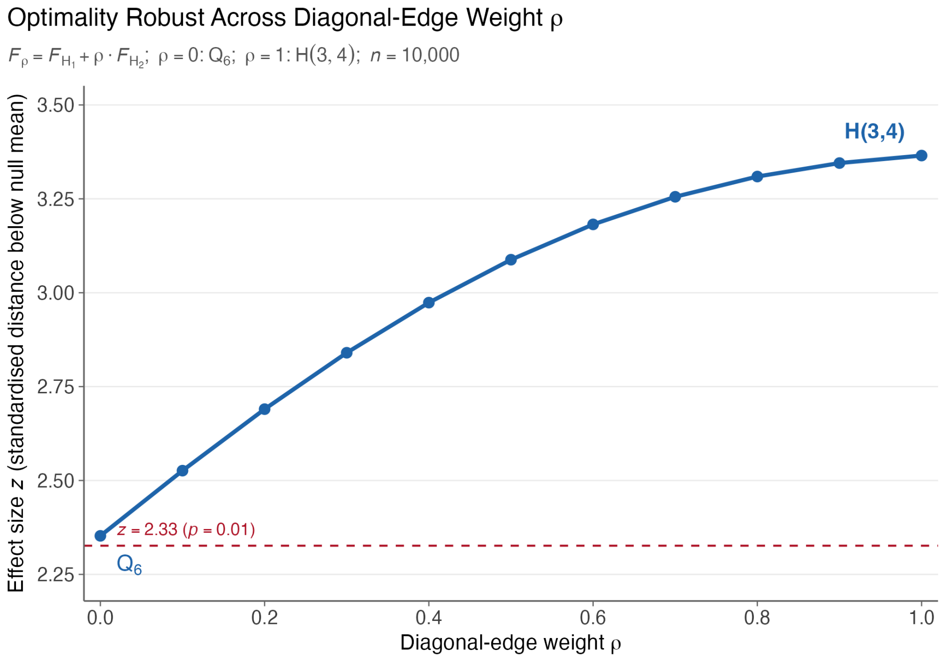


Figure S5: Coloring optimality across the $\rho$-interpolation between $Q_6$ (Hamming-1 only, $\rho = 0$) and $H(3,4)$ (full single-nucleotide mutation graph, $\rho = 1$). $F_\rho = F_{H_1} + \rho \cdot F_{H_2}$ is the diagonal-edge-weighted mismatch score; a lower $F_\rho$-percentile indicates stronger optimality. The standard code retains top-percentile placement across the full $\rho \in [0, 1]$ range under the default encoding, so the coloring-optimality result is not a knife-edge property of a specific weighting of the two edge classes. Companion to main-text §3.2.

| Universe | $\|\mathcal{M}\|$ | Definition |
|---|---|---|
| U1: 21-label, no identity (primary) | 1,280 | $y \in \mathcal{A}_{20} \cup \{\text{Stop}\}, y \neq C(x)$; each codon has exactly 20 alternative labels |
| U2: AA-only, no identity | 1,219 | $y \in \mathcal{A}_{20}, y \neq C(x)$; 61 sense codons get 19 AA alternatives, 3 stop codons get 20 AA alternatives |
| U3: AA, with no-ops | 1,280 | $y \in \mathcal{A}_{20}$, no identity restriction; each of 64 codons gets 20 AA candidates. Because $y \in \mathcal{A}_{20}$ excludes Stop, the 3 stop codons have no AA identity to match, so only the 61 sense codons contribute an AA no-op ($y = C(x)$); the row total 1,280 = 61 no-ops + 1,219 changes. |
| U4: stop-inclusive with no-ops | 1,344 | $y \in \mathcal{A}_{20} \cup \{\text{Stop}\}$, no identity restriction; 64 no-ops included |

Table S5: **Candidate-universe denominator definitions for the topology-avoidance test.** The four variants (U1–U4) interact along three definitional axes: whether stop-codon targets are admitted ($\mathcal{A}_{20} \cup \{\text{Stop}\}$ vs $\mathcal{A}_{20}$), whether identity moves $y = C(x)$ are excluded, and whether the source codon may itself be a stop codon. We adopt U1 as the primary universe because it (i) has a uniform alternative count per codon, (ii) cleanly excludes identity moves which contribute no signal, and (iii) admits stop-codon reassignment, which is biologically attested.

Topology-avoidance results under U2 and U4 are qualitatively identical (depletion remains $p < 10^{-5}$); the per-cell counts differ only by the constant rescaling $K_2 = K_1 - (\text{stop-target candidates})$ for U2. **Observed sense-to-Stop events under U2:** the observed reassignment corpus contains several sense-to-Stop events (e.g. NCBI translation table 22 UCA→Stop in *Scenedesmus obliquus* mitochondrial). Because U2 excludes Stop targets, these events are **not eligible** under U2 and are dropped from both the observed numerator and the candidate denominator of the U2 hypergeometric; the U2 test therefore evaluates whether the sense-to-sense subset of natural reassignments is topology-depleted against the sense-to-sense candidate landscape. Sense-to-Stop events remain in U1 and U4. Detailed per-universe numbers, including the exact observed counts kept or dropped under each variant, are released as part of the `codontopo` repository's `output/` artifacts (the `denominator_sensitivity` block in the topology-avoidance results file).

### S6 Conditional logit: IIA assumption and explanatory framing

The conditional-logit framework assumes Independence of Irrelevant Alternatives (IIA): for any two candidate moves $m_1$ and $m_2$ in a choice set $\mathcal{N}$, the relative probability $P(m_1)/P(m_2)$ is the same regardless of which other candidates are in $\mathcal{N}$. In the reassignment context, candidate moves are not exchangeable: a move that targets an amino acid already serviced by many codons is plausibly more substitutable with similar moves than the IIA structure allows.

We adopt IIA here because the goal is *explanatory* rather than *predictive*: the test asks whether topology adds explanatory value beyond physicochemical cost (LR test, ΔAICc) within the same candidate set, not whether the model accurately predicts which specific reassignment will occur next. IIA remains an **untested structural assumption** of the estimator, however, even for the explanatory reading: every candidate contributes to the choice-set denominator, so the composition and similarity of the non-observed alternatives affect the fitted likelihood, coefficients, and LR statistics. The restricted-candidate refits in Section 6.1 (and the alternative-specification fits reported alongside them) are best read as **sensitivity analyses** against candidate-set composition, not as evidence that IIA is harmless. A mixed logit relaxing IIA would be more appropriate for both explanatory calibration and prediction; future work could pursue this once a larger event set permits identification of mixed-logit covariance parameters. The parametric predictive simulation reported in the main text (§3.5) provides a calibration check on the model's marginal feature distribution rather than just in-sample AICc: the event-step topology-breaking rate under $Q_6$ adjacency with the **increase-in-components** ($\Delta\beta_0 > 0$) definition, observed at 5/66 = 0.076, is reproduced by the M3 simulation at mean $0.077 \pm 0.033$ ($n$ =10,000 draws; parametric predictive $p = 0.60$). This is the event-step / $Q_6$ / $\Delta\beta_0 > 0$ rate; the lineage-collapsed / $H(3,4)$ / new-disconnection rate quoted for the primary topology-avoidance test in §3.4 (6/28 = 0.21) is a different quantity and is not directly comparable.

A second, structurally distinct concern about the candidate set is its **composition**: whether the universe of ≈ 1,280 moves contains so many biologically implausible alternatives that the fitted topology coefficient is partly identifying their absence rather than a topology-specific avoidance effect. We address this concern in Section 6.1 by refitting M1–M4 on candidate sets that have been pruned to biologically plausible alternatives only.

#### S6.1 Candidate-set composition: restricted-candidate sensitivity

A separate concern about the candidate set is that the universe of ≈1,280 single-codon moves admits biologically catastrophic alternatives (reassigning AUG-Met, simultaneous multi-codon changes implicit in the single-step framing, reassignments to stop in essential codons) that natural selection has already removed from the option set. Models with strongly negative coefficients on $\Delta_{\text{topo}}$ and $\Delta_{\text{phys}}$ may therefore be partly rediscovering that natural reassignments are not biologically catastrophic, inflating ΔAICc magnitudes beyond what the explanatory thesis (topology adds value beyond physicochemistry) strictly requires. The qualitative claim is unaffected by this concern, but the magnitudes need calibration.

We address this by refitting M1–M4 (and the $H(3,4)$ verification variants) on a **restricted candidate set**: at each event-step we retain only candidates whose target amino acid is already serviced by a codon at Hamming distance $\leq d$ from the reassigned codon (i.e., $\Delta_{\text{tRNA}} \leq d$), with $d \in \{1,2,3\}$. The observed move is always retained regardless of its $\Delta_{\text{tRNA}}$ so the likelihood remains well-defined. We report all three filters as a **bracketing** sensitivity rather than designating any single $d$ as the calibrated biological cut, because the relevant decoding literature does not pin a single distance. The strictest cut, $d = 1$, is the closest match to canonical wobble-position mismatch tolerance (Crick, 1966) and is the most defensible biological-plausibility floor; $d = 3$ permits near-cognate routes and is included as a loose upper bound. We report $d \in \{1,2,3\}$ as a bracketing sensitivity rather than designating any single value as primary, because the decoding literature does not pin a single canonical distance and designating one without an independent biological anchor would risk reporting the most interpretable effect size rather than the most defensible one. Table S6 reports the resulting ΔAICc gaps under all four candidate sets.

| Filter | Mean cand. | Obs. kept | ΔAICc M1→M3 | ΔAICc M2→M3 | ΔAICc M3→M4 | ΔAICc M1→M3$H(3,4)$ |
|---|---|---|---|---|---|---|
| Unrestricted | 1280 | — | 108 | 89 | −2.1 | 91 |
| $\Delta_{\text{tRNA}} \leq 3$ | 1088 | 66 / 66 | 95 | 85 | −1.6 | 77 |
| $\Delta_{\text{tRNA}} \leq 2$ | 727 | 66 / 66 | 60 | 77 | 17.3 | 41 |
| $\Delta_{\text{tRNA}} \leq 1$ | 275 | 66 / 66 | 14 | 73 | 95.3 | 15 |

Table S6: Restricted-candidate sensitivity for the conditional-logit comparison. Each row refits M1, M2, M3, M4 (and the $H(3,4)$ topology variants) on a candidate set filtered to $\Delta_{\text{tRNA}} \leq d$, where $\Delta_{\text{tRNA}}$ is the Hamming distance from the reassigned codon to the nearest existing codon for the target amino acid. Observed moves are always retained so likelihoods remain comparable. The "Unrestricted" row reproduces the unrestricted main-text numbers. ΔAICc gaps shrink as the candidate set is restricted. This is expected: removing biologically implausible candidates removes contrasts the model otherwise exploits, so the table should be read as a **bracket**: the strictest biological-plausibility floor at $d = 1$ ( 275 candidates) gives

ΔAICc(M1→M3) $\approx$ 14, just above the conventional Burnham–Anderson reference, while the looser $d = 2$ filter ( 727 candidates) gives $\approx$ 60 and the looser $d = 3$ filter $\approx$ 95. ΔAICc(M2→M3) stays large at every threshold. The qualitative claim "topology adds explanatory value beyond physicochemistry" is therefore robust to candidate-set composition; the **magnitude** of the topology–physicochemistry separation is best characterised by the $d = 1$ floor rather than by the unrestricted $\approx$ 110 figure. The ΔAICc(M3→M4) column under the restricted filters is **not** interpretable, because the filter is defined on $\Delta_{\mathrm{tRNA}}$ so the M4 tRNA-feature distribution shifts mechanically with the filter; the unrestricted-set ΔAICc(M3→M4) of 2 is the calibrated reading and shows the heuristic tRNA proxy is uninformative.

Two structural points are worth flagging beyond the table itself. First, the ΔAICc(M1→M3) gap is exactly the quantity sensitive to candidate-set composition: physicochemistry alone (M1) is a flat baseline that benefits most from the implausible-candidate filler being present, so removing it shrinks the M1→M3 distance more than the M2→M3 distance. Second, ΔAICc(M2→M3) is robust across all three thresholds (≈73–85), reflecting that physicochemistry contributes the same incremental signal regardless of which candidates surround the observed events. The overall reading is therefore conservative: the unrestricted-set ΔAICc(M1 →M3) ≈ 110 over-states the **magnitude** of the topology–physicochemistry separation but does not over-state its existence, and the $d = 1$ floor ($\approx$ 14) provides the most defensible biological-plausibility-bounded effect size and shows the qualitative claim survives even under the strictest cut.

#### S7 Conditional logit: clade-exclusion sensitivity

The conditional-logit framework can confound a single clade's reassignment events with a global topology-avoidance signal if that clade is unusually extensive (or unusually idiosyncratic). To bound this concern, we refit M1–M4 under each of seven clade-exclusion regimes (Table S7), removing the indicated NCBI translation tables from the event-step list and re-running the full conditional-logit pipeline end-to-end (build candidate sets → enumerate event orderings up to $k!$ for $k \leq 6$ → vectorised maximum-likelihood fit → ΔAICc against M1 and M2). The seven exclusion regimes match those applied to the hypergeometric topology-avoidance test (Section 9), which themselves follow the phylogenetic-distribution analysis in Sengupta et al. (2007).

| **Excluded clade** | **Tables out** | $n$ | **ΔAICc M1→M3** | **ΔAICc M2→M3** | **ΔAICc M3→M4** |
|---|---|---|---|---|---|
| ciliates | 6, 10, 15, 27, 28, 29, 30 | 52 | 83.7 | 31.6 | 1.8 |
| yeast mito | 3 | 60 | 103.1 | 87.7 | 2.2 |
| CUG clade | 12, 26 | 64 | 116.7 | 94.9 | 2.2 |
| metazoan mito | 2, 5, 9, 13, 14, 21 | 40 | 48.8 | 88.6 | 0.4 |
| algal mito | 16, 22 | 63 | 115.5 | 85.7 | 2 |
| protist mito | 4, 23 | 63 | 100.7 | 92.4 | 1.9 |
| hemichordate mito | 24, 33 | 59 | 92.3 | 83.5 | 2.1 |

Table S7: **Per-regime conditional-logit clade-exclusion sensitivity (7 regimes).** Each row refits M1–M4 with the indicated NCBI translation tables removed; $n$ = remaining event-steps. M3 (physicochemistry + $Q_6$ topology) remains favored over both M1 (physicochemistry only) and M2 (topology only) in every regime. The minimum ΔAICc(M1→M3) across all seven regimes is 49 the maximum is 117. The conventional Burnham–Anderson reference of ΔAICc$>$ 10 was calibrated on linear-regression contexts, so we use it as a reference threshold rather than a formally calibrated cut-off in this conditional-logit setting; the parametric predictive simulation reported in main-text §3.5 (event-step topology-breaking rate under $Q_6$ / $\Delta\beta_0 > 0$: observed 5/66 = 0.076 vs simulated 0.077, $p = 0.60$) is the more directly interpretable calibration check. ΔAICc(M3→M4) values near 2 indicate that adding the heuristic tRNA-distance proxy provides no incremental fit.

#### S8 Per-table optimality: standard-code-proximity audit

This audit grounds the disaggregation reported in main-text §3.3, where we separated the per-table BH-FDR result into "informative-distance" tables ($\geq$ 3 reassignments from standard) and "near-standard" tables ($\leq$ 2 reassignments).

The methodological concern: a variant code that differs from the standard by only a few reassignments has a per-table quartet-pattern shuffle null distribution dominated by permutations very close to the standard code, simply because few permutations of the variant's block structure yield codes that are far from standard. In that limit, "table $X$ falls in the bottom 5% of permutations preserving $X$'s block structure" partly tests whether $X$ is close to the standard code (which it is, by construction), not whether $X$ is independently optimal.

For each NCBI translation table we computed three quantities alongside the unconditional per-table quantile: (i) the Hamming distance $d_H$ from the standard code (number of codons with a different AA label) for both the observed variant code and each quartet-pattern shuffle null draw, (ii) the variant's null quantile **conditional** on null draws within $\pm 2$ codons of the variant's $d_H$, and (iii) the fraction of null draws whose $d_H$ is at most the variant's $d_H$ (a complementary "proximity rank" of the variant against the null). The motivation for the conditional quantile was to ask whether variants with low unconditional $p$-values would still appear unusually low-cost when restricted to null draws of equivalent $d_H$. As Table S8 shows, the conditional bucket turns out to be empty for every variant: under the quartet-pattern shuffle, every null draw has $d_H \geq 30$ from the standard code (the lowest observed $d_H$ in the null distribution across all 27 tables is 30, while the highest variant $d_H$ in the registry is 6), so no null draws fall within $\pm 2$ codons of any variant's $d_H$. The conditional quantile is therefore **not informative** about whether a variant's optimality is intrinsic versus standard-code-proximity-driven.

The proximity-rank diagnostic in the rightmost column is informative: every variant in the registry has $d_H$ smaller than every null draw, so all variants sit at the extreme low-$d_H$ tail of the quartet-pattern shuffle null distribution. This means each variant's per-table $p$-value is, structurally, partly a proximity-to-standard-code measurement, and the "informative-distance" vs "near-standard" disaggregation in main-text §3.3 is therefore based on absolute $d_H$ thresholds (3 reassignments) rather than on this audit's conditional quantile.

| **Table** | **$d_H$ (variant)** | **Null $d_H$ range** | **Quantile (uncond.)** | **$n$ in $d_H \pm 2$ bucket** | **Frac. null $d_H \leq d_H^{\text{obs}}$** |
|---|---|---|---|---|---|
| **Informative-distance tables ($d_H \geq 3$ codon reassignments from standard).** | | | | | |
| 3 | 6 | 37–60 | 7.4% | 0 | 0% |
| 21 | 5 | 39–60 | 1.4% | 0 | 0% |
| 14 | 5 | 38–61 | 2.2% | 0 | 0% |
| 33 | 4 | 41–61 | 2% | 0 | 0% |
| 13 | 4 | 32–60 | 2.4% | 0 | 0% |

| Table | $d_H$ (variant) | Null $d_H$ range | Quantile (uncond.) | $n$ in $d_H \pm 2$ bucket | Frac. null $d_H \leq d_H^{\text{obs}}$ |
|---|---|---|---|---|---|
| 9 | 4 | 36–60 | 1.4% | 0 | 0% |
| 5 | 4 | 38–60 | 2.4% | 0 | 0% |
| 2 | 4 | 36–58 | 1.4% | 0 | 0% |
| 31 | 3 | 44–64 | 0.3% | 0 | 0% |
| 28 | 3 | 44–64 | 0.2% | 0 | 0% |
| 27 | 3 | 37–64 | 0.2% | 0 | 0% |
| 24 | 3 | 40–60 | 1.5% | 0 | 0% |
| **Near-standard tables ($d_H \leq 2$ codon reassignments from standard).** | | | | | |
| 30 | 2 | 38–60 | 0% | 0 | 0% |
| 29 | 2 | 38–60 | 2.4% | 0 | 0% |
| 23 | 2 | 35–57 | 0.8% | 0 | 0% |
| 22 | 2 | 30–54 | 0.8% | 0 | 0% |
| 6 | 2 | 40–60 | 0.3% | 0 | 0% |
| 32 | 1 | 35–57 | 1.5% | 0 | 0% |
| 26 | 1 | 32–56 | 1% | 0 | 0% |
| 25 | 1 | 36–60 | 1.1% | 0 | 0% |
| 16 | 1 | 33–57 | 1.9% | 0 | 0% |
| 15 | 1 | 35–57 | 0.3% | 0 | 0% |
| 12 | 1 | 35–56 | 0.8% | 0 | 0% |
| 10 | 1 | 40–60 | 0.5% | 0 | 0% |
| 4 | 1 | 38–60 | 1% | 0 | 0% |
| 11 | 0 | 34–56 | 1.2% | 0 | 0% |

Table S8: **Per-table standard-code-proximity audit.** For each NCBI translation table other than the standard code (table 1), $d_H$ is the number of codons with a different AA label from the standard code. **Null $d_H$ range** is the (min, max) of $d_H$ across the 10,000 quartet-pattern shuffle null draws for that table. **Quantile (uncond.)** is the variant's per-table quartet-pattern shuffle null quantile from main-text §3.3 (lower = more error-minimising). **$n$ in $d_H \pm 2$ bucket** counts null draws with $d_H$ within $\pm 2$ of the variant's $d_H$ (would have been the conditional-quantile denominator); this is zero for every table because every quartet-pattern shuffle null draw has $d_H \geq 30$ while every variant has $d_H \leq 6$. **Frac. null $d_H \leq d_H^{\text{obs}}$** is the fraction of null draws closer to (or as close as) the variant in $d_H$; this is uniformly 0%, confirming that every variant sits at the extreme low-$d_H$ tail of its quartet-pattern shuffle null. The conditional-quantile diagnostic is therefore not informative on this data, and the disaggregation in main-text §3.3 rests on the absolute $d_H \geq 3$ threshold rather than on this conditional analysis.

The full per-table audit, including $d_H$ summary statistics for the null distribution and the per-table observed score, is released in `output/coloring_optimality.json` under the `per_table_proximity_audit` key.

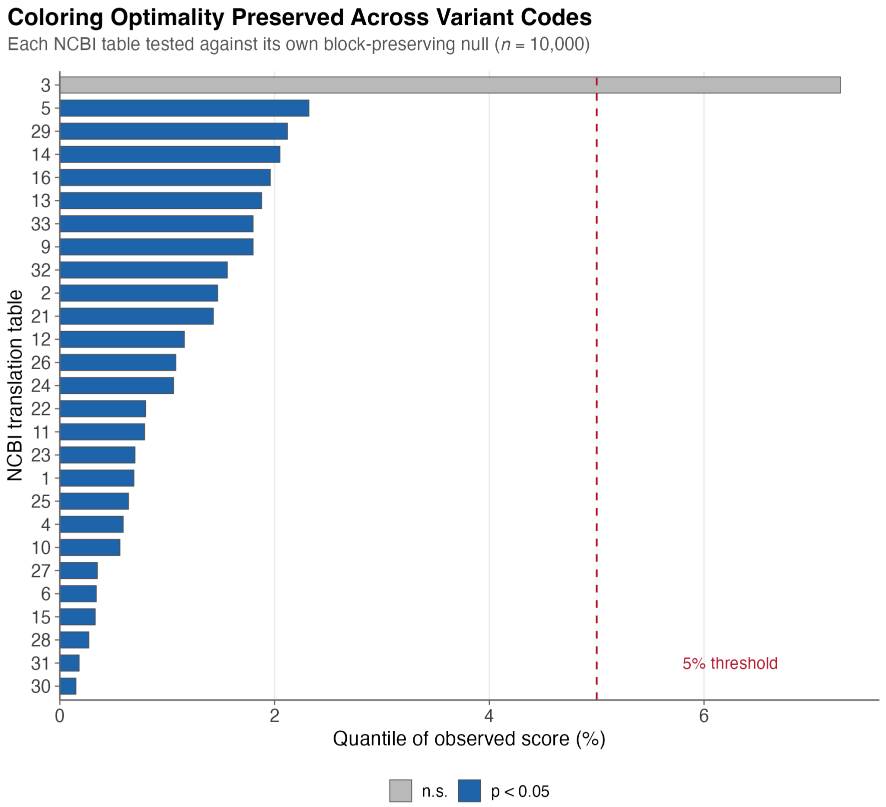


Figure S6: Coloring optimality preserved across variant codes. Per-table Grantham quartet-pattern shuffle null quantile for each of the 27 NCBI translation tables, tested against its own quartet-pattern shuffle null ($n$ =10,000 per table). Bars sorted by quantile; the dashed reference line marks the 5% threshold. Companion to main-text §3.3 and Figure 2C; the "informative-distance" vs "near-standard" disaggregation of §3.3 is not visible in this plot because it is a proximity-audit output rather than a null-quantile statistic (see Table S8).

### S9 Topology avoidance: clade-exclusion sensitivity (hypergeometric/permutation)

This section provides the hypergeometric/permutation counterpart to the conditional-logit clade-exclusion analysis in Section 7. The two tests probe related but distinct quantities: Section 7 tests whether the $\Delta_{topo}$ regression coefficient survives clade exclusion; this section tests whether the global landscape-vs-observed depletion ratio survives clade exclusion. Following the phylogenetic-distribution analysis of mitochondrial reassignment events by Sengupta et al. (2007), we iteratively excluded each of seven major taxonomic groups (Table S9) and re-ran the $Q_6$ topology-avoidance hypergeometric on the reduced event set, holding the 1,280-move candidate landscape fixed.

| Excluded clade | Tables out | $n$ | Breakers | Rate (%) | Hyper. $p$ |
|---|---|---|---|---|---|
| ciliates | 6, 10, 15, 27, 28, 29, 30 | 24 | 6 | 25 | $1.22 \times 10^{-6}$ |
| yeast mito | 3 | 24 | 2 | 8.3 | $3.64 \times 10^{-11}$ |
| CUG clade | 12, 26 | 26 | 5 | 19.2 | $1.41 \times 10^{-8}$ |
| metazoan mito | 2, 5, 9, 13, 14, 21 | 22 | 6 | 27.3 | $9.78 \times 10^{-6}$ |
| algal mito | 16, 22 | 26 | 5 | 19.2 | $1.41 \times 10^{-8}$ |

| Excluded clade | Tables out | $n$ | Breakers | Rate (%) | Hyper. $p$ |
|---|---|---|---|---|---|
| protist mito | 4, 23 | 27 | 6 | 22.2 | $4.78 \times 10^{-8}$ |
| hemichordate mito | 24, 33 | 27 | 6 | 22.2 | $4.78 \times 10^{-8}$ |

Table S9: **Hypergeometric clade-exclusion sensitivity for the $Q_6$ topology-avoidance test.** Each row removes the indicated NCBI translation tables (matching the clade definitions of Sengupta et al. (2007)) from the de-duplicated event list and re-runs the hypergeometric depletion test against the same 1,280-move candidate landscape. **Breakers** = topology-breaking events (new disconnection, $Q_6$ adjacency); **Rate** = observed breaker rate (vs candidate-landscape rate of 72.7% under the default encoding). All seven exclusions yield $p < 10^{-5}$. Excluding yeast mitochondrial (NCBI translation table 3) **strengthens** the depletion to $p \approx 3.6 \times 10^{-11}$ because that table contributes 4 of the 6 lineage-collapsed topology-breakers; this is the denominator-effect noted in main-text §3.4.

In every exclusion, the $Q_6$ depletion remains highly significant ($p < 10^{-5}$), confirming that the $\approx 3.4$-fold depletion of topology-breaking changes is a pan-taxonomic pattern, not an artifact of any single lineage.

The primary-cell $H(3,4)$ / $\Delta\beta_0 > 0$ analogue of the same seven-row exclusion is Table S10. It uses the encoding-independent nucleotide-adjacency graph and the increase-in-components definition matched to the conditional-logit feature (main-text §3.5), and confirms that the $\approx 3.1$-fold $H(3,4)$ depletion is also robust to clade exclusion.

| Excluded clade | Tables out | $n$ | Breakers | Rate (%) | Hyper. $p$ |
|---|---|---|---|---|---|
| ciliates | 6, 10, 15, 27, 28, 29, 30 | 24 | 6 | 25 | $3.96 \times 10^{-5}$ |
| yeast mito | 3 | 24 | 2 | 8.3 | $4.21 \times 10^{-9}$ |
| CUG clade | 12, 26 | 26 | 4 | 15.4 | $1.08 \times 10^{-7}$ |
| metazoan mito | 2, 5, 9, 13, 14, 21 | 22 | 6 | 27.3 | $2.03 \times 10^{-4}$ |
| algal mito | 16, 22 | 26 | 6 | 23.1 | $7.29 \times 10^{-6}$ |
| protist mito | 4, 23 | 27 | 6 | 22.2 | $3.07 \times 10^{-6}$ |
| hemichordate mito | 24, 33 | 27 | 6 | 22.2 | $3.07 \times 10^{-6}$ |

Table S10: **Hypergeometric clade-exclusion sensitivity for the primary $H(3,4)$ topology-avoidance test.** Each row removes the indicated NCBI translation tables (matching the clade definitions of Sengupta et al. (2007)) from the de-duplicated event list and re-runs the hypergeometric depletion test under the encoding-independent $H(3,4) = K_4^3$ adjacency with the increase-in-components ($\Delta\beta_0 > 0$) topology-breaking definition (candidate landscape: 846 breakers of 1,280 moves = 66.1%). **Breakers** = topology-breaking events by $\Delta\beta_0 > 0$; **Rate** = observed breaker rate. All seven exclusions yield $p < 10^{-3}$. Excluding yeast mitochondrial (NCBI translation table 3) strengthens the depletion to $p \approx 4.2 \times 10^{-9}$ because that table contributes 4 of the 6 lineage-collapsed $H(3,4)$-breakers — the direct calculation quoted in main-text §3.4 for the yeast-mito-excluded regime.

The clade-exclusion robustness is a denominator effect (removing clades that contributed zero or one topology-breaking event each leaves the breakage rate essentially unchanged) rather than a numerator effect; the **avoidance** of topology-breaking moves is what propagates across many lineages, while the **attempts** to break topology are concentrated in yeast mitochondrial. Detailed per-clade counts are released as `output/phylogenetic_sensitivity.json` (the $Q_6$ / new-disconnection cell) and `output/phylogenetic_sensitivity_k43.json` (the $H(3,4)$ / $\Delta\beta_0 > 0$ cell).

### S10 Complete tRNA gene count data

This section documents the empirical foundation for the tRNA enrichment analysis in main-text §3.6. We ran tRNAscan-SE 2.0.12 on 18 NCBI genome assemblies; 15 of these 18 organisms enter the 24-pairing Fisher–Stouffer enrichment analysis as either variant-code or standard-code-control repertoires (Table S11 below). The remaining 3 (*Blastocrithidia nonstop* P57; *Mycoplasmoides genitalium* G37; *Mycoplasmoides pneumoniae* M129) are used only as **mechanistic boundary cases** in main-text §4.3: their reassignment routes (anticodon stem shortening and anticodon modification respectively) act on a single tRNA gene without changing the gene-count distribution, so a Fisher-count enrichment test would report "no enrichment" and misinterpret the mechanism as an absence-of-response. Adding literature/database counts for a small number of legacy repertoires (see Table S11 "source-class" column: annotation, GtRNAdb, literature) brings the total pairing set to 24 pairings across 24 organisms with mixed provenance.

The Fisher-exact denominator convention used throughout is the **by-amino-acid sum**: for a focal amino acid $a$ in variant genome $V$ the $2 \times 2$ table is $\left[\left[a_V, \left(\sum_{a' \neq a} a'_V\right)\right], \left[a_C, \left(\sum_{a' \neq a} a'_C\right)\right]\right]$, where the by-amino-acid sum is over the **Std20** column of Table S13. This excludes SeC, undetermined-isotype, suppressor, and pseudogene rows so that the ratio $\frac{a}{\sum_{a'}} a'$ has a consistent interpretation across genomes (fractional tRNA-repertoire share for the focal AA). The 24 exact $2 \times 2$ tables and per-pairing Fisher $p$-values are printed in Table S11 and the machine-readable CSV `output/tables/T-trna-24row-provenance.csv`. The first-pass Isotype/Anticodon totals reported parenthetically in the Reassigned-AA column of Table S13 below are provided so the reader can see the per-anticodon breakdown; they are **not** the Fisher denominators.

#### S10.1 24-pairing input table with provenance and Fisher $2 \times 2$ counts

| **Variant** | **Control** | **AA** | **Reassignment** | $Q_6$ **break?** | **H(3,4)** | **Var** $\frac{a}{N}$ | **Ctl** $\frac{a}{N}$ | **OR** | **Fisher** $p$ |
|---|---|---|---|---|---|---|---|---|---|
| scerevisiae | ylipolytica | Thr | CUA: Leu→Thr; CUC: Leu→Thr; C... | Y | creates | 2/24 | 1/27 | 2.36 | 0.455 |
| sobliquus | creinhardtii | Leu | UAG: Stop→Leu | Y | no-effect | 2/22 | 1/16 | 1.5 | 0.621 |
| ptannophilus | lthermotolerans | Ala | CUG: Leu→Ala | Y | creates | 14/207 | 11/206 | 1.29 | 0.345 |
| calbicans | lthermotolerans | Ser | CUG: Leu→Ser | Y | extends | 16/207 | 13/206 | 1.24 | 0.355 |
| tthermophila | imultifiliis | Gln | UAA: Stop→Gln; UAG: Stop→Gln | n | no-effect | 54/712 | 3/141 | 3.78 | 0.008 |
| ptetraurelia | imultifiliis | Gln | UAA: Stop→Gln; UAG: Stop→Gln | n | no-effect | 18/215 | 3/141 | 4.2 | 0.01 |
| tthermophila | scoeruleus | Gln | UAA: Stop→Gln; UAG: Stop→Gln | n | no-effect | 54/712 | 11/267 | 1.91 | 0.032 |
| ptetraurelia | scoeruleus | Gln | UAA: Stop→Gln; UAG: Stop→Gln | n | no-effect | 18/215 | 11/267 | 2.13 | 0.04 |
| oxytricha | scoeruleus | Gln | UAA: Stop→Gln; UAG: Stop→Gln | n | no-effect | 8/90 | 11/267 | 2.27 | 0.075 |
| eoctocarinatus | scoeruleus | Cys | UGA: Stop→Cys | n | no-effect | 5/114 | 7/267 | 1.7 | 0.272 |
| bjaponicum | scoeruleus | Trp | (no direct reassignment for t... | n | no-effect | 5/115 | 6/267 | 1.98 | 0.21 |
| eaediculatus | scoeruleus | Cys | UGA: Stop→Cys | n | no-effect | 4/77 | 7/267 | 2.04 | 0.215 |

| Variant | Control | AA | Reassignment | $Q_6$ break? | H(3,4) | Var $\frac{a}{N}$ | Ctl $\frac{a}{N}$ | OR | Fisher $p$ |
|---|---|---|---|---|---|---|---|---|---|
| eaediculatus | fsalina | Cys | UGA: Stop→Cys | n | no-effect | 4/77 | 1/86 | 4.66 | 0.151 |
| eamieti | scoeruleus | Cys | UGA: Stop→Cys | n | no-effect | 8/116 | 7/267 | 2.75 | 0.049 |
| efocardii | fsalina | Cys | UGA: Stop→Cys | n | no-effect | 3/59 | 1/86 | 4.55 | 0.184 |
| eparawoodruffi | scoeruleus | Cys | UGA: Stop→Cys | n | no-effect | 9/145 | 7/267 | 2.46 | 0.065 |
| eweissei | fsalina | Cys | UGA: Stop→Cys | n | no-effect | 20/463 | 1/86 | 3.84 | 0.132 |
| ewoodruffi | scoeruleus | Cys | UGA: Stop→Cys | n | no-effect | 4/79 | 7/267 | 1.98 | 0.226 |
| ppersalinus | imultifiliis | Gln | UAA: Stop→Gln; UAG: Stop→Gln | n | no-effect | 20/260 | 3/141 | 3.83 | 0.015 |
| ppersalinus | fsalina | Gln | UAA: Stop→Gln; UAG: Stop→Gln | n | no-effect | 20/260 | 3/86 | 2.31 | 0.132 |
| hgrandinella | scoeruleus | Gln | UAA: Stop→Gln; UAG: Stop→Gln | n | no-effect | 9/129 | 11/267 | 1.75 | 0.165 |
| hgrandinella | fsalina | Gln | UAA: Stop→Gln; UAG: Stop→Gln | n | no-effect | 9/129 | 3/86 | 2.08 | 0.218 |
| bstoltei | scoeruleus | Trp | (no direct reassignment for t... | n | no-effect | 6/166 | 6/267 | 1.63 | 0.289 |
| bstoltei | fsalina | Trp | (no direct reassignment for t... | n | no-effect | 6/166 | 2/86 | 1.58 | 0.447 |

Table S11: **Complete 24-pairing tRNA-enrichment input table (analysis columns).** For each pairing: variant and control organism (short key; full organism name in Table S13), reassigned amino acid, brief reassignment description, whether the reassignment creates a new $Q_6$ AA-family disconnection at $\varepsilon = 1$ (Y/n), and the $H(3,4)$ impact (creates / extends / no-effect / n/a for pairings without a $Q_6$ disconnection). **Var** $\frac{a}{N}$, **Ctl** $\frac{a}{N}$: focal AA count over the by-amino-acid-sum denominator. **OR**: Fisher-exact odds ratio (variant vs control). **Fisher** $p$: one-sided upper-tail $p$-value. Row-level provenance (compartment, NCBI translation table, source class, and assembly accession or primary citation for each of the 48 organism-slots) is given in the companion Table S12. Source-class breakdown across the 48 organism-slots (24 variant + 24 control): tRNAscan-SE 2.0.12 = 38, literature = 5, annotation = 3, GtRNAdb = 2. Unique organisms: 24; unique organisms with tRNAscan-SE-verified counts: 15. Machine-readable CSV (with `variant_source_full`, `control_source_full`, and every column shown here): `output/tables/T-trna-24row-provenance.csv`.

| Pairing (V ← C) | AA | Cmpt | NCBI tbl | Variant source (class; accession or citation) | Control source (class; accession or citation) |
|---|---|---|---|---|---|
| scerevisiae ← ylipolytica | Thr | M | 3 | annotation Bonitz et al. 1980 PNAS 77:3167-3170 (PMC349575); Su et al. 2011 PMC3113583; RefSeq NC_001224.1 (S. cerevisiae S288C mitochondrion) | literature Kerscher et al. 2001 |
| sobliquus ← creinhardtii | Leu | M | 22 | literature NCBI Scenedesmus obliquus mitochondrial genome; Knaap et al. 2002 | annotation Chlamydomonas reinhardtii mitochondrial genome annotation |
| ptannophilus ← lthermotolerans | Ala | N | 26 | literature Muhlhausen & Kollmar 2014, Genome Biology and Evolution | GtRNAdb GtRNAdb for Lachancea thermotolerans CBS 6340 |
| calbicans ← lthermotolerans | Ser | N | 12 | annotation Candida Genome Database; Santos et al. 1996-2011 reviews | GtRNAdb GtRNAdb for Lachancea thermotolerans CBS 6340 |

| Pairing (V ← C) | AA | Cmpt | NCBI tbl | Variant source (class; accession or citation) | Control source (class; accession or citation) |
|---|---|---|---|---|---|
| tthermophila ← imultifiliis | Gln | N | 6 | tRNAscan-SE 2.0.12 GCF_000189635.1 | tRNAscan-SE 2.0.12 GCF_000220395.1 |
| ptetraurelia ← imultifiliis | Gln | N | 6 | tRNAscan-SE 2.0.12 GCF_000165425.1 | tRNAscan-SE 2.0.12 GCF_000220395.1 |
| tthermophila ← scoeruleus | Gln | N | 6 | tRNAscan-SE 2.0.12 GCF_000189635.1 | tRNAscan-SE 2.0.12 GCA_001970955.1 |
| ptetraurelia ← scoeruleus | Gln | N | 6 | tRNAscan-SE 2.0.12 GCF_000165425.1 | tRNAscan-SE 2.0.12 GCA_001970955.1 |
| oxytricha ← scoeruleus | Gln | N | 6 | tRNAscan-SE 2.0.12 GCA_000295675.1 | tRNAscan-SE 2.0.12 GCA_001970955.1 |
| eoctocarinatus ← scoeruleus | Cys | N | 10 | literature Meyer et al. 1991 PNAS 88:3758-3761; Grimm et al. 1998 NAR 26:4557; Wang et al. 2016 Sci Rep 6:21139 (genome/transcriptome) | tRNAscan-SE 2.0.12 GCA_001970955.1 |
| bjaponicum ← scoeruleus | Trp | N | 15 | literature Liang & Heckmann 1993; Swart et al. 2016 Mol Biol Evol 33:2885 (code survey) | tRNAscan-SE 2.0.12 GCA_001970955.1 |
| eaediculatus ← scoeruleus | Cys | N | 10 | tRNAscan-SE 2.0.12 GCA_030463445.1 | tRNAscan-SE 2.0.12 GCA_001970955.1 |
| eaediculatus ← fsalina | Cys | N | 10 | tRNAscan-SE 2.0.12 GCA_030463445.1 | tRNAscan-SE 2.0.12 GCA_022984795.1 |
| eamieti ← scoeruleus | Cys | N | 10 | tRNAscan-SE 2.0.12 GCA_048569255.1 | tRNAscan-SE 2.0.12 GCA_001970955.1 |
| efocardii ← fsalina | Cys | N | 10 | tRNAscan-SE 2.0.12 GCA_001880345.2 | tRNAscan-SE 2.0.12 GCA_022984795.1 |
| eparawoodruffi ← scoeruleus | Cys | N | 10 | tRNAscan-SE 2.0.12 GCA_021440025.1 | tRNAscan-SE 2.0.12 GCA_001970955.1 |
| eweissei ← fsalina | Cys | N | 10 | tRNAscan-SE 2.0.12 GCA_021440005.1 | tRNAscan-SE 2.0.12 GCA_022984795.1 |
| ewoodruffi ← scoeruleus | Cys | N | 10 | tRNAscan-SE 2.0.12 GCA_027382605.1 | tRNAscan-SE 2.0.12 GCA_001970955.1 |
| ppersalinus ← imultifiliis | Gln | N | 6 | tRNAscan-SE 2.0.12 GCA_001447515.1 | tRNAscan-SE 2.0.12 GCF_000220395.1 |
| ppersalinus ← fsalina | Gln | N | 6 | tRNAscan-SE 2.0.12 GCA_001447515.1 | tRNAscan-SE 2.0.12 GCA_022984795.1 |
| hgrandinella ← scoeruleus | Gln | N | 6 | tRNAscan-SE 2.0.12 GCA_006369765.1 | tRNAscan-SE 2.0.12 GCA_001970955.1 |
| hgrandinella ← fsalina | Gln | N | 6 | tRNAscan-SE 2.0.12 GCA_006369765.1 | tRNAscan-SE 2.0.12 GCA_022984795.1 |
| bstoltei ← scoeruleus | Trp | N | 15 | tRNAscan-SE 2.0.12 GCA_965603825.1 | tRNAscan-SE 2.0.12 GCA_001970955.1 |
| bstoltei ← fsalina | Trp | N | 15 | tRNAscan-SE 2.0.12 GCA_965603825.1 | tRNAscan-SE 2.0.12 GCA_022984795.1 |

Table S12: **Row-level provenance for the 24 analysis pairings.** Compartment (M = mitochondrial, N = nuclear; both if variant and control differ). **NCBI tbl** is the variant's NCBI translation table (the control is always table 1). **Source class** is one of tRNAscan-SE 2.0.12, GtRNAdb, literature, or annotation. Assembly accessions are shown for tRNAscan-verified rows; primary literature citations replace accessions for literature-derived rows (notably *S. cerevisiae* mitochondrial from Bonitz et al. (1980) and *Y. lipolytica* mitochondrial from Kerscher et al. (2001)). This table is emitted from the same 24-row input as

Table S11 and its full-fidelity source strings are in `output/tables/T-trna-24row-provenance.csv` under the `variant_source_full` and `control_source_full` fields.

#### S10.2 tRNAscan-SE verified organisms

All tRNA gene counts were obtained by running tRNAscan-SE 2.0.12 (Chan and Lowe, 2019) with Infernal 1.1.4 on NCBI genome assemblies. Eukaryotic organisms were scanned in `-E` mode; *Mycoplasmoides* species in `-B` (bacterial) mode. Total counts are Infernal-confirmed (the more conservative count after the second-pass filter) and match the Total column of Table S13 below; the summary that underlies the main-text Fisher–Stouffer combination is reported as main-text Table 7. The five-column breakdown in Table S13 sums exactly to Total: Std20 (decoding the standard 20 amino acids) + SeC (selenocysteine, anticodon UCA, scanned via the dedicated `TRNAinf-euk-SeC.cm` model) + Supp (possible suppressor tRNAs with CTA/TTA/UCA anticodons) + Undet (predicted tRNAs whose isotype could not be determined) + Pseudo (predicted pseudogenes filtered by the Infernal second-pass isotype validation).

| Organism | Tbl | Assembly | Total | Std20 | SeC | Supp | Undet | Pseudo | Reassigned AA |
|---|---|---|---|---|---|---|---|---|---|
| *T. thermophila* | 6 | GCF_000189635.1 | 718 | 672 | 1 | 37 | 4 | 4 | Gln: 54 (15+39) |
| *P. tetraurelia* | 6 | GCF_000165425.1 | 216 | 202 | 1 | 11 | 0 | 2 | Gln: 18 (7+11) |
| *O. trifallax* | 6 | GCA_000295675.1 | 94 | 83 | 2 | 6 | 2 | 1 | Gln: 8 (2+6) |
| *P. persalinus* | 6 | GCA_001447515.1 | 262 | 228 | 1 | 15 | 0 | 18 | Gln: 20 (5+15) |
| *H. grandinella* | 6 | GCA_006369765.1 | 130 | 121 | 1 | 3 | 0 | 5 | Gln: 9 (6+3) |
| *E. aediculatus* | 10 | GCA_030463445.1 | 80 | 76 | 1 | 2 | 1 | 0 | Cys: 4 (3+1 UCA) |
| *E. amieti* | 10 | GCA_048569255.1 | 120 | 103 | 1 | 6 | 1 | 9 | Cys: 8 (4+4 UCA) |
| *E. focardii* | 10 | GCA_001880345.2 | 62 | 56 | 1 | 3 | 0 | 2 | Cys: 3 (1+2 UCA) |
| *E. parawoodruffi* | 10 | GCA_021440025.1 | 149 | 128 | 0 | 3 | 3 | 15 | Cys: 9 (5+4 UCA) |
| *E. weissei* | 10 | GCA_021440005.1 | 495 | 390 | 0 | 12 | 14 | 79 | Cys: 20 (17+3 UCA) |
| *E. woodruffi* | 10 | GCA_027382605.1 | 83 | 74 | 1 | 3 | 1 | 4 | Cys: 4 (2+2 UCA) |
| *B. stoltei*[§] | 15 | GCA_965603825.1 | 169 | 165 | 1 | 0 | 1 | 2 | Trp: 6 |
| *B. nonstop* P57 | 31 | GCA_028554745.1 | 68 | 65 | 1 | 2 | 0 | 0 | Trp: 2[*] |
| *M. genitalium* | 4 | GCA_000027325.1 | 36 | 35 | 0 | 1 | 0 | 0 | Trp: 1[†] |
| *M. pneumoniae* | 4 | GCF_910574535.1 | 37 | 36 | 0 | 1 | 0 | 0 | Trp: 1[†] |
| *S. coeruleus* | 1 | GCA_001970955.1 | 272 | 265 | 1 | 1 | 3 | 2 | — |
| *I. multifiliis* | 1 | GCF_000220395.1 | 150 | 141 | 1 | 8 | 0 | 0 | — |
| *F. salina* | 1 | GCA_022984795.1 | 89 | 85 | 1 | 1 | 1 | 1 | — |

Table S13: Complete tRNAscan-SE 2.0.12 results. The **Supp** column and the **Std20/SeC/Undet/Pseudo** columns are Infernal-confirmed second-pass counts and sum exactly to **Total**: **Std20** = standard 20-AA tRNAs; **SeC** = selenocysteine (anticodon UCA); **Supp** = possible suppressor tRNAs (CTA/TTA/UCA anticodons that could read stop codons) **after** Infernal's isotype-validation filter; **Undet** = predicted tRNAs whose isotype could not be determined; **Pseudo** = predicted pseudogenes filtered by the second-pass validation. The Reassigned-AA breakdown, in contrast, uses tRNAscan-SE's **first-pass** Isotype/Anticodon counts, which are the per-anticodon totals reported directly by the covariance-model scan before the Infernal second-pass reclassification. Consequently the first-pass suppressor total in the Reassigned-AA parenthetical can exceed the second-pass **Supp** column when Infernal reclassifies borderline hits into **Std20** or **Pseudo**; the differences here are 0–2 tRNAs per organism (e.g. *T. thermophila*: 37 second-pass suppressors versus 39 first-pass = 9 CTA + 30 TTA; *E. parawoodruffi*: 3 versus 5 = 0 CTA + 1 TTA + 4 UCA). All Fisher-exact tests in Table 7 and §S10 use the first-pass per-amino-acid totals shown in the Reassigned-AA column, which are the biologically interpretable per-anticodon counts. [§]*Blepharisma stoltei* is indexed as NCBI translation table 15 (Blepharisma Nuclear Code, defined as UAG→Gln) by virtue of its genus assignment, but the strain-specific MAC genome reads UGA→Trp via a dedicated suppressor tRNA-Trp(UCA) (Singh et al., 2023). The analysis here tests Trp-tRNA enrichment on that empirical reading rather than the legacy table-15 nominal code. [*]Anticodon stem shortening (4-bp stem rather than canonical 5-bp). [†]Post-transcriptional modification (a single tRNA-Trp(CCA) reads both UGG and UGA after base modification).

#### S10.3 MIS (maximal independent set) analysis

To address non-independence from shared control organisms, we constructed a conflict graph in which edges connect pairings sharing an organism. Enumerating all maximal independent sets (MIS) is equivalent to enumerating all maximal cliques of the complement graph, which we perform via the Bron–Kerbosch algorithm applied to the complement (with pivoting). Each MIS represents a set of pairings where no two share an organism and no additional pairing can be added without creating a conflict.

The 24-pairing conflict graph admits **332** MIS, all of size 6. Across these 332 sets the Stouffer combined $p$-value distribution is as follows: median $p = 0.037$, best $p = 0.012$, worst $p = 0.104$; 264 of 332 (79.5%) fall below the 0.05 threshold and 0 of 332 below 0.01. The extreme cases:

- **Best-case MIS** ($p = 0.012$): *S. cerevisiae* mito/Thr, *S. obliquus* mito/Leu, *P. tannophilus*/Ala, *P. tetraurelia*/Gln, *T. thermophila*/Gln, *P. persalinus*/Gln.
- **Worst-case MIS** ($p = 0.104$): *S. cerevisiae* mito/Thr, *S. obliquus* mito/Leu, *C. albicans*/Ser, *E. octocarinatus*/Cys, *P. persalinus*/Gln, *B. stoltei*/Trp.

Because the median just clears the 0.05 threshold while the worst case does not, the tRNA enrichment result is treated as **exploratory** in the claim hierarchy (Table S1): the signal is present in the median-independent-subset sense but is not robust to worst-case subset choice.

**Independence-graph caveat.** The conflict graph encodes shared **organisms**, not phylogenetic distance. Two ciliates from different orders are treated as independent in this construction even though they may share more evolutionary signal than two arbitrary eukaryotes; at the same time, every MIS retains the *S. cerevisiae* mito/Thr pairing whose tRNA counts come from GtRNAdb rather than from a tRNAscan-SE 2.0.12 run on the assembly (see §S10.3). A phylogenetic-distance–based independence criterion would yield a smaller effective $n$ (the same 4–6 phylogenetically independent origins discussed in main-text Limitations); the MIS analysis should be read as bounding subset variability under the organism-shared-by-pairings criterion, not as a phylogeny-aware test.

**Topology-breaking subset.** As a pre-specified mechanistic complement to the all-pairings analysis, we restrict the Fisher–Stouffer combination to the four pairings whose underlying reassignment creates a new amino-acid disconnection under $Q_6$: *S. cerevisiae* mito/Thr (table 3), *S. obliquus* mito/Leu (table 22, equivalent to chlorophycean mito table 16), *P. tannophilus*/Ala (table 26), and *C. albicans*/Ser (table 12). This subset combination yields Stouffer $p = 0.387$, i.e. null. The all-pairings signal is therefore driven largely by topology-preserving stop-to-sense reassignment systems (UAR→Gln in ciliates, UGA→Cys in *Euplotes*, UGA→Trp in *Blepharisma*), and the tRNA-duplication mechanism is not evidenced by the topology-breaking cases in isolation. This is why the claim hierarchy treats the tRNA result as exploratory-decoding-accommodation rather than as evidence of compensation-for-disconnection.

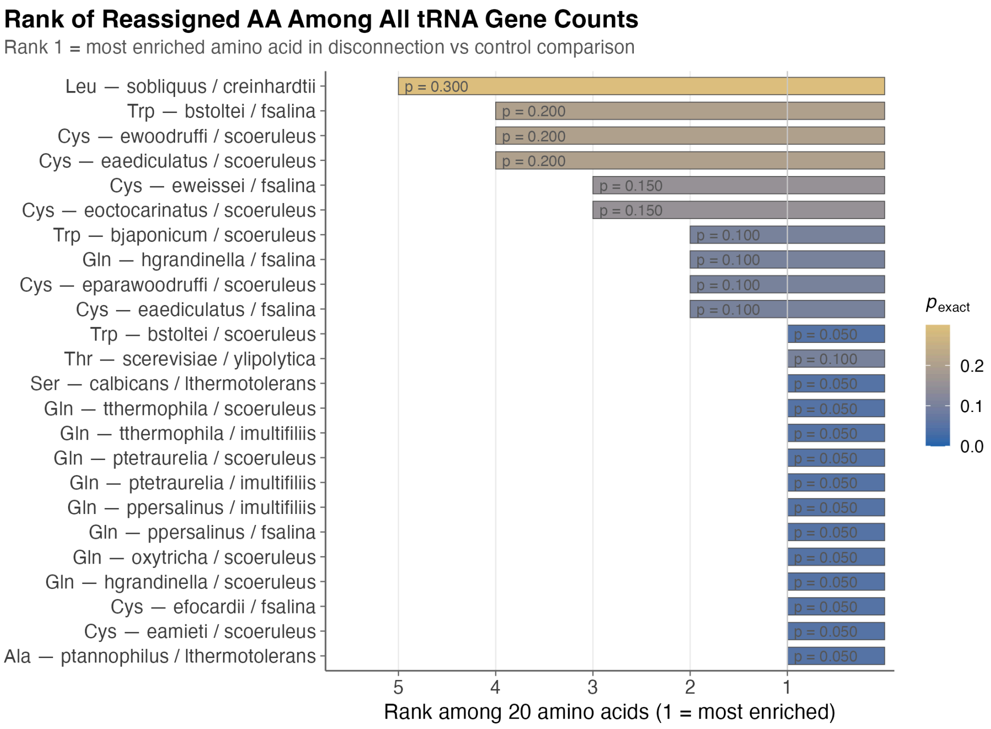


Figure S7: Rank of the reassigned amino acid among all tRNA gene counts, per variant-code × control pairing. Rank 1 = most enriched amino acid in the variant-code-vs-control comparison. (Most of the pairings here are topology-preserving stop-to-sense reassignments — UAR→Gln, UGA→Cys, UGA→Trp — rather than topology-breaking; the "disconnection" framing that appears in earlier drafts is inaccurate for the majority of pairings.) The distribution is left-heavy toward rank 1–3 across pairings, consistent with heterogeneous accommodation-of-decoding across variant-code lineages. The associated statistical summary (main-text §3.6, Table 7): 332 MIS of size 6, median Stouffer $p = 0.037$, worst $p = 0.104$; topology-breaking-only subset ($n = 4$) is null (Stouffer $p = 0.387$).

##### S10.4 *Saccharomyces cerevisiae* literature-derived control

The *Saccharomyces cerevisiae* tRNA gene counts used in the yeast-mito Thr disconnection pairing come from GtRNAdb (Chan and Lowe, 2016) rather than tRNAscan-SE 2.0.12 in this work. We retain them to keep the well-characterized yeast-mito Thr case in the pairing set and flag the difference in source explicitly.

#### S11 Complete reassignment database

The reassignment database underpins both the topology-avoidance test (main-text §3.4) and the conditional-logit fits (main-text §3.5). It comprises every codon reassignment event across the 27 NCBI translation tables, relative to the standard code (table 1). Each event records the codon, source amino acid, target amino acid, and Hamming distance to the nearest codon already encoding the target amino acid in the standard code. De-duplication to unique (codon, target amino acid) pairs yields the event set used in the topology-avoidance hypergeometric and table-preserving permutation tests. The conditional-logit model operates on the full per-table event-step list, which preserves recurrent reassignments across independent lineages. Exact counts depend on the most recent NCBI gc.prt revision (currently v4.6 retrieved 2026-04-25); a current pipeline run yields 66 raw events de-duplicating to 28 unique (codon, target-AA) pairs.

**Why two counts ($n$ =28 hypergeometric vs $n$ =66 conditional-logit).** The hypergeometric topology-avoidance test (main-text §3.4) uses the de-duplicated $n$ =28 unique (codon, target-AA) pairs, since the test is a sample-from-a-finite-landscape contrast that should not double-count the same move. The conditional-logit comparison (main-text §3.5) uses the full per-table $n$ =66 event-step list because each event-step is a **separate choice**: a recurrent reassignment such as UGA→Trp seen across many independent mitochondrial lineages contributes one independent choice per lineage to the conditional likelihood, since each lineage's reassignment is a distinct evolutionary event conditional on that lineage's candidate set. The conditional-logit clade-exclusion sensitivity (Section 7) inherits this convention: removing a clade removes all of its event-steps from the per-table list, including recurrent UGA→Trp instances within that clade, so the ΔAICc magnitudes report the **clade-removed-as-event-steps** sensitivity. We did not additionally re-weight by recurrence (e.g., one event per (codon, target) pair rather than per lineage); that alternative weighting would correspond more closely to the de-duplicated hypergeometric framing and would shrink ΔAICc magnitudes toward the restricted-candidate $d = 1$ floor reported in Section 6.1, but does not change the qualitative ranking M3 $>$ M1, M2.

| Table | NCBI translation table | Events |
|---|---|---|
| 2 | Vertebrate Mitochondrial | 4 |
| 3 | Yeast Mitochondrial | 6 |
| 4 | Mold/Protozoan/Coelenterate Mito | 1 |
| 5 | Invertebrate Mitochondrial | 4 |
| 6 | Ciliate Nuclear | 2 |
| 9 | Echinoderm/Flatworm Mito | 4 |
| 10 | Euplotid Nuclear | 1 |
| 12 | Alternative Yeast Nuclear | 1 |
| 13 | Ascidian Mitochondrial | 4 |
| 14 | Alternative Flatworm Mito | 5 |
| 15 | Blepharisma Macronuclear | 1 |
| 16 | Chlorophycean Mito | 1 |
| 21 | Trematode Mitochondrial | 5 |
| 22 | Scenedesmus obliquus Mito | 2 |
| 23 | Thraustochytrium Mito | 2 |
| 24 | Rhabdopleuridae Mito | 3 |
| 25 | Candidate Division SR1 / Gracilibacteria | 1 |
| 26 | Pachysolen tannophilus Nuclear | 1 |
| 27 | Karyorelict Nuclear | 3 |
| 28 | Condylostoma Nuclear | 3 |
| 29 | Mesodinium Nuclear | 2 |
| 30 | Peritrich Nuclear | 2 |
| 31 | Blastocrithidia Nuclear | 3 |
| 32 | Balanophoraceae Plastid | 1 |
| 33 | Cephalodiscidae Mito | 4 |

Table S14: **Per-table reassignment-event counts (raw, before de-duplication).** The 25 NCBI translation tables that differ from the standard code account for 66 raw codon-reassignment events. Tables 1 (Standard) and 11 (Bacterial / Archaeal / Plant Plastid) share identical sense-codon mappings and contribute zero reassignment events. Tables 27 (Karyorelict Nuclear) and 28 (Condylostoma Nuclear) likewise share identical sense-codon mappings and are listed separately to match NCBI numbering. The 25-table counts sum to the all-events total, which de-duplicates to 28 unique (codon, target-AA) pairs across the registry. The complete event-level database (with codon, source-AA, target-AA, Hamming-to-nearest-target columns) is released as `output/tables/T10_reassignment_db.csv` in the `codontopo` repository.

#### S12 Structural-preservation index (visualization-only)

For Figure 5A of the main text we use a heuristic **structural-preservation index** $S(m) \in [0, 1]$ for each candidate single-codon reassignment $m$ from the standard code. The index is **not** used in any inferential test in the manuscript; it is a visualization aid for delineating high- versus low-structure-preserving regions of the 1,280-move candidate landscape. We report the exact implementation here for completeness.

The index is a weighted sum of four discrete indicators. Given the reassignment $m$, let $I_{\mathrm{tw}(m)}, I_{\mathrm{ff}(m)} \in \{0, 1\}$ record whether the post-reassignment code preserves the two-fold bit-5 filtration and the four-fold prefix filtration described in main-text §2.2; let $I_{\mathrm{ser}(m)} \in \{0, 1\}$ record whether Serine remains disconnected at $\varepsilon = 1$; and let $n_{\mathrm{disc}(m)}$ denote the number of amino acids disconnected at $\varepsilon = 1$ under $m$. Then

$$S(m) = 0.25 \cdot I_{\mathrm{tw}(m)} + 0.25 \cdot I_{\mathrm{ff}(m)} + 0.30 \cdot I_{\mathrm{ser}(m)} + w\Big(n_{\mathrm{disc}(m)}\Big),$$

where $w(1) = 0.20$ (a single disconnected amino acid, as in the standard code), $w(0) = 0.10$ (partial credit for a fully connected variant), and $w(n) = 0.00$ for $n \geq 2$. Because every component is discrete, $S(m)$ can take only a small number of values across the 1,280-move candidate universe; the four bars visible in Figure 5A ($S = 0.55, 0.75, 0.80, 1.00$; 193/27/738/322 events) enumerate the values actually realised. The weights are a hand-tuned prioritisation of the four structural features and are not derived from any calibration; the index is descriptive, not inferential. Exact implementation: `src/codon_topo/analysis/synbio_feasibility.py` in the `codontopo` repository.

#### S13 KRAS–Fano clinical prediction: detailed results

The KRAS–Fano clinical prediction is the most concrete biomedical extrapolation considered for the $\mathrm{GF}(2)^6$ framework, and we tested it directly to delimit scope. The conjecture: XOR ("Fano") relationships in $\mathrm{GF}(2)^6$ predict enrichment of specific amino acids at KRAS G12 co-mutation sites. For each of the six common somatic KRAS G12 variants, we computed the predicted Fano partner codon as $GGU \oplus m_i = \mathrm{partner}_i$ where $m_i$ is the mutant codon for that variant and $\oplus$ is bitwise XOR in $\mathrm{GF}(2)^6$ (Table S15). We then tested for co-mutation enrichment of the predicted partner amino acid against 1,670 KRAS-mutant samples from the MSK-IMPACT pan-cancer dataset (Zehir et al., 2017) using Fisher's exact test with Bonferroni correction across the six variants.

| Variant | WT codon | Mutant codon | Fano partner codon | Predicted partner AA | Fisher $p$ (Bonf.) |
|---|---|---|---|---|---|
| G12V | GGU | GUU | CAC | His (H) | 1.0 |
| G12D | GGU | GAU | CUC | Leu (L) | 1.0 |

| Variant | WT codon | Mutant codon | Fano partner codon | Predicted partner AA | Fisher $p$ (Bonf.) |
|---|---|---|---|---|---|
| G12A | GGU | GCU | CGC | Arg (R) | 1.0 |
| G12R | GGU | CGU | GCC | Ala (A) | 1.0 |
| G12C | GGU | UGU | ACC | Thr (T) | 1.0 |
| G12S | GGU | AGU | UCC | Ser (S) | 1.0 |

Table S15: **KRAS–Fano clinical prediction tested against MSK-IMPACT.** For each KRAS G12 variant, the Fano partner codon is computed as the bitwise XOR (in $\mathrm{GF}(2)^6$) of the wild-type and mutant codons against a fixed reference; the predicted partner amino acid is the standard-code translation of that codon. Fisher's exact test for co-mutation enrichment of the predicted partner amino acid in MSK-IMPACT samples bearing each KRAS G12 variant ($n$ =1,670 KRAS-mutant tumours; (Zehir et al., 2017)) yields $p = 1.0$ for all six variants after Bonferroni correction across the six tests. Odds ratios are near 1.0 throughout. The conjecture is cleanly falsified.

This result cleanly separates code-level error-minimization, which operates on the amino acid assignment structure, from mutation-level algebraic predictions, which would require DNA polymerase errors to respect the binary encoding, which is a biologically implausible mechanism. The Fano-line construction was a meaningful test of whether the binary representation has predictive power at the **mutation** level (in addition to the **assignment** level where it does carry signal); the answer is no, and we record this falsification explicitly in Table S1.

### S14 Serine minimum inter-family distance across encodings

The claim that Serine's minimum inter-family Hamming distance (UCN–AGY) equals 4 under all 24 base-to-bit encodings is false. Of the 24 encodings, 16 yield minimum distance 2 and only 8 yield distance 4. The distance-4 result obtains only when both nucleotide pairs distinguishing UCN from AGY ($U \sim A$ and $C \sim G$ in the first two positions) are encoded at maximal Hamming distance. The correct encoding-invariant statement is: Serine is disconnected at $\varepsilon = 1$ under every encoding, and its inter-family distance ($\geq 2$) is the largest among the three 6-codon amino acids (Leucine and Arginine both have inter-family distance 1). The rejection of the distance-4 invariant is the reason the manuscript now presents $H(3,4)$ rather than $Q_6$ as the primary adjacency for encoding-independent claims.

### S15 PSL(2,7) symmetry: pre-rejection by irrep dimensions

The claim that PSL(2,7) is the fundamental symmetry group of the genetic code was pre-rejected by Antoneli and Forger (2011), who showed that PSL(2,7) has no 64-dimensional irreducible representation (its irreducible representations have dimensions 1, 3, 6, 7, and 8). A group acting on the 64-codon set must therefore decompose that action into representations that sum to 64. Any PSL(2,7)-based decomposition of the codon space is a composite of the available irreps rather than a canonical single-irrep symmetry, so PSL(2,7) does not act on the codon set as a single symmetry group in the sense of the original claim. We record this pre-rejection to explicitly distinguish the surviving graph-theoretic and hypercube-coloring analyses from the falsified algebraic-symmetry claim.

#### S16 Holomorphic embedding: character-identity failure

The claim that the coordinate-wise map $\mathrm{GF}(2)^6 \to \mathbb{C}^3$ sending base pairs to fourth roots of unity is a holomorphic embedding extending a character of $\mathrm{GF}(8)^*$ is incorrect on two counts. First, the domain is a finite discrete set (64 points), not a complex manifold, so "holomorphic" is not defined in the standard analytic sense. Second, treating the map $\chi$ as a group character requires $\chi(x+x) = \chi(x)^2$; the map fails this since $i^2 = -1 \neq 1 = \chi(0)$. The map is a bijective coordinate labelling but not a character or holomorphic embedding, and cannot support the analytic properties the original claim required.

#### S17 Source-neighborhood burden: null result

The source-neighborhood burden test asks whether reassigned codons sit in worse Hamming neighborhoods with higher Grantham distance to their neighbors. Mann–Whitney comparison of Grantham-neighborhood sums for reassigned versus non-reassigned codons yields $U = 301$, $p = 0.70$. Reassignment is therefore not driven by local escape from costly source neighborhoods. This null does not conflict with the conditional-logit finding that natural events favor candidate moves with lower $\Delta_{\text{local}}$ ($\beta_{\text{phys}} = -0.004$ in the main-text M3 fit): the two tests probe different quantities. $\Delta_{\text{local}}$ is the change in mismatch cost induced by a candidate move (a destination-quality measure); the source-neighborhood burden test asks about the absolute pre-move source-neighborhood cost. Variant codes need not originate from unusually costly source neighborhoods (no absolute source burden), yet still favor candidate moves that lower local mismatch when a reassignment is triggered (opportunistic destination selection). This indicates that the topology-avoidance constraint operates at the global graph-connectivity level, not at the local per-codon level.

#### S18 Cross-family multiple-comparison correction

Multiplicity control was applied **within** analysis families (metric family, $\rho$-sweep, topology-avoidance $2 \times 2$ audit, per-table BH–FDR, Ostrov segment tests) rather than as one cross-family correction. We did not file an external pre-registration, and the eight analysis families use non-exchangeable test statistics: hypergeometric tests on landscape counts, quartet-pattern-shuffle Monte Carlo permutation $p$-values, likelihood-ratio tests on nested conditional-logit models, BH-adjusted Fisher combinations across independent 2×2 tables, and the Bron–Kerbosch MIS Stouffer combination. These are not commensurable as one $p$-value list, so a formal cross-family Bonferroni or BH correction is not defined; ΔAICc is a model-selection quantity, not a $p$-value, and does not enter any $\alpha$-based comparison.

For descriptive transparency only, we report the primary or smallest $p$-value from each family alongside a common conservative reference threshold $\alpha^* = \frac{0.05}{8} = 6.25 \times 10^{-3}$ (Bonferroni for eight families, treated as a descriptive floor, not a formal correction). Family-primary $p$-values below $\alpha^*$: the $H(3,4)$ topology depletion ($p = 1.28 \times 10^{-6}$); the cross-metric coloring optimality (Grantham $p = 0.0062$, others $\leq 0.003$); the $\rho$-sweep ($p \leq 0.0062$ at every $\rho$); the M1→M3 conditional-logit comparison (likelihood-ratio $\chi^2 = 110.4$ on 1 df, $p \ll 10^{-20}$; the accompanying $\Delta$AICc of 108 is reported alongside as a descriptive model-selection statistic); and the per-table BH–FDR result (11 of 12 informative tables significant). Family-primary $p$-values **above** $\alpha^*$: the MIS median tRNA enrichment ($p = 0.037$; exceeds $\alpha^*$ by a factor of $\approx 5.9$) and the worst-case MIS ($p = 0.104$). Consistent with these outcomes, the tRNA enrichment is classified as **exploratory** rather than confirmatory.

Within-family corrections (applied formally): four-metric family $\alpha = 0.0125$, $\rho$-sweep family $\alpha = 0.01$, topology-avoidance $2 \times 2$ family $\alpha = 0.0125$, Ostrov segment tests $\alpha = 0.0167$; the per-table tests use BH–FDR (26 of 27 significant). Exploratory analyses are labeled as such throughout the manuscript.

### S19 Event-level conditional logit model: full details

This section gives the implementation-level detail for the discrete-choice analysis reported in main-text §3.5: the candidate universe, the three feature definitions, the optimisation routine, the order-averaging procedure for unknown event sequences, the fitted raw and normalised coefficients, the likelihood-ratio tests, the predictor-confounding diagnostic, and the per-event observed-move ranks. The complete fitted JSON output is also released as `output/evolutionary_simulation.json` in the public repository.

#### S19.1 Model specification and candidate universe

The conditional logit model treats each observed reassignment as a discrete choice from the set of all single-codon reassignments available at the current code state. For a code with 64 codons and 21 possible amino acid/stop labels, the candidate set $\mathcal{N}(C)$ contains exactly 1,280 moves (universe U1 in §S5: 64 codons $\times$ 20 alternative labels, excluding identity assignments). All 1,280 candidates are evaluated at each step regardless of biological plausibility; the model's purpose is to test whether the observed moves are statistically distinguishable from uniform sampling given the three feature classes. As a conditional logit, the model implies an independence of irrelevant alternatives (IIA) structure within each choice set. We use the model for explanatory comparison (§S6), not as an unconditional endorsement of IIA: IIA remains an untested structural assumption, the restricted-candidate refits (§S6.1) are sensitivity analyses against candidate-set composition rather than evidence that IIA is harmless, and coefficients and likelihoods could shift under a mixed-logit relaxation.

#### S19.2 Feature definitions

**Local physicochemical mismatch change ($\boldsymbol{\Delta}_{\textbf{phys}}$)**: For a reassignment of codon $c$ from amino acid $a$ to $a'$, this is the change in the sum of Grantham (1974) distances across all Hamming-1 edges incident to $c$:

$$\Delta_{\text{phys}} = \sum_{\{c,c'\}:d(c,c')=1} [\Delta(a', \text{code}(c')) - \Delta(a, \text{code}(c'))]$$

This captures the local impact on error-minimization at the reassigned position. When a neighboring codon is assigned Stop, we set $\Delta(a, \text{Stop})$ equal to the maximum Grantham distance (215), consistent with the stop-penalty convention used in the coloring objective (Methods §2.2 of the main manuscript).

**Topology disruption ($\boldsymbol{\Delta}_{\textbf{topo,Q\_6}}$)**: The total increase in connected components (at $\varepsilon = 1$) summed across all amino acid codon graphs under $Q_6$ adjacency. Stop codons are excluded from the topology sum; $\Delta_{\text{topo}}$ counts connected-component changes only for amino acid codon families. A move that splits one amino acid's codon family into two components contributes $+1$; a move that fragments two families contributes $+2$; topology-preserving moves contribute 0. The default $\Delta_{\text{topo,Q\_6}}$ uses $Q_6$ adjacency (Hamming-1 in the default $\text{GF}(2)^6$ encoding $C \to 00$, $U \to 01$, $A \to 10$, $G \to 11$). Because $Q_6$ adjacency is encoding-dependent (8 of 24 base-to-bit bijections give no $Q_6$ topology depletion at the candidate-landscape level; Section 4), we also compute $\Delta_{\text{topo,H(3,4)}}$ using the encoding-independent $H(3,4)$ adjacency (two codons are neighbors iff they differ at exactly one nucleotide position) and refit two model variants: M2$H(3,4)$ (topology-only with $\Delta_{\text{topo,H(3,4)}}$) and M3$H(3,4)$ (physicochemistry + $\Delta_{\text{topo,H(3,4)}}$). The encoding-robustness comparison $\Delta$AICc(M1 $\to$ M3$H(3,4)$) vs $\Delta$AICc(M1 $\to$ M3) is reported in the next subsection.

**tRNA complexity proxy ($\boldsymbol{\Delta}_{\textbf{tRNA}}$)**: The Hamming distance from the reassigned codon to the nearest codon already encoding the target amino acid in the current code. This serves as a heuristic for the tRNA repertoire change required to service the reassigned codon, with larger distances implying more novel tRNA machinery needed.

#### S19.3 Fitted coefficients

Table S16 reports the maximum-likelihood coefficient estimates for all six model variants (M1, M2, M3, M4 under $Q_6$ topology, plus M2$H(3,4)$ and M3$H(3,4)$ under encoding-independent $H(3,4)$ topology).

| Model | Feature | $\hat{\beta}$ (raw) | $\hat{\beta}$ (normalized) |
|---|---|---|---|
| M1 | $\Delta_{\text{phys}}$ | −0.0049 | −1.33 |
| M2 ($Q_6$) | $\Delta_{\text{topo,Q\_6}}$ | −3.5838 | −1.67 |
| M3 ($Q_6$) | $\Delta_{\text{phys}}$ | −0.0043 | −1.17 |
| M3 ($Q_6$) | $\Delta_{\text{topo,Q\_6}}$ | −3.3158 | −1.54 |
| M4 | $\Delta_{\text{phys}}$ | −0.0043 | −1.17 |
| M4 | $\Delta_{\text{topo,Q\_6}}$ | −2.9855 | −1.39 |
| M4 | $\Delta_{\text{tRNA}}$ | −0.2033 | −0.21 |
| M2 ($H(3,4)$) | $\Delta_{\text{topo,H(3,4)}}$ | −3.6957 | −1.79 |
| M3 ($H(3,4)$) | $\Delta_{\text{phys}}$ | −0.0046 | −1.24 |
| M3 ($H(3,4)$) | $\Delta_{\text{topo,H(3,4)}}$ | −3.452 | −1.67 |

Table S16: Conditional logit coefficient estimates from the 27-table re-fit. Raw coefficients are on the original feature scale; normalized coefficients are on $z$-scored features (multiplying raw $\hat{\beta}$ by the global feature standard deviation). All $\hat{\beta}$ values are negative, indicating that observed reassignment histories preferentially populate moves that reduce physicochemical mismatch, avoid topology disruption, and (weakly) prefer target amino acids already serviced by nearby codons. The tRNA proxy coefficient is small and non-significant (LR $= 0.12$, $p = 0.73$). The $H(3,4)$ verification variants ($M2_{H(3,4)}$, $M3_{H(3,4)}$) replace the encoding-dependent $\Delta_{\text{topo,Q\_6}}$ feature with the encoding-independent $\Delta_{\text{topo,H(3,4)}}$. Their fitted ΔAICc values match the main-text encoding-robustness numbers.

#### S19.4 Likelihood ratio tests

Table S17 reports nested-model likelihood-ratio statistics under both $Q_6$ and $H(3,4)$ topology variants.

| Restricted | Full | LR | df | $p$ |
|---|---|---|---|---|
| M1 (phys) | M3 (phys+topo, $Q_6$) | 110.4 | 1 | $\ll 10^{-10}$ |
| M2 (topo, $Q_6$) | M3 (phys+topo, $Q_6$) | 91.2 | 1 | $\ll 10^{-10}$ |
| M3 (phys+topo, $Q_6$) | M4 (full) | 0.1 | 1 | 0.73 |
| M1 (phys) | M3$H(3,4)$ (phys+topo, $H(3,4)$) | 93.4 | 1 | $\ll 10^{-10}$ |
| M2$H(3,4)$ (topo, $H(3,4)$) | M3$H(3,4)$ (phys+topo, $H(3,4)$) | 97.3 | 1 | $\ll 10^{-10}$ |

Table S17: Likelihood-ratio tests for nested conditional logit models, including both $Q_6$ topology variants (legacy primary) and $H(3,4)$ topology verification variants. Both topology features (Q_6 added to M1, H(3,4) added to M1) and physicochemistry (added to M2 / $M2_{H(3,4)}$) provide highly significant improvements. The tRNA-complexity proxy does not improve on the phys+topo model.

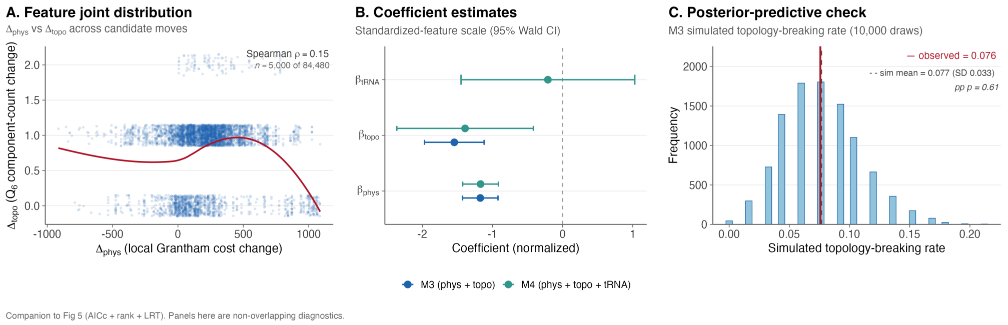


Figure S8: Conditional-logit model diagnostics. Fit quality and confounding diagnostics for the six candidate models M1–M4 + $H(3,4)$-verification variants (likelihood-ratio tests were applied only to nested pairs): joint distribution of $\Delta_{\text{phys}}$ vs $\Delta_{\text{topo}}$ across the candidate landscape (weakly correlated, $\rho = 0.15$), coefficient estimates with 95% confidence intervals, and parametric predictive replication of the observed topology-breaking rate under M3. Companion to main-text Figure 4 (which reports the AICc comparison and observed-move percentile-rank distribution).

#### S19.5 Confounding diagnostic

Across ≈84,000 candidate moves pooled from all choice sets, the Spearman correlation between $\Delta_{\text{phys}}$ and $\Delta_{\text{topo,Q\_6}}$ is $\rho = 0.15$ (Figure S8, panel A), indicating that the two predictors carry largely independent information. Moves that reduce physicochemical mismatch are only slightly more likely to also preserve topology, and the conditional logit framework accounts for any residual collinearity through simultaneous estimation. Coefficient estimates with 95% CIs (panel B) and parametric predictive replication of the observed topology-breaking rate (panel C) complete the diagnostic set.

#### S19.6 Observed move percentile ranks

Under the best model (M3), observed natural reassignments rank at the 89.5th percentile on average (mean rank 134 out of 1,280 candidates). Notable individual events: UGA→Trp ranks consistently above the 98th percentile across all tables where it occurs, indicating that this reassignment is the highest-ranked under the combined phys+topo score. The one clear outlier is the yeast mitochondrial CUU→Thr reassignment (30th percentile), consistent with NCBI translation table 3′s status as the sole marginal exception in the per-table optimality analysis.

#### S19.7 Order-averaging implementation

For tables with $k > 1$ reassignment events, the temporal ordering is unknown. We marginalized the conditional logit likelihood over all $k!$ orderings, computing $L_{\text{table}} = (1/k!) \sum_{\sigma} \prod_{s} P\left(m^*_{\sigma(s)} \mid \mathcal{N}(C_{\sigma,s})\right)$ using log-sum-exp for numerical stability. For all tables with $k \leq 6$ events (which covers every table in the dataset, including the largest, yeast mitochondrial, $k = 6$, $6! = 720$ orderings), we enumerate all orderings exactly. The implementation precomputes feature matrices once per (table, ordering, step) into stacked numpy arrays, after which each Nelder–Mead / L-BFGS-B function evaluation reduces to a small batch of matrix multiplies and `scipy.special.logsumexp` calls; the four nested models are fit concurrently on a single shared bundle via `joblib` threads to avoid memory duplication.

#### S20 ProtSub matrix and metric correlation analysis

This section evaluates whether the standard-code optimality result is confined to a single amino-acid distance scale. We quantify metric overlap across amino-acid pairs and null-code scores, and report a structure-aware ProtSub sensitivity analysis with the caveat that ProtSub is alignment-derived and therefore code-dependent in the sense of the Di Giulio (2001) critique. The analyses motivate the treatment in Methods §2.2 and Results §3.1 of the main text.

##### S20.1 ProtSub as a code-dependent robustness check

ProtSub is derived from co-evolving residue pairs in 2,320 Pfam multiple-sequence alignments of proteins encoded under the standard genetic code; it is therefore a code-dependent (MSA-derived) substitution matrix in the sense of Di Giulio (2001). We converted the EMBOSS log-odds form to a positive distance via the standard diagonal-anchored relation $d(i,j) = (s(i,i) + s(j,j))/2 - s(i,j)$, which guarantees $d(i,i) = 0$ and yields strictly positive off-diagonal entries (range 1.0 to 15.0). Sanity values: $d(\mathrm{I},\mathrm{L}) = 2.0$, $d(\mathrm{K},\mathrm{R}) = 4.0$, $d(\mathrm{F},\mathrm{Y}) = 4.0$, $d(\mathrm{D},\mathrm{E}) = 3.5$, $d(\mathrm{C},\mathrm{W}) = 14.5$.

Under the same quartet-pattern shuffle null used for the four primary metrics ($n =$10,000, seed 135325), the standard code achieves $F = 1089.5$ against null mean $1180.2 \pm 29.7$ ($z = 3.05$, quantile 0.04%, $p = 5 \times 10^{-4}$). This is the most extreme percentile of any metric in the panel. Buschmann et al. (2026) report that BLOSUM62, an AlphaFold-derived substitution matrix (AFSM), and 16 other matrix families perform similarly across multiple sequence-alignment tasks, with the interpretation that substitution matrices implicitly encode physicochemical reality; their result bounds the Di Giulio (2001) tautology rather than eliminating it. We therefore treat ProtSub as a robustness check rather than as a primary test.

##### S20.2 Metric correlation matrices

For each pair of metrics, we computed Spearman correlations at two levels: (i) across the 190 unordered amino-acid pairs of the 20 standard amino acids; and (ii) across 2,000 random codes drawn from the same quartet-pattern shuffle null used for the optimality tests. The AA-pair-level correlations bound how similar two metrics' **distance scales** are; the code-level correlations bound how similarly they **rank** codes.

| | Grantham | Miyata | PR | KD | ProtSub |
|---|---|---|---|---|---|
| Grantham | 1.00 | 0.71 | 0.47 | 0.36 | 0.63 |
| Miyata | 0.71 | 1.00 | 0.73 | 0.54 | 0.49 |
| PR | 0.47 | 0.73 | 1.00 | 0.44 | 0.26 |
| KD | 0.36 | 0.54 | 0.44 | 1.00 | 0.23 |
| ProtSub | 0.63 | 0.49 | 0.26 | 0.23 | 1.00 |

Table S18: Pairwise Spearman correlations of amino-acid distance values across the 190 unordered AA pairs (20 standard amino acids). PR = Woese polar requirement. KD = Kyte–Doolittle hydropathy. Median off-diagonal $\rho = 0.49$; minimum 0.23 (KD vs ProtSub).

| | Grantham | Miyata | PR | KD | ProtSub |
|---|---|---|---|---|---|
| Grantham | 1.00 | 0.85 | 0.75 | 0.71 | 0.83 |
| Miyata | 0.85 | 1.00 | 0.87 | 0.89 | 0.82 |
| PR | 0.75 | 0.87 | 1.00 | 0.78 | 0.74 |
| KD | 0.71 | 0.89 | 0.78 | 1.00 | 0.79 |

| | Grantham | Miyata | PR | KD | ProtSub |
|---|---|---|---|---|---|
| ProtSub | 0.83 | 0.82 | 0.74 | 0.79 | 1.00 |

Table S19: Pairwise Spearman correlations of $F$-scores across 2,000 random codes drawn from the quartet-pattern shuffle null. Higher than the AA-pair-level correlations (Table S18), as expected when distances are summed over the 192 $Q_6$ edges. Median off-diagonal $\rho = 0.82$.

| Metric pair | Partial Spearman $\rho$ |
|---|---|
| Grantham $\sim$ Miyata | +0.48 |
| Grantham $\sim$ polar requirement | +0.03 |
| Grantham $\sim$ Kyte–Doolittle | −0.31 |
| Grantham $\sim$ ProtSub | +0.54 |
| Miyata $\sim$ polar requirement | +0.50 |
| Miyata $\sim$ Kyte–Doolittle | +0.62 |
| Miyata $\sim$ ProtSub | −0.06 |
| Polar requirement $\sim$ Kyte–Doolittle | +0.01 |
| Polar requirement $\sim$ ProtSub | +0.08 |
| Kyte–Doolittle $\sim$ ProtSub | +0.39 |

Table S20: Partial Spearman correlations of code-level $F$-scores (2,000 null codes), with each pair's relationship controlled for all three other metrics simultaneously. Four of ten pairs become approximately uncorrelated after adjustment ($|\rho_{\text{partial}}| < 0.1$; no formal confidence intervals or $p$-values are reported here, and "approximately uncorrelated" is a descriptive statement about the point estimate rather than a test of statistical independence), consistent with each metric contributing unique variance not fully explained by the other four. The high pairwise correlations in Table S19 reflect a shared physicochemical signal; the partial correlations isolate the residual unique contribution of each metric.

The picture from Table S18, Table S19, and Table S20 is consistent with our framing: the metrics share substantial common variance (median pairwise $\rho \approx 0.82$ at the code level), but each contributes unique signal not captured by any combination of the others. The five-metric convergence is convergent operationalisation, not redundant replication.

### S21 Walsh–Hadamard / 2-adic spectral probe

This section establishes a finite, computable bridge between our discrete $\mathrm{GF}(2)^6$ hypercube framework and the 2-adic codon algebra developed in the BioSystems literature by Khrennikov and Kozyrev (2007), Dragovich and Mišić (2019), and Yurova Axelsson and Khrennikov (2024). Reproducible via `codon-topo` (module `src/codon_topo/analysis/walsh_2adic.py`; tests `tests/test_walsh_2adic.py`, 17/17 passing).

For each amino-acid block we form the 0/1 indicator function on the 64-codon space (indexed under the default $C \to 00$, $U \to 01$, $A \to 10$, $G \to 11$ encoding), compute its Walsh–Hadamard transform (the Fourier transform on the group $\mathrm{GF}(2)^6$), take the 2-adic valuation $v_2$ of each integer Walsh coefficient, and sum across the spectrum and across all blocks. We call this aggregate the **Walsh spectral depth** of a code.

#### S21.1 Block-size null

Under a block-size-matched null ($n$ =2,000 random partitions with the same multiset of block sizes as the standard code), the standard code's spectral depth is anomalously low (Table S21). The depth value is **mathematically invariant** across all 24 base-to-bit bijections (the statistic depends only on the underlying partition geometry, not on the encoding).

| Null model | Observed | Null mean ± SD | $z$ | Frac. ≤ obs. |
|---|---|---|---|---|
| Block-size matched ($n$ =2,000) | 544 | 689.3 ± 8.2 | −17.74 | 0.0000 |
| Wobble-box-preserving label permutation ($n$ = 2,000) | 544 | 544.0 ± 0.0 | n/a (invariant) | 1.0000 |
| Encoding sweep (24 encodings × 1,500 nulls each) | 544 | n/a per encoding | [−18.9, −16.9] | 0.0000 |

Table S21: Walsh spectral depth of the standard code under three null models. The block-size matched null gives strong shallowness ($z = -17.74$), but the stricter wobble-box-preserving label permutation reveals that the depth is mathematically constant under this null: an algebraic invariant of the (wobble box × AA slot multiset) structure. The Walsh signature of the standard code is fully encoded in the hierarchical wobble-box decomposition of synonymous codons.

#### S21.2 Wobble-box-preserving label-permutation null

The block-size-matched null does not control for the first-two-base wobble box structure that synonymous codons share. We additionally constructed a stricter null that fixes the 16 first-two-base boxes and each box's internal partition shape, randomising only the AA labels assigned to box-slots and preserving each AA's multiset of slot sizes from the standard code. Under this null the spectral depth statistic is mathematically constant at 544 (Table S21, row 2). This is consistent with the cautionary remark anticipated in the module's docstring (`WOBBLE_CAVEAT`): the spectral-depth statistic is invariant under several "natural" randomisations of the wobble structure, and the present null is one of them.

We do **not** interpret this as evidence of beyond-wobble optimisation. The honest reading is that the standard code's 2-adic Walsh signature is an algebraic invariant of the wobble-box decomposition, which is itself the hierarchical structure that Khrennikov and Kozyrev (2007) and follow-up work in the BioSystems literature model continuously on $\mathbb{Z}_2$. The Walsh–Hadamard / 2-adic probe is therefore a methodological bridge between the discrete (hypercube) and continuous (p-adic) descriptions of codon space, not a separate optimality test.

#### S21.3 Encoding invariance

Across all 24 base-to-bit bijections, the standard code's spectral depth equals 544 (mathematically invariant). The $z$-score against the block-size matched null also remains in the range $[-18.9, -16.9]$, mean $-17.9$, across all encodings (each evaluated with its own independent null draw from a deterministic seed). Figure S9 visualises both the block-size null density (panel A) and the encoding-invariance sweep (panel B).

#### S21.4 A second Walsh invariant: the wobble-free label spectrum

The block-indicator spectral depth (§S21.1–§S21.2) characterises the **geometry** of the synonymous partition. A complementary Walsh statistic characterises the **amino-acid labeling** attached to that geometry. For each sense amino acid $a$ let $f_a \in \{0,1\}^{64}$ be its indicator vector and $F_a$ its Walsh–Hadamard transform;

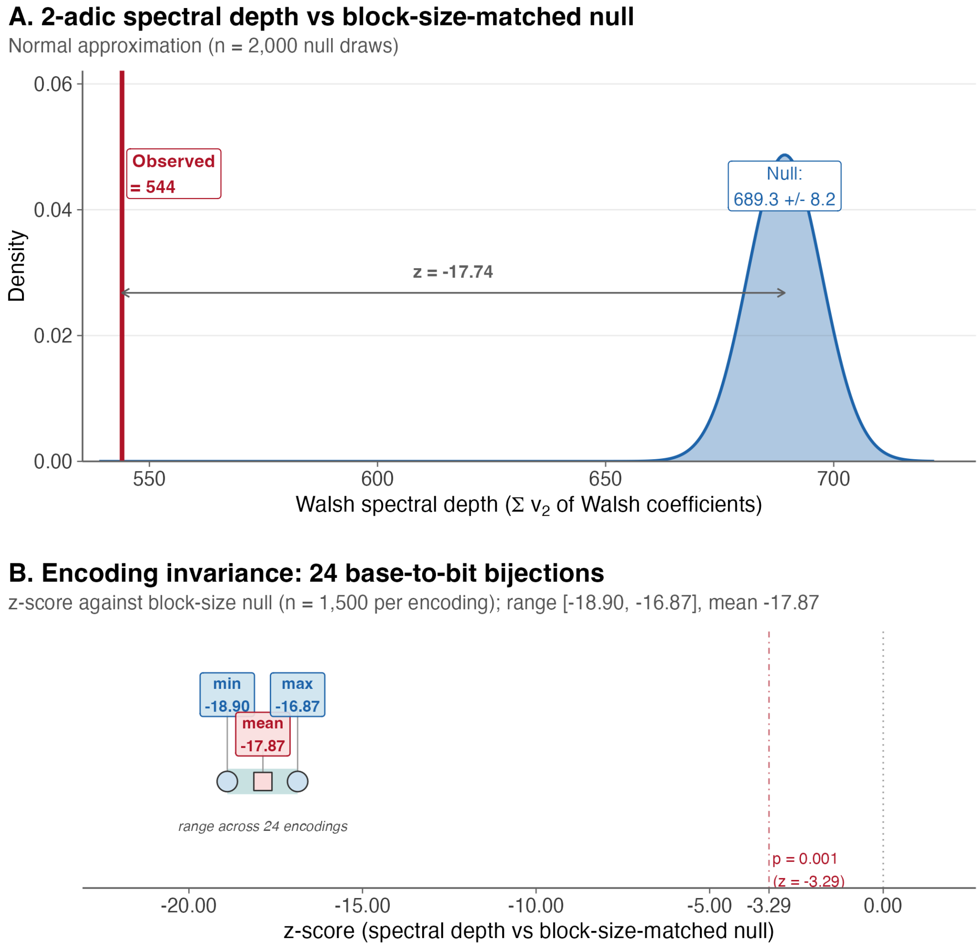


Figure S9: 2-adic Walsh spectral-depth signature of the standard code, providing a descriptive bridge to the Khrennikov/Dragovich/Axelsson 2-adic codon framework. **Panel A:** block-size-matched null distribution of the Walsh spectral depth ($n$ =2,000 random partitions with the same multiset of block sizes as the standard code). Because per-draw null samples are not persisted to `output/walsh_2adic.json`, the null is shown as a normal approximation with mean $= 689.3$ and SD $= 8.2$; the standard code's observed depth $=$ 544 (red) sits $z = -17.74$ standard deviations below the null mean (empirical fraction of null $\leq$ observed $= 0$). **Panel B:** encoding-invariance sweep across all 24 base-to-bit bijections. Each encoding is evaluated against its own independent block-size null ($n$ =1,500 per encoding, deterministic seed); z-scores fall in the tight range $[-18.90, -16.87]$ (mean $-17.87$), all $\approx 5$ times the magnitude of the two-sided $p = 0.001$ threshold ($|z| = 3.29$, dashed red line; note that the $z$-magnitude ratio is not a significance comparison and is quoted only as a descriptive scale). The observed spectral depth itself equals 544 under every encoding (mathematically invariant; see §S21.3). Together the two panels show that the standard code's Walsh signature is anomalously shallow versus a block-size-matched null and is a property of the code, not of any particular base-to-bit bijection.

partition the 63 non-DC Walsh frequencies $y \in [1, 64)$ into a **wobble-free** layer (frequencies with zero support on the two position-3 wobble bits; 15 frequencies) and a **wobble-active** layer (the remaining 48). The label-spectrum fraction is

$$S = \frac{\sum_{a \in \mathcal{A} \setminus \{\text{Stop}\}} \sum_{y \text{ wobble-free}, y \neq 0} |F_a(y)|^2}{\sum_{a \in \mathcal{A} \setminus \{\text{Stop}\}} \sum_{y \neq 0} |F_a(y)|^2}$$

For the standard code, $S = 0.7514309076$, mathematically invariant across all 24 base-to-bit bijections (`output/walsh_label_spectrum.json`).

**Two nulls, two distinct findings.** Under the **free** label-permutation null preserving only the amino-acid count multiset (stops fixed; $n$ =10,000, seed 135325): null mean $= 0.2383$, sd $= 0.0169$, range $[0.2020, 0.3328]$, $z = +30.33$, 0 of 10,000 draws reach the observed value (empirical $p < \frac{1}{n+1} \approx 10^{-4}$). Under the stricter **wobble-box-preserving** label-permutation null (the same null used for `spectral_depth` in §S21.2: fixes the 16 first-two-base boxes and each box's internal partition shape, randomises only which amino-acid label is assigned to each box-slot, preserves each AA's slot multiset): $S = 0.7514309076$ in every draw – mathematically invariant.

**Honest interpretation.** The free-permutation null shatters the wobble-box structure of 4-fold-degenerate amino acids, scattering their Walsh energy into wobble-active frequencies; the box-preserving null does not. The extreme $z = +30$ against the free null therefore reflects **exactly** the wobble-box alignment property of the standard code, not a separate optimisation layer. We do not claim "beyond wobble" optimisation. Rather, $S$ provides an exact algebraic signature of the wobble phenomenon in the Walsh basis: biological wobble degeneracy maps directly to spectral concentration on the wobble-free layer. The free-null calibration measures spectral concentration against a structurally naive baseline; it is not a test of selection beyond wobble.

**Coset quantisation of the null distribution.** The free label-permutation null is highly quantised: in 10,000 draws it produced only 15 distinct values of $S$. This reflects the coarse interaction between the AA degeneracy classes (1, 2, 3, 4, 6) and the wobble-coset partition of the Walsh frequency domain: $S$ depends only on how those classes intersect the wobble cosets, not on the within-class label assignment. The observed $S = 0.7514$ lay above all sampled null values, so the empirical upper-tail probability is bounded by the Monte Carlo resolution ($p \leq 1/(n+1) \approx 10^{-4}$).

**Mechanistic decomposition.** Each 4-fold-degenerate amino acid that occupies a complete wobble box (Pro, Ala, Thr, Val, Gly, plus the CUx, CGx, UCx cores of Leu, Arg, Ser) contributes $\approx 100\%$ of its Walsh energy to the wobble-free layer: its indicator function is constant on a wobble coset, so its Walsh transform is supported exclusively on frequencies with zero support on the two position-3 wobble bits. Singleton amino acids (Met, Trp) contribute the baseline $15/63 \approx 0.238$; a single $\delta$-function has uniformly distributed Walsh energy. The intermediate $(2, 2)$-split and $(4, 2)$-split blocks contribute mixed weights. The aggregate $S = 0.7514$ is therefore in effect a "weighted box-faithfulness of the labeling."

**Final framing for the bridge to the 2-adic literature.** The standard genetic code has two encoding-invariant Walsh signatures of wobble structure: spectral depth $= 544$ captures the synonymous-block geometry (§S21.1), and $S = 0.7514$ captures the amino-acid labeling's concentration on wobble-free frequency layers. Both support the descriptive bridge to 2-adic descriptions of the genetic code (Khrennikov and Kozyrev, 2007, Yurova Axelsson and Khrennikov, 2024) neither is evidence of selection beyond wobble.

### S22 Slavov (Tsour et al. 2026) SAAP cross-analysis

Tsour et al. (2026) published an LC-MS proteomic survey identifying 8,746 unique amino-acid substitutions across 1,767 genes in human and mouse tissues arising from alternate RNA decoding. Their high-confidence subset (Supplementary Data 8 of Tsour et al. (2026) positional probability $> 0.9$) contains 5,873 events spanning 178 unique amino-acid substitution types. We tested whether the observed substitution events are enriched for low-codon-Hamming-distance pairs, as predicted by the codon–anticodon mismatch mechanism that underlies our $GF(2)^6$ adjacency framework.

For each event we computed the minimum nucleotide-Hamming distance between any source codon (in the standard code) and any target codon. Filtering to unambiguous 20-AA pairs left 5,611 events and 166 unique observed substitution types. Table S22 tabulates the distance-1/2/3 event counts and baseline fractions; Figure S10 plots the same distribution as grouped bars for visual comparison.

| Min codon NT distance | Events | Event % | Baseline (380 pairs) % |
|---|---|---|---|
| 1 | 3,649 | 65.0% | 39.5% |
| 2 | 1,062 | 18.9% | 53.2% |
| 3 | 900 | 16.0% | 7.4% |

Table S22: Distribution of minimum nucleotide-Hamming distance between source and target codons for the 5,611 high-confidence alternate-translation events of Tsour et al. (2026) (Supplementary Data 8). Single-nucleotide-distance substitutions are dramatically over-represented relative to the all-380-pair baseline (binomial test, $p < 10^{-100}$). This is the empirical signature of codon–anticodon mismatch decoding (Hamming-1) on top of lower-rate mechanisms (RNA modifications, near-cognate decoding) that produce higher-distance substitutions.

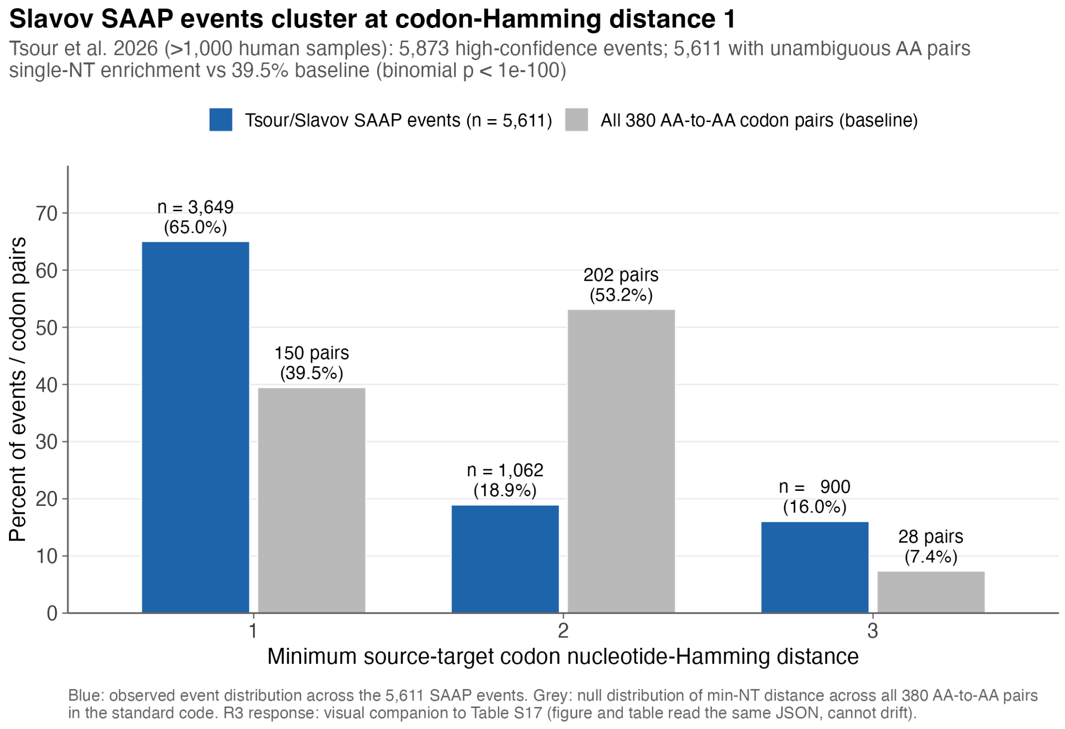


Figure S10: Visual companion to Table S22. Grouped bars compare the observed nucleotide-Hamming-distance distribution of Tsour et al.'s 5,611 high-confidence sense-codon recoding events (blue) against the null distribution across all 380 amino-acid-to-amino-acid pairs in the standard code (grey). At distance 1 the observed events are 65.0% vs a 39.5% baseline (frequency contrast is the primary descriptor; the accompanying event-weighted binomial $p \ll 10^{-100}$ ignores clustering by substitution type, gene, sample, and measurement opportunity and is reported here as a descriptive naive summary rather than a calibrated hypothesis test; (Tsour et al., 2026)). At the level of substitution **types** (unique amino-acid pairs) the enrichment is modest (41.6% at min-NT-1 vs 39.5% baseline; Fisher exact OR = 1.17, $p = 0.26$), so the effect is on event **frequency**, not type coverage. At distance 2 the observed events are **depleted** (18.9% vs 53.2%); at distance 3 they are moderately enriched (16.0% vs 7.4%). The Hamming-1 concentration is the empirical signature of codon–anticodon mismatch decoding operating on top of lower-rate mechanisms (RNA modifications, near-cognate decoding) that produce higher-distance substitutions. Figure and table are rendered from the same `output/slavov_saap_codon_distances.json` (event and baseline distance distributions).

At the level of substitution **types** (unique amino-acid pairs rather than events), the enrichment is modest (41.6% of observed types are at min-NT-1 vs 39.5% baseline; Fisher exact OR = 1.17, $p = 0.26$; Mann–Whitney U on NT distances, observed vs unobserved AAS types: $p = 0.18$). This indicates that essentially every amino-acid substitution type can occur in the Tsour et al. dataset: it is the **frequency** of each type that is biased toward single-nucleotide-distance pairs. The observation matches the mechanistic inventory reported by Tsour et al. (2026): codon–anticodon mismatch is the dominant high-frequency mechanism, with RNA modifications and near-cognate decoding contributing lower-frequency events at higher codon-Hamming distance. The Hamming-1 enrichment is independent of which amino-acid distance metric is used and is an external empirical test of the codon-adjacency assumption underlying the main-text edge-mismatch objective (equation 1).

#### S23 Exploratory observations: full detail

The main text (§3.7) summarises three exploratory observations without inferential weight; this section records the underlying detail for each.

##### S23.1 Bit-position bias in codon reassignments

The distribution of bit-flips across the 6 coordinates of $\mathrm{GF}(2)^6$ in natural codon reassignments shows apparent positional skew under a uniform null ($\chi^2 = 16.26$, $p = 0.006$, $\mathrm{df} = 5$). However, this signal is substantially attenuated after de-duplication to 20 unique (codon, target amino acid) pairs ($p = 0.075$) and vanishes entirely under a codon-preserving permutation null ($p = 1.0$). The apparent bias is therefore explained by which codons are recurrently reassigned across lineages, not by a genuine positional preference in $\mathrm{GF}(2)^6$. This is an example of a signal that survives one null model (uniform) but fails another (codon-preserving), and is retained in the record only to document the diagnostic sequence.

##### S23.2 Variant-code disconnection catalogue

A systematic survey across all 27 NCBI translation tables identifies four lineage-collapsed variant-code amino-acid disconnections at $\varepsilon = 1$ in $\mathrm{GF}(2)^6$ under the default encoding: threonine in the yeast mitochondrial code (translation table 3, CUN→Thr); leucine in the chlorophycean mitochondrial codes (translation tables 16 and 22, both with UAG→Leu; table 16 is the chlorophycean mitochondrial code (Hayashi-Ishimaru et al., 1996) and table 22 is the closely related *Scenedesmus obliquus* mitochondrial code, which additionally reassigns UCA Ser→Stop, and both produce equivalent $\varepsilon = 2$ Leu reconnection profiles so they collapse to a single algal-mitochondrial event); alanine in *Pachysolen tannophilus* nuclear code (translation table 26, CUG→Ala); and a tripartite serine in the *Candida*-clade alternative yeast nuclear code (translation table 12, CUG→Ser; (Santos et al., 1999)). These cases, combined with the universal serine disconnection, make up the complete inventory of amino-acid graph disconnections at unit Hamming distance under the default encoding.

A separate and weaker geometric exception (specific to filtration rather than to disconnection) arises in translation table 32 (Balanophoraceae plastid; UAG→Trp). Trp is 1-fold (UGG only) under the standard code; in table 32 it becomes 2-fold (UGG, UAG). The pair differs in the second nucleotide (G↔A), i.e. at bit position 3 in our 0-based 6-bit indexing (the second bit of the second nucleotide), not at the wobble bit (position 5) where every standard-code 2-fold pair sits. Hamming distance 1, so the pair is connected at $\varepsilon = 1$ and adds no new entry to the disconnection catalogue. An empirical scan over every 2-fold amino-acid pair in all 27 NCBI tables confirms that this is the unique deviation from the bit-5 two-fold filtration: the

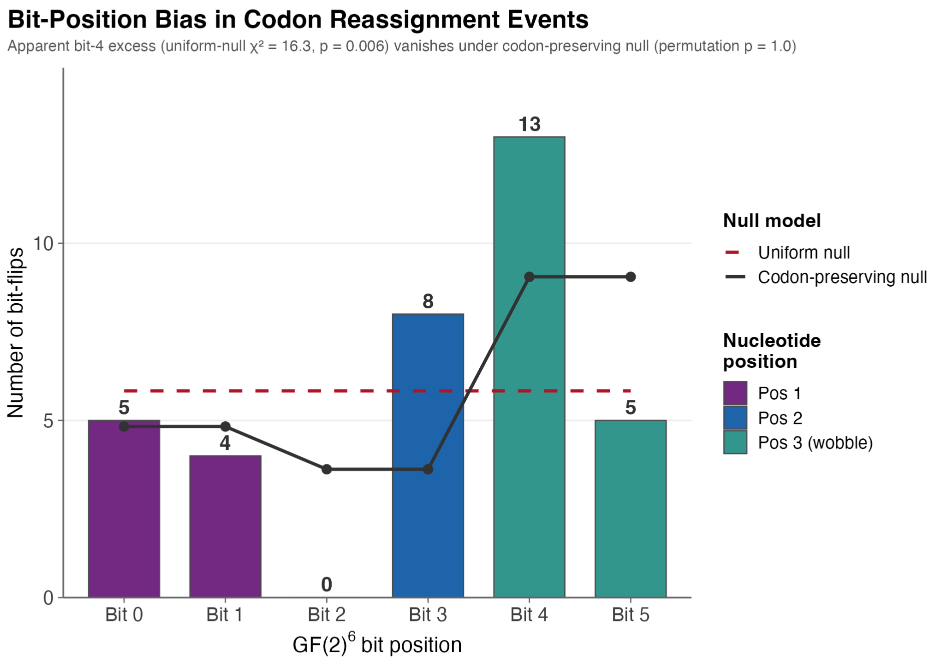


Figure S11: Bit-position bias in codon-reassignment events. Observed bit-flip counts at each of the 6 coordinates of $\mathrm{GF}(2)^6$, overlaid on the uniform-null expectation and the codon-preserving permutation null. The apparent uniform-null skew ($p = 0.006$) is fully explained by the codon-preserving null ($p = 1.0$), consistent with a recurrent-codon effect rather than a genuine positional preference. Retained as a null-model-diagnostic example.

standard code's nine 2-fold amino acids and every analogous pair introduced by the other 26 tables differ at bit 5.

#### S23.3 Atchley Factor 3 and Serine convergence

Serine has the most extreme Atchley Factor 3 score among the 20 amino acids ($F_3 = -4.760$, 2.24 SD below the mean; (Atchley et al., 2005)), and it is the only amino acid disconnected at $\varepsilon = 1$ under every base-to-bit encoding. This convergence is unsurprising: Atchley $F_3$ is a composite of molecular size and codon diversity that partly captures codon-family structure, so the $\mathrm{GF}(2)^6$ topology and $F_3$ are not fully independent views of Serine's anomaly. Both reflect Serine's disproportionate codon diversity (6 codons in two disconnected families) relative to its small physicochemical footprint; the $\mathrm{GF}(2)^6$ framework provides a complementary structural account rather than independent corroboration.

#### S23.4 Three-tier tRNA mechanistic landscape (from main-text §3.6)

The manuscript §3.6 summarises the tRNA enrichment result; the mechanistic detail moved from that section is recorded here for completeness. The largest observed case is *Tetrahymena thermophila* (NCBI translation table 6, UAA/UAG reassigned to Gln; (Hanyu et al., 1986)), which carries 54 glutamine tRNA genes (including 39 suppressor tRNAs reading the reassigned stop codons), compared to 3 Gln tRNAs in the standard-code ciliate *Ichthyophthirius multifiliis* (assembly GCF_000220395.1), 11 in *Stentor coeruleus* (assembly GCA_001970955.1), and 3 in *Fabrea salina* (assembly GCA_022984795.1 from (Zhang et al., 2022)). All four counts were generated in this work by running tRNAscan-SE 2.0.12 on the listed assemblies. The pattern extends across reassignment types. Among Gln-reassignment ciliates, *Pseudocohnilembus persalinus* (20 Gln tRNAs including 15 suppressors) and *Halteria grandinella* (9 Gln tRNAs including 3 suppressors; (Zheng et al., 2021)) represent independent lineages within Oligohymenophorea and Spirotrichea respectively. Among Cys-reassignment ciliates (NCBI translation table 10, UGA→Cys), six tRNAscan-SE–verified *Euplotes* species all carry tRNA-Cys genes with the non-canonical TCA anticodon (reading UGA), with 1–4 such genes per species alongside standard GCA-anticodon Cys tRNAs.

However, the pattern is not universal. *Blastocrithidia nonstop* (NCBI translation table 31) reassigned all three stop codons (as did *Condylostoma magnum*, where stop-codon function is context-dependent; (Heaphy et al., 2016)) but achieved UGA→Trp via anticodon stem shortening (5 bp → 4 bp) of tRNA-Trp(CCA), combined with an eRF1 Ser74Gly mutation, rather than gene duplication (Kachale et al., 2023). Similarly, *Mycoplasmoides* species with UGA→Trp use a single tRNA-Trp with anticodon modification. These boundary cases define a three-tier mechanistic landscape: (i) tRNA gene duplication in large nuclear genomes, (ii) anticodon structural modification in streamlined genomes, and (iii) anticodon base modification in minimal genomes.

#### S24 Software and reproducibility

This section provides the metadata needed to reproduce every number in the manuscript and supplement **within numerical and rendering tolerance** from the public repository. Every figure, table, and inline statistic is rendered by the Typst sources `manuscript.typ` and `supplement.typ` (also in the repository) from the JSON outputs of a single `codon-topo all` invocation, so within a single pipeline run the manuscript and supplement are generated from shared versioned artifacts. We do not claim bit-for-bit reproducibility: the pinned software floor below fixes randomness (seeded RNG) and language versions, but does not pin every transitive dependency, every optimizer default, or every renderer version, so exact byte-level identity of PDFs, PNG rasters, and floating-point outputs across systems is not guaranteed. Numerical values in the JSON artifacts, quantile ranks, and permutation-test $p$-values should reproduce to at least three significant figures; rendered PDFs may differ in font subsetting, image compression, and Typst layout across Typst versions.

All analyses were performed using the `codon-topo` Python package (version 0.6.1, tag `v0.6.1`). The code is publicly released at https://github.com/biostochastics/codontopo. Dependencies and runtime requirements:

- Python 3.11 (tested on 3.11.14), NumPy 1.24+, SciPy 1.10+
- R 4.5, ggplot2, ggpubr, viridis, patchwork (for figures)
- tRNAscan-SE 2.0.12 with Infernal 1.1.4
- Typst 0.14.2 (for manuscript and supplement PDFs)
- Random seed: 135325 (all Monte Carlo analyses)
- Exact package versions used in the reference build are listed in `requirements.lock` (Python) and `renv.lock` (R) in the release tag; a `Dockerfile` is provided for a containerized rebuild. Users who install unpinned dependencies should expect reproducibility only within numerical/rendering tolerance rather than bit-for-bit.

Analyses are fully reproducible from a clean clone at tag `v0.6.1` by one canonical command:

```
git clone https://github.com/biostochastics/codontopo && cd codontopo
git checkout v0.6.1
pip install -e ".[dev]"
bash scripts/build_publisher_release.sh
```

The wrapper `scripts/build_publisher_release.sh` runs the analysis pipeline (`codon-topo all --seed=135325 --n=10000`), the provenance-emit and table-generation scripts, both R figure scripts, and the two Typst PDF compilations, in that order. The `[dev]` extra installs the test toolchain used during development and continuous integration.

Zehir, A., Benayed, R., Shah, R.H., Syed, A., Middha, S., Kim, H.R., Srinivasan, P., Gao, J., Chakravarty, D., Devlin, S.M., Hellmann, M.D., Barron, D.A., Schram, A.M., Hameed, M., Dogan, S., Ross, D.S., Hechtman, J.F., DeLair, D.F., Yao, J., Mandelker, D.L., Cheng, D.T., Chandramohan, R., Mohanty, A.S., Ptashkin, R.N., Jayakumaran, G., Prasad, M., Syed, M.H., Rema, A.B., Liu, Z.Y., Nafa, K., Borsu, L., Sadowska, J., Casanova, J., Bacares, R., Kiecka, I.J., Razumova, A., Son, J.B., Stewart, L., Baldi, T., Mullaney, K.A., Al-Ahmadie, H., Vakiani, E., Abeshouse, A.A., Penson, A.V., Jonsson, P., Camacho, N., Chang, M.T., Won, H.H., Gross, B.E., Kundra, R., Heins, Z.J., Chen, H.-W., Phillips, S., Zhang, H., Wang, J., Ochoa, A., Wills, J., Eubank, M., Thomas, S.B., Gardos, S.M., Reales, D.N., Galle, J., Durany, R., Cambria, R., Abida, W., Cercek, A., Feldman, D.R., Gounder, M.M., Hakimi, A.A., Harding, J.J., Iyer, G., Janjigian, Y.Y., Jordan, E.J., Kelly, C.M., Lowery, M.A., Morris, L.G.T., Omuro, A.M., Raj, N., Razavi, P., Shoushtari, A.N., Shukla, N., Soumerai, T.E., Varghese, A.M., Yaeger, R., Coleman, J., Bochner, B., Riely, G.J., Saltz, L.B., Scher, H.I., Sabbatini, P.J., Robson, M.E., Klimstra, D.S., Taylor, B.S., Baselga, J., Schultz, N., Hyman, D.M., Arcila, M.E., Solit, D.B., Ladanyi, M., Berger, M.F., 2017. Mutational landscape of metastatic cancer revealed from prospective clinical sequencing of 10,000 patients. Nature Medicine 23, 703–713. https://doi.org/10.1038/nm.4333

Zhang, B., Hou, L., Qi, H., Hou, L., Zhang, T., Zhao, F., Miao, M., 2022. An extremely streamlined macronuclear genome in the free-living protozoan Fabrea salina. Molecular Biology and Evolution 39, msac62. https://doi.org/10.1093/molbev/msac062

Zheng, W., Wang, C., Lynch, M., Gao, S., 2021. The compact macronuclear genome of the ciliate Halteria grandinella: a transcriptome-like genome with 23,000 nanochromosomes. mBio 12, e1964–20. https://doi.org/10.1128/mBio.01964-20
